# Heterogeneous strains in tissue collagen show that high strains locally suppress degradation by collagenase

**DOI:** 10.1101/2021.02.07.430141

**Authors:** K. Saini, M. Tiwari, S. Cho, A. Jalil, M. Vashisth, J. Irianto, J. Andrechak, L. Dooling, C. Alvey, A. Kasznel, D. Chenoweth, K. Yamamoto, D. Discher

## Abstract

Collagen, the most abundant protein in mammals, contributes to the physical properties of different tissues during development, homeostasis, and disease. The adaptation of physical properties of tissues to mechanical stimuli is thus dependent on the control of tissue collagen levels by well-regulated synthesis and degradation of collagen. Importantly, how various molecular-level events within a tissue sustaining a range of mechanical strains contribute towards maintaining its collagen levels, remains unclear to date. Such molecular level processes in tissues are studied here in the case of isolated tendons consisting of collagen fibrils oriented along tissue loading-axis and beating embryonic hearts to gain understanding of mechanical load dependent tissue sculpting. Using a novel bioreactor design, starved mice tail tendon fascicles were used as a “cell-free” model and were subjected to heterogeneous and uniaxial deformation modes. Patterned photobleaching of fluorescent probes, a novel Aza-peptide or dye, on fascicles used to quantify tissue strains. Tissue microstructure was simultaneously imaged using second harmonic generation (SHG) signal to assess tissue collagen content while deformed fascicle samples were exposed to purified matrix metalloproteinase-1 (MMP-1) or bacterial collagenase (BC). A decrease in the degradation rate (relative to strain-free) was observed for physiological strain limits of tendon tissue (i.e. ∼5-8%) while at higher strains (i.e. pathological) the degradation rate was independent of strain magnitude changes. Interestingly, the strain dependence of degradation rate was independent of cleavage-site specificity of the collagenase molecules and the mode of tendon tissue deformation. Although spatially different within a tissue sample, the values of strain, degradation rate and collagen fiber organization with time during degradation of each tendon fascicle region were highly correlated. Tendon regions dominated by collagen fibers inclined to fascicle-axis were observed to follow non-affine deformation. The dependence of the degradation rate on mechanical strain is due to sequestration of collagen cleavage sites within fibrils. Permeation, tissue mass density and mobility of fluorescent collagenase and dextran are strain-independent for fascicle strains up to ∼5-8% while the degradation rate is positively correlated to unfolded triple-helical collagen content. Normal beating chick hearts subjected to ∼5% peak strain in a spatiotemporal coordinate contractile wave were observed to maintain their collagen mass until the beating strain is suppressed by inhibition of myosin-II. Based on the presence of exogeneous MMP inhibitors, endogenous MMPs within the non-beating hearts degrade the collagens immediately (in ∼30-60 mins). Both tissue systems under mechanical strains suggest degradative sculpting where mechanical strain-dependent collagen fibril architecture changes appear to play a key role in determining collagen lifetime within tissues.

**Graphical abstract:** 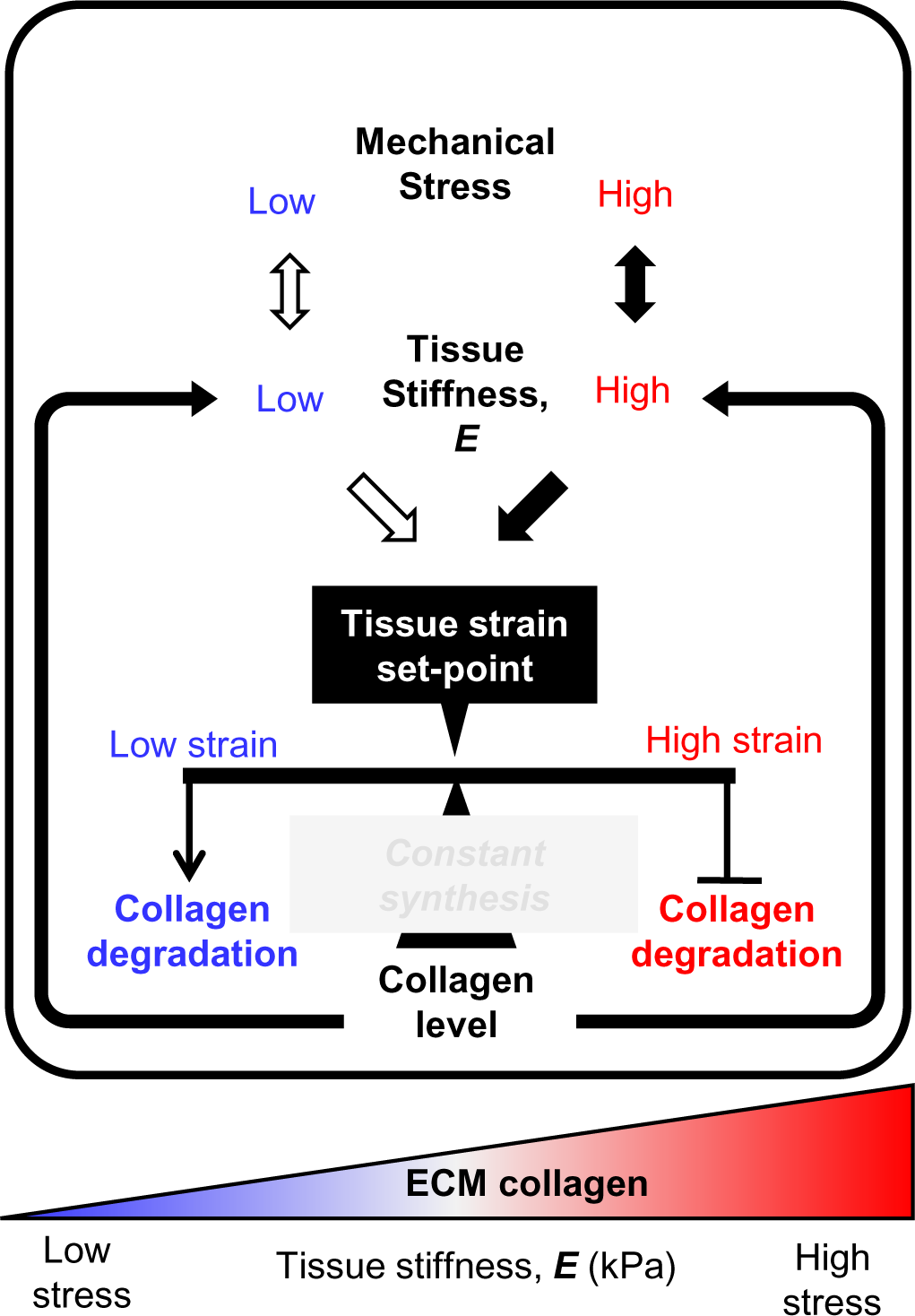

## 1. Introduction

The extracellular matrix (ECM) of tissues is composed of a variety of structural proteins that establishes its stiffness where the addition of bacterial collagenase (BC) rapidly fluidizes a tissue. Physical properties of polymer systems, for example elasticity and viscosity, generally scale with polymer concentration (Gennes 1979) which holds true for tissue’s most abundant biopolymer i.e. collagen type-I (Yang, Leone et al. 2009), and polymer interactions, assembly and crosslinking generally modulate such physical properties (Doi and Edwards 1988). Indeed, tissue stiffness (*E*) scales with the levels of collagen type-I and several other fibrillar collagens as *E ∼* [*collagen*]*^m^, with m ∼* 0.6 to 1.1. In agreement with the power-law scaling relationship, softer tissues of an adult mice such as brain and bone marrow possess less collagen than their stiffer counterparts such as muscle or bone (Swift, Ivanovska et al. 2013). Collagen accumulation has also been shown to stiffen the developing heart of chick embryos during its first few days of beating (Cho, Vashisth et al. 2019). Hence, tissue collagen level is a key determinant of the physical properties of tissues.

For processes like development, homeostasis and disease in tissues, mechanoregulation of stable gene expression levels for structural proteins such as nuclear lamin A, mini-filaments of myosin-II, and collagen fibrils, obey the concept of “*use it or loose it*” through stress inhibited degradation (Dingal and Discher 2014) (Buxboim, Swift et al. 2014) (Saini, Cho et al. 2019). Homeostasis is strictly regulated by a balance of degradation and synthesis of ECM proteins including various collagens in a tissue subjected to mechanical stresses. Mechanical stresses sustained and/or generated by stiffer tissues tend to be higher than those in softer tissues, consistent with an inter-relationship between: (*i*) tissue stiffness, (*ii*) the typical stress in the tissue (i.e. tissue stress ∼ *E*), and (*iii*) the homeostatic balance of collagen synthesis and degradation. The adaptation of tissue physical properties in terms of stiffness, cross-sectional area, etc., against sustained stresses, is theoretically a function of synthesis and degradation rates of proteins in addition to the magnitude and duration of the stresses (Bohm, Mersmann et al. 2014). Tissue collagens are degraded by specialized enzymes called matrix metallo-proteinases (MMPs) that putatively target collagen monomers at a specific site to produce ¾ and ¼ fragments (Chung, Dinakarpandian et al. 2004). Another related proteinase, cathepsin-K, is capable of cleaving collagen at multiple sites similar to BC (Garnero, Borel et al. 1998), and further degrades the fragments initially produced by MMPs (Garnero, Ferreras et al. 2003). In the case of tendons, the load deprivation for longer duration (days to weeks) results in decreased tissue stiffness, size and increased in ECM MMP levels (Arnoczky, Tian et al. 2004) (Sun, Thoreson et al. 2010). Physical exercise immediately suppresses the rate of collagen synthesis in the human Achilles tendon followed by a return to its basal level within 24 hours and an increase by ∼3 fold after 72 hours (Langberg, Skovgaard et al. 1999) (Dideriksen, Sindby et al. 2013). Interestingly, collagen degradation rate within human achilles tendon, measured in terms of the level of MMP-degraded collagen fragments, decreases by ∼0.5 fold immediately following exercise and reaches its basal level again after 72 hours (Langberg, Skovgaard et al. 1999). Recently, circadian rhythms has been shown to contribute a change of ∼±3% in mice tendon collagen level, and the newly synthesized collagen, produced as non-covalently cross-linked smaller diameter fibrils, was observed not to influence tissue elastic properties significantly (Chang, Garva et al. 2020). Therefore, physical loading of tendons appears to regulate tissue collagen mass by instantly influencing the collagen degradation rate, presumably independent of collagen synthesis.

Tissue ECM is composed of insoluble multiple highly-conserved, triple-helical collagens (types I, II, III, V and XI) in the form of fibrils (Kadler, Hill et al. 2008) (Gelse, Poschl et al. 2003) (Holmes, Lu et al. 2018) (Kadler 2017) besides other non-collagenous proteins whose interactions with a soluble phase might change during the course of mechanical deformation of a tissue. In this light, it becomes fundamentally important to understand how mechanical strain in tissues influences collagen degradation by the action of different collagenases, tissue structure and enzyme transport. The explicit relationship among tissue strain, collagen degradation and structural changes in physiological environments and associated molecular mechanisms remains unclear despite reports on the mechanosenstive nature of fibrillar collagen (Huang and Yannas 1977) (Wyatt, Bourne et al. 2009) (Flynn, Bhole et al. 2010). Hence, we aimed (a) to develop a relationship between mechanical strain and collagen degradation rate in “cell free” tissues exposed to different collagenases, (b) to determine the influence of mechanical deformation mode on strain-dependent tissue collagen degradation, and changes in tissue strain-dependent collagen fiber organization during collagen degradation, (c) to assess tissue strain-dependent changes in local tissue mass density, solute permeation and mobility, (d) to measure the dependence of tissue collagen levels on mechanical strain ex-vivo, and (e) to determine the contribution of collagen molecular structural alterations (via crosslinking using exogenous enzyme) on strain-dependent tissue collagen degradation. We developed a novel bioreactor simultaneously facilitating microscale fascicle tissue deformation and second harmonic generation (SHG) imaging in both forward and backward directions. In order to fluorescently label the tissue samples for pattern-photobleaching without influencing collagen degradation rate, we used a novel Aza-peptide in addition to using commercially available probes. First, we selected starved mice tail tendon fascicles, a cell-free model that excludes the influence of collagen synthesis, under heterogeneous strain for exposure to exogeneous collagenases (i.e. purified MMP-1 and BC) to develop a relationship between tissue mechanical strain and collagen degradation. SHG imaging, a label-free collagen fibrillar structure sensitive method, was employed to quantify fibrillar collagen content in the tissue (Williams, Zipfel et al. 2005) (Lacomb, Nadiarnykh et al. 2008). Secondly, in addition to three-point bending deformation mode causing heterogenous strain distribution, tendon fascicles were deformed uniaxially along the fascicle-axis to predict strain-dependent collagen degradation and associated collagen fiber orientation changes. Thirdly, in the absence of collagen degradation, tendon fascicles were investigated for mechanical strain induced local changes in the organization of collagen fibers, permeation and mobility of the molecules, and mass density of the tissue. Fourthly, the development of the chick heart to embryonic day 4 (E4) parallels the expression of collagen type-I protein in addition to other excitation-contraction proteins (Majkut, Idema et al. 2013). We elucidated strain-dependent changes in collagen levels in beating embryonic chick hearts by pharmocologically inhibiting tissue contractility ex vivo. Finally, collagen was artificially crosslinked using exogeneous enzyme i.e. transglutaminase (TGM) to reveal the relationship between microscale tissue strain, collagen degradation and deformation-induced, molecular-level changes in the fascicle tissue collagen.

## 2. Materials and Methods

### 2.1. Materials

#### 2.1.1. Mice tail tendon fascicles

In accordance with institutional animal care and use procedures, tendon fascicles from the tails of wild-type (WT) mice (C57BL/6J, 9-12 weeks), euthanized for other unrelated studies, were used in this investigation. Following euthanasia and using a sterile scalpel blade, the tail was sectioned between the coccygeal vertebrae at the base and distal tip of the tail, resulting a total length of up to ∼70 mm. The tail was immediately washed and placed in PBS (Corning 21-031-CV) at room temperature for immediate extraction of fascicles or at −20°C for long term storage. Fascicles stored at - 20°C were thawed at 4°C (for few hours) and at room temperature (∼60-90 minutes) before fascicle extraction. For determination of spatial distribution of cells within a live tail tendon fascicle, NSG mice (9-12 weeks) were selected in addition to WT mice. For samples used in heterogeneous deformation, tail was sectioned into at least ∼10-15 mm sub-length specimens while for uniaxial deformation testing, tail samples at least ∼50 mm long were considered. Fascicles (of diameter up to ∼200 *μ*m) were teased out of the tail samples within ∼2 hours while maintaining tissue hydration using PBS. Each group of 3-4 fascicles were kept in small Eppendorf tubes (in 1.5 ml of PBS) for storage at −20°C. Before start of each experiment, each fascicle sample was thawed at 4 °C overnight followed by washing few times in PBS at room temperature. For all enzymatic reactions involving BC i.e. Clostridium histolyticum (Sigma-Aldrich, C7657), PBS with Ca/Mg (Thermo. Scientific, 14040117) was used as reaction buffer. For all studies involving purified and activated MMP-1, 50mM Tris-HCl/ 150 mM NaCl/ 10mM CaCl_2_ with 0.01%, bovine serum albumin (BSA) solution were freshly prepared for use as reaction buffer (Suzuki, Enghild et al. 1990) (Chung, Dinakarpandian et al. 2004). Fluorescent labelling of fascicles was done using Col-F dye (Immunochemistry Tech., cat. no. 6346) or aza-peptide [5(6)-CTAMRA-NH(PEG)_3_CO-(azGPO)_3_-NH_2_] with M.W. as 1481.58. The solutions of 4.4 kDa, 155 kDa or 500 kDa dextrans (Sigma Aldrich, T-1037, T-1287, 52194) in PBS (0.1 mg/mL) were used for FRAP studies. Fluorescent-BC (0.01 mg/mL) in PBS was prepared (see supp. materials, Fig. S19) by labelling Clostridium histolyticum (Sigma-Aldrich, C7657) at N-terminus using Alexa Fluor™ 555 NHS Ester (Invitrogen™: A20009) as per manufacturer protocol. For artificial cross-linking, fascicle samples were incubated in ∼1-5 mg/mL Transglutaminase (TGM) (Sigma-Aldrich, T5398) solution in DMEM (Thermo. Scientific, 21063029) having 10% FBS + 1% P/S. For detection of unfolded collagen within samples, Biotin conjugate of collagen hybridizing peptide (CHP), (3Helix, Inc., B-CHP) and fluorescent Streptavidin conjugate Alexa-594 (Thermofisher -S11227) were used following biotin blocking of tissues samples using endogenous Biotin-blocking kit (Life Technologies Corporation/Thermo fisher, E21390). For Aza-peptide preparation, the following reagents and solvents were used as received: 2-chlorotrityl chloride resin (ChemPep.), Low-loading rink amide MBHA resin (Novabiochem), Fmoc-Pro-OH, CDT, and Fmoc-NH(PEG)_3_-COOH (Chem-Impex); Fmoc-Hyp(tBu)-OH (AK Scientific); COMU CreoSalus (Advanced ChemTech); DIEA (Acrōs); TFE, TIPS, HBTU, and Fmoc-carbazate (Oakwood); Piperidine (EMD Millipore); 5(6)-CTAMRA (Carbosynth/Novabiochem); TFA (Alfa Aesar); all other solvents were acquired from Fisher or Sigma.

#### 2.1.2. Embryonic chick hearts

White Leghorn chicken eggs (Charles River Laboratories; Specific Pathogen Free (SPF) Fertilized eggs, Premium #10100326) were used to extract embryonic hearts. SPF chicken eggs from Charles River are produced using filtered-air positive-pressure (FAPP) poultry housing and careful selection of layer flocks. Every flock’s SPF status is validated in compliance with USDA memorandum 800.65 and European Pharmacopoeia 5.2.2 guidelines. SPF eggs were incubated at 37°C with 5% CO_2_ and rotated once per day until the desired developmental stage (e.g. four days for E4; Hamburger-Hamilton stage 23-24 (HH23-24)). Embryos were extracted at room temperature (RT) by windowing eggs, carefully removing extra-embryonic membranes with sterile sonicated forceps, and cutting major blood vessels to the embryonic disc tissue to free the embryo. The extracted embryo was then placed in a dish containing PBS on a 37°C-heated plate, and quickly decapitated. For early E2-E5 embryos, whole heart tubes were extracted by severing the conotruncus and sinovenosus (for methods, see (Cho, Vashisth et al. 2019)). All tissues were incubated at 37°C in pre-warmed chick heart media (α-MEM supplemented with 10 % FBS and 1% penn-strep, Gibco, #12571) for at least 1 h for stabilization, until ready for use. Bacterial collagenase (BC) i.e. Clostridium histolyticum (Sigma-Aldrich, C7657) was used as exogenous collagenase in chick heart experiments and actomyosin perturbation was done by blebbistatin (EMD Millipore, #203390) and MMP activity was inhibited by ilomastat (GM6001).

### 2.2. Methods

#### 2.2.1. Environment control mechanical manipulation bioreactor

The schematic of environment control bioreactor, installed on a MP microscope, used for the heterogeneous deformation of the tendon fascicles in the present work is shown in Fig. S2 (for detailed design, installation of the bioreactor on MP-microscope as well as placing sample within the bioreactor, see supp. info. S1). Due to smaller displacement between sample and signal detector(s), SHG signal is captured easily in both backward and forward directions for the sample (of diameters up to ∼200 *μ*m and length larger than ∼10 mm) kept in the bioreactor. The movable plate facilitates in-plane deformation of the sample via three-point bending and results its heterogeneous deformation. The design of the bioreactor facilitates immediate exposure of the sample to any solute (e.g. collagenase, dextrans, etc.) added to the reaction buffer within the petri-dish while maintained at 37°C by the bottom heating plate through a separate temperature control unit. In case of uniaxial deformation tests (shown in Fig. 2A-*i*), each fascicle sample was cut into two sub-lengths (to avoid sample to sample variation) where one sub-length was kept strain-free while other sub-length was strained along fascicle-axis whereas other experimental setting were similar to that of heterogeneous deformation. Glass coverslips were used (instead of polystyrene) as base plate in the bioreactor in order to avoid the adhesion contact friction. The samples were immersed in reaction buffer medium during throughout the experiment. The strain-free sub-length of the sample was kept at constant strain via gluing at both ends whereas to-be-strained sub-length was glued only at one end while other end was kept free for subsequent uniaxial deformation (followed by initial imaging in the undeformed configuration). It must be kept in mind that during use of bioreactor on an upright microscope, evaporation of reaction medium takes place that needs regular compensation. In order to avoid reaction buffer evaporation from edges of the bioreactor while maintaining same temperature and composition of reaction buffer, the petri dish carrying the deformed sample was covered and moved to incubator between time points of imaging only for the degradation experiments involving MMP-1 (for composition dependence of MMP-1 activity, see Fig. S8).

#### 2.2.2. Fluorescent labeling of fascicles

The schematic in Fig. S4A shows various structural hierarchy levels of a tendon tissue and highlights the fascicle level selected for the present investigation. In order to quantify tissue strain magnitudes (see section 2.2.3), the fascicles were fluorescently labelled with either Aza-peptide or Col-F dye. For aza-peptide synthesis & purification (for details, see section 2.2.13) (Kasznel 2020), HPLC was performed using Agilent 1260 Infinity II systems equipped with Phenomenex PFP(2) columns. Flash chromatography for purification of building block Fmoc-azGPO(tBu)-OH was performed using a Teledyne ISCO CombiFlash Rf chromatography system. MALDI-TOF MS was performed using a Bruker MALDI-TOF Ultraflex III mass spectrometer. UV-vis spectroscopy was performed using a JASCO V-650 UV-vis spectrophotometer. Probe aliquots were dried using an Eppendorf Vacufuge plus concentrator for storage prior to use in subsequent assays. Fmoc-azGPO(tBu)-OH was synthesized as described in supp. info. S10. For fluorescent labelling with aza-peptide, fascicles were kept in 30 *μ*M Aza-peptide [5(6)-CTAMRA-NH(PEG)_3_CO-(azGPO)_3_-NH_2_] solution in PBS. Aza-peptide solution was heated to 80°C for 5-7 minutes (to obtain its monomeric form) and quickly cooled to 4°C (by keeping on ice for ∼25 secs.) before application to samples for overnight incubation at 4°C. The low molecular weight fluorescent probe, Col-F (fluorescein conjugated to Physostigmine), exhibiting non-covalent affinity to ECM fibers (Biela, Galas et al. 2013), was used to label fascicles. Fascicles were kept for 30 minutes in the dye solution (at ∼50 *μ*M) at room temperature, followed by at least two PBS washes of 30 minutes each using tube revolver in order to remove the free dye molecules. Fluorescently labeled fascicles were protected from light and were kept at room temperature until their use within ∼2 hours (or kept on ice in case of longer waiting durations).

#### 2.2.3. Fluorescent pattern photobleaching (FPP) and strain quantification

For creating a user-defined pattern using the FPP method, a region on the fascicle was selected (positioned at ∼half diameter deep into the fascicle) that carried ten equally spaced rectangular stripes of ∼5 *μ*m width and ∼50 *μ*m spacing between them (edge to edge) as shown in Fig. S4B-*i*. In order to quantify the displacement between each pair of photobleach stripes, a projected sum composite image, from z-stack for the fascicle region carrying photobleach stripes, was created. The intensity in each photobleach stripe region followed Gaussian distribution and therefore, the Gaussian peak was used to represent each stripe position on the fascicle. Mechanical deformation of the fascicle in the bioreactor resulted in a change in the displacement between each pair of consecutive photobleach stripes as shown in Fig. S4(B-*C*). Each point on the ROI path (a representative thick yellow line in Fig. S4B-*ii*) represents the average intensity of pixels (over few pixels in transverse direction to ROI path) while ROI path direction is kept parallel to fascicle longitudinal direction (between each pair of photobleach stripes). A fixed number of points on the ROI path were selected for Gaussian curve fitting throughout data analysis in order to determine the position of photobleach stripes on the fascicle and corresponding displacement between each pair of photobleach stripes using a custom MATLAB script (see supp. info. S2). The magnitudes of displacement between each pair of consecutive photobleach stripes in undeformed and deformed fascicle (represented as *D_f_* and *D_i_* respectively) were then used to calculate the magnitude of engineering strain, *ɛ* (= (*D*_*f*_ − *D*_*i*_)/*D*_*i*_ ∗ 100) within each region of the fascicle (Fig. S4C-*ii*).

#### 2.2.4. Second Harmonic Generation (SHG) imaging

Within a tendon fascicle, collagen fibrils assemble longitudinally to form collagen fibers aligned along the fascicle-axis. SHG intensity has been observed to depend on the angular orientation of the fascicle-axis with respect to the direction of polarization of incident light [as shown in Fig. S16A] (Williams, Zipfel et al. 2005). The orientation dependent variation in SHG signal magnitude with respect to light polarization direction can be used to estimate absolute SHG intensities within a focal volume of a mechanically deformed fascicle containing collagen fibers with many orientations, (for directionality-based scaling of the measured SHG signal, see Fig. S16). Here, it is important to keep in mind that smaller diameter collagen fibrils (∼λ_SHG_/10) emit more backward SHG signal than forward SHG while larger ones emit more forward SHG signal (Lacomb, Nadiarnykh et al. 2008), suggesting SHG signal in backward and forward directions indicates the dominance of various size collagen fibrils in tissue collagen content. Upon degradation of collagen molecules present in the fascicle by the collagenase, the insoluble collagen fibrils are converted into soluble smaller-mass peptide fragments incapable of producing sufficient SHG signal, and therefoere suggesting a basis for the decrease in SHG signal magnitude during tissue collagen degradation. For experiments involving comparison of the absolute SHG intensities, either the linearly polarized fundamental was aligned parallel to the fascicle-axis or obtained values were scaled based on the angular orientation directions of collagen fibers (for more information on collagen content measurement using SHG signal, see supp. info. S3).

#### 2.2.5. Evaluation of collagen fiber deformation behavior and degradation dependent fascicle structure changes

In order to understand the relationship between tendon fascicle structure and mechanical strains in the tissue, we evaluated associated collagen fiber kinematics within different regions of the sample using a customized script in MATLAB. Based on the structure tensor, SHG signal was used to calculate local collagen fiber orientation within the fascicle samples (Rezakhaniha, Agianniotis et al. 2012) (Avila and Bueno 2015). The fiber orientation distribution within an undeformed ROI (i.e. region between a pair of consecutive photobleach stripes) of a fascicle and ROI strain were used to predict the fiber orientation distribution of respective deformed configuration of the ROI assuming affine deformation by using following relation

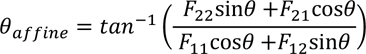

where *θ* represents collagen fiber orientation with respect to fascicle-axis in undeformed configuration and *F*_*ij*_ are elements of deformation gradient matrix of the ROI. For a deformed ROI within a fascicle, *F*_11_ represents the change in displacement between the photobleach stripes (along longitudinal-axis of the tissue) while *F*_22_ represents the change in ROI size in the transverse direction (i.e. fascicle-radius change based on averaged transverse cross-sectional area (CSA) along fascicle-axis). *F*_21_, *F*_12_ were assumed negligible (adjusted to zero) based on both uniaxial deformation of the ROIs along fascicle-axis as well as parallelism of photobleach-stripes upon ROI deformation. Experimentally observed and affined predicted fiber orientation distributions were quantitatively compared using projection plots (see supp. info. S6) (Lake, Cortes et al. 2012) to quantify extent of affine or non-affine for each ROI upon fascicle deformation.

#### 2.2.6. Fluorescence recovery after photobleaching (FRAP)

The kinetics of various size solute particles within mechanically strained regions of the fascicle were quantified using FRAP method. Here, fluorescent BC molecules labelled at N-terminus (for more information on protocol, refer to section 2.2.7) or several size TRITC-dextran molecules (for size specifications of various dextran molecules, refer to table 1) were used as different solute particles in the FRAP measurements. The addition of small volume at 100X concentration of the fluorescent solute solutions (fluorescent BC or TRITC-dextran) to the medium carrying fluorescently labelled mechanically deformed fascicle kept in the bioreactor was made to achieve the desired final concentration. Each sample in the petri-dish carrying the solute solution at 37°C was kept overnight (protected from light with tightly closed lid to avoid evaporation) before making FRAP measurements. Because of the different evaporation rates of the reaction buffer with solutes together with the longer experimental duration (each strained region measurements took ∼2 hours), the compensation evaporation losses from reaction buffer was made by addition DI water (compensation volume was calculated based on respective evaporation rate as shown in Fig. S7) every ∼2 hours. For each solute solution (fluorescent BC or each size TRITC-dextran), different stain sub-volumes (or regions) of 3−4 fascicles were selected for measurements. FRAP data from different locations of each strained sub-volume of the fascicle was normalized and binned using a custom MATLAB script. The obtained data was then used for determination fluorescence recovery characteristics in terms of mobile fraction (*m*) and time constant (*t_1_*) using a single exponential fitting (for FRAP data normalization and curve fitting, refer to section S9).

**Table 1:**
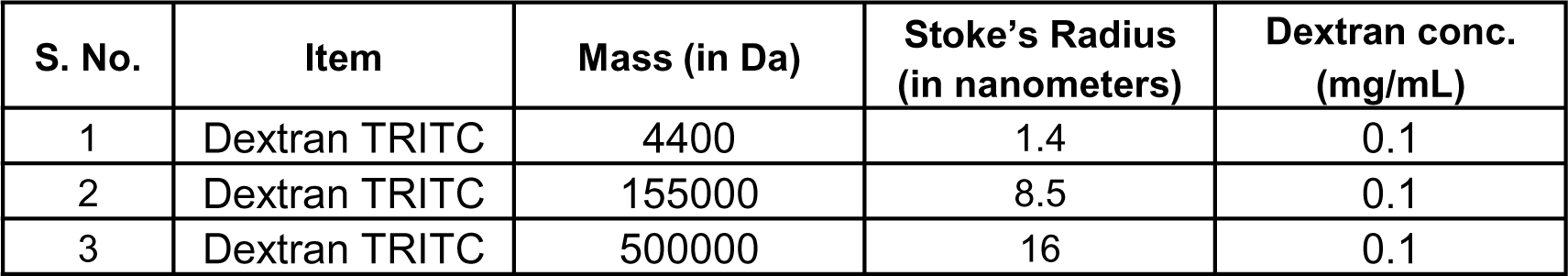
Dextrans – Properties and Concentrations.

| S. No. | Item | Mass (in Da) | Stoke's Radius<br>(in nanometers) | Dextran conc.<br>(mg/mL) |
| --- | --- | --- | --- | --- |
| 1 | Dextran TRITC | 4400 | 1.4 | 0.1 |
| 2 | Dextran TRITC | 155000 | 8.5 | 0.1 |
| 3 | Dextran TRITC | 500000 | 16 | 0.1 |

**Table 2:**
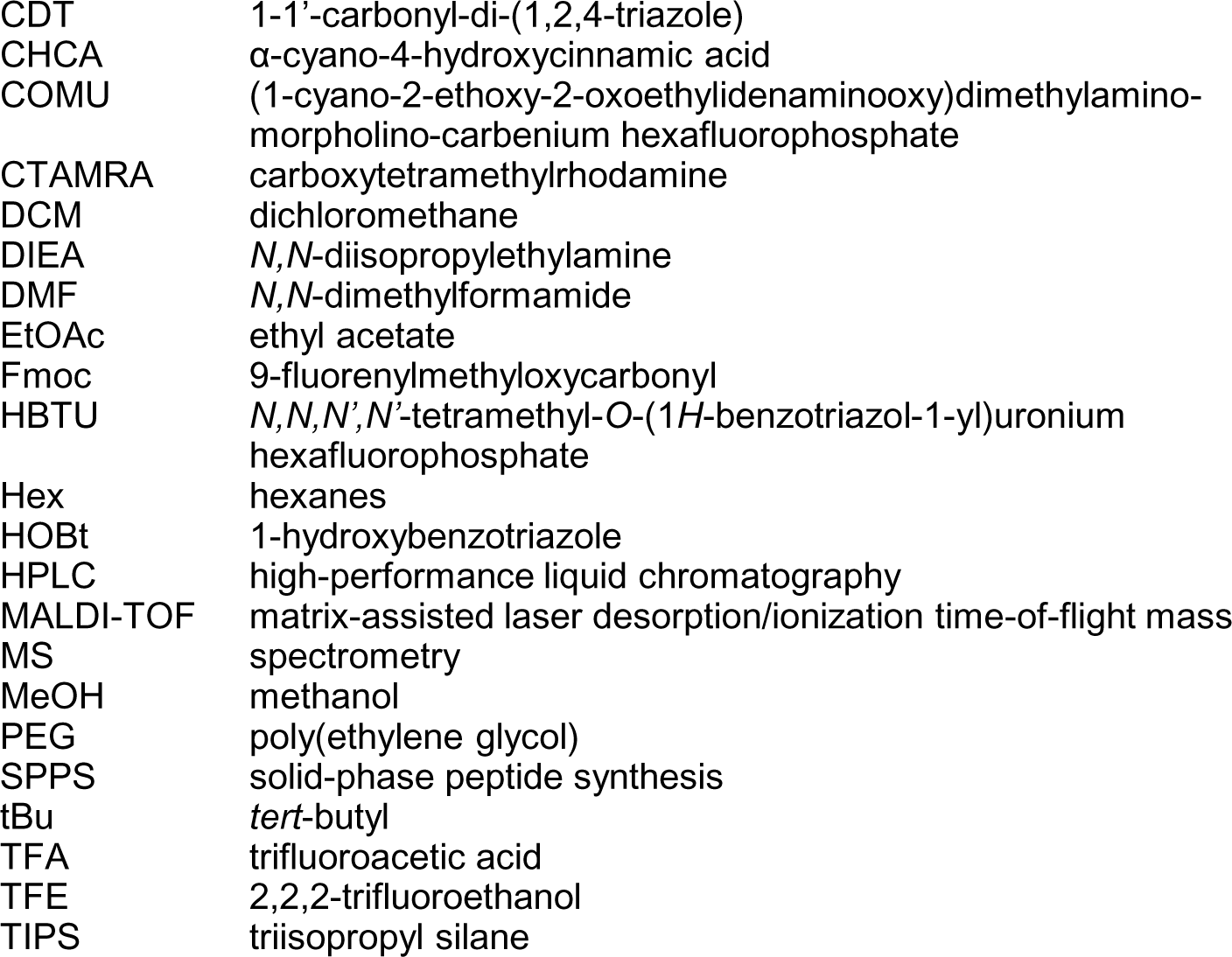
Abbreviations.

|  |  |
| --- | --- |
| CDT | 1-1'-carbonyl-di-(1,2,4-triazole) |
| CHCA | α-cyano-4-hydroxycinnamic acid |
| COMU | (1-cyano-2-ethoxy-2-oxoethylideneaminoxy)dimethylamino-morpholino-carbenium hexafluorophosphate |
| CTAMRA | carboxytetramethylrhodamine |
| DCM | dichloromethane |
| DIEA | *N,N*-diisopropylethylamine |
| DMF | *N,N*-dimethylformamide |
| EtOAc | ethyl acetate |
| Fmoc | 9-fluorenylmethyloxycarbonyl |
| HBTU | *N,N,N',N'*-tetramethyl-*O*-(1*H*-benzotriazol-1-yl)uronium hexafluorophosphate |
| Hex | hexanes |
| HOBt | 1-hydroxybenzotriazole |
| HPLC | high-performance liquid chromatography |
| MALDI-TOF | matrix-assisted laser desorption/ionization time-of-flight mass |
| MS | spectrometry |
| MeOH | methanol |
| PEG | poly(ethylene glycol) |
| SPPS | solid-phase peptide synthesis |
| *t*Bu | *tert*-butyl |
| TFA | trifluoroacetic acid |
| TFE | 2,2,2-trifluoroethanol |
| TIPS | triisopropyl silane |

#### 2.2.7. Fluorescent labeling of BC molecules at N-terminus

For determining the kinetics of collagenase molecules in a mechanically deformed fascicle, fluorescent labelling of BC molecules at N-terminus was done using Alexa Fluor™ 555 NHS Ester (as per manufacturer’s protocol). In order to obtain fluorescent BC molecules, two different stoichiometries of BC with respect to fluorophore (Alexa Fluor™ 555 NHS Ester) were reacted and observed to result different labelling efficiency as indicated by UV (ultraviolet) scanning and HPLC (high performance liquid chromatography) measurements (see Fig. S19 and supp. info. S8).

#### 2.2.8. Artificial cross-linking of fascicle samples

For determining the influence of collagen molecular structural perturbations on degradation of tissue collagen, tendon fascicle samples were artificially cross-linked using ∼1-5 mg/mL TGM solution in DMEM (with FBS+P/S) through ∼24 hours incubation at 37°C. One sub-length of each fascicle sample was artificially cross-linked while the other was used as control to overcome sample to sample variations. After TGM treatment, samples were washed twice with PBS before further processing.

#### 2.2.9. Detection of unfolded collagen content within fascicle samples

For detection of unfolded collagen within samples, Biotin conjugate of collagen hybridizing peptide was used per manufacturer’s protocol while fluorescent Streptavidin conjugate Alexa-594 was used for its detection. Briefly, following endogenous biotin blocking of tissues samples, Biotin-CHP stock solution (50 *μ*M) was heated at 80 °C for 10 min to thermally dissociate trimeric CHP to its monomeric state, and quenched by immersion in 4 °C water for 15 s. Each fascicle sample was kept in 10 *µ*M CHP solution at least for ∼2 hours at 4 °C followed by its twice washing with PBS. Samples were then incubated in fluorescent Streptavidin conjugate Alexa-594 solution in dark for ∼30 minutes and were washed twice with PBS subsequently. Fluorescent imaging of tissue bound CHP was made using two-photon laser (excitation 810 nm, detection 610-660 nm using Hyd detector) by creating z-stacks through tissue sample depth. Composite images obtained using projection of the z-stacks were used to quantify fluorescence from CHP, and intensities were averaged over rectangular areas (at least 50 *µ*m x 50 *µ*m) for making calculations.

#### 2.2.10. Mass spectrometry (LC-MS/MS) of whole heart lysates

Mass spectrometry (LC-MS/MS, or ‘MS’) samples were prepared using procedures outlined (Swift, Ivanovska et al. 2013). Briefly, ∼1 mm^3^ gel sections were carefully excised from SDS–PAGE gels and were washed in 50% 0.2 M ammonium bicarbonate (AB), 50% acetonitrile (ACN) solution for 30 min at 37°C. The washed slices were lyophilized for >15 min, incubated with a reducing agent (20 mM TCEP in 25 mM AB solution), and alkylated (40 mM iodoacetamide (IAM) in 25 mM AB solution). The gel sections were lyophilized again before in-gel trypsinization (20 mg/mL sequencing grade modified trypsin, Promega) overnight at 37°C with gentle shaking. The resulting tryptic peptides were extracted by adding 50% digest dilution buffer (60 mM AB solution with 3% formic acid) and injected into a high-pressure liquid chromatography (HPLC) system coupled to a hybrid LTQ-Orbitrap XL mass spectrometer (Thermo Fisher Scientific) via a nano-electrospray ion source. Raw data from each MS sample was processed using MaxQuant (version 1.5.3.8, Max Planck Institute of Biochemistry). MaxQuant’s built-in Label-Free Quantification (LFQ) algorithm was employed with full tryptic digestion and up to 2 missed cleavage sites. Peptides were searched against a FASTA database compiled from UniRef100 gallus (chicken; downloaded from UniProt), plus contaminants and a reverse decoy database. The software’s decoy search mode was set as ‘revert’ and a MS/MS tolerance limit of 20 ppm was used, along with a false discovery rate (FDR) of 1%. The minimum number of amino acid residues per tryptic peptide was set to 7, and MaxQuant’s ‘match between runs’ feature was used for transfer of peak identifications across samples. All other parameters were run under default settings. The output tables from MaxQuant were fed into its bioinformatics suite, Perseus (version 1.5.2.4), for protein annotation and sorting.

#### 2.2.11. Gel electrophoresis & Coomassie Brilliant Blue stain

Hearts isolated from chick embryos (E4, E6, and E10; n ≥ 8, 4, and 2 per sample, respectively) were rinsed with pre-warmed PBS, excised into sub-millimeter pieces, quickly suspended in ice-cold 1x NuPAGE LDS buffer (Invitrogen; diluted 1:4 in 1x RIPA buffer, plus 1% protease inhibitor cocktail, 1% β-mercaptoethanol), and lysed by probe-sonication on ice (10 x 3s pulses, intermediate power setting). Samples were then heated to 80°C for 10 min and centrifuged at maximum speed for 30 min at 4°C. SDS-PAGE gels were loaded with 5 - 15 *μ*L of lysate per lane (NuPAGE 4-12% Bis-Tris; Invitrogen). Lysates were diluted with additional 1x NuPAGE LDS buffer if necessary. Gel electrophoresis was run for 10 min at 100 V and 1 h at 160 V. Gels were then stained with Coomassie Brilliant Blue for >1 h, then washed using de-stain solution (10% acetic acid, 22% methanol, 68% DI water) overnight.

#### 2.2.12. *Ex-vivo* drug perturbations of embryonic hearts

For actomyosin perturbation experiments, intact E4 hearts were incubated in 25 *μ*M blebbistatin (EMD Millipore, #203390, stock solution 50 mg/ml in DMSO) and were kept away from light to minimize photo-deactivation. The activity of endogenous MMPs was inhibited by ilomastat. Treated hearts were compared to a control sample treated with an equal concentration of vehicle solvent DMSO in heart culture media. For collagen matrix perturbations, E4 heart tissue was incubated in 1 mg/ml concentrations of BC (Sigma, #C7657) for ∼45 min. Enzyme activity was blocked by replacing with chick heart media containing 5% BSA. A minimum of 8 hearts were treated and pooled per lysate/experimental condition.

#### 2.2.13. Probe Synthesis & Purification

##### Solid-Phase Synthesis

General probe synthesis procedures were adapted from those of Mason Smith based on his previous work in peptide synthesis (Smith, Billings et al. 2017). All probes were synthesized on solid phase using Fmoc chemistry. Low-loading rink amide MBHA resin was swelled for 10 min in DCM followed by 3 min in DMF. N-terminal Fmoc group was removed by stirring in 20% piperidine/DMF for 5 min, after which solution was replaced and stirred for an additional 15 min. Resin was then washed 5 times with DMF. Coupling solutions were prepared by combining reagents and DMF solvent in conical vials, mixing, and allowing to activate briefly at room temperature (≥5 min). Each coupling solution contained: 5 eq. Fmoc-protected building block, 5 eq. COMU, and 10 eq. DIEA, with all equivalents determined relative to the resin. Activated coupling solutions were added to deprotected resin and stirred for ≥45 min. Resin was then washed again with DMF (5x). Following coupling of PEG linker during synthesis of probe-1, resin was washed an additional 5 times with DCM. When stopping synthesis at any point, resin was washed an additional 5 times with DCM and dried by vacuum; when resuming synthesis, resin was swelled as described above. These steps of Fmoc deprotection, washing, coupling, and washing were repeated for coupling of each Fmoc-protected building block. Final fluorophore labeling reaction was adapted from established protocols (Stahl, Cruz et al. 2012). Briefly, 6 eq. 5(6)-CTAMRA, 6 eq. COMU, and 12 eq. DIEA were combined in DMF and allowed to activate as described above. Activated coupling solution was then added to deprotected resin and stirred for 3 days. For synthesis of probe-1, resin was subsequently washed with DMF (5x) then stirred in 20% piperidine/DMF for 30 min.

##### Final Cleavage & HPLC Purification

After completing synthesis, each probe was purified as follows: Resin was washed with DMF (5x), followed by DCM (5x), and then dried by vacuum. Probe was cleaved from resin and tBu protecting groups were cleaved from Hyp residues by stirring in TFA:TIPS:H_2_O (95:2.5:2.5) for 1.5-2 h. Crude probe solution was then filtered through frit in reaction vessel into pear-shaped flask. Resin was then was rinsed twice with TFA, and rinses were added to flask. Solvent was evaporated *in vacuo*, then product was resuspended in 10% citric acid or 1N HCl. Organic fraction was extracted ≥3 times with DCM. Remaining aqueous fraction was concentrated to minimal volume *in vacuo*, resuspended in 50:50 H_2_O:ACN, 0.22 *μ*m filtered, and purified by HPLC in gradient of ACN/H_2_O (0.1% TFA) at 80 °C (Due to low yield for synthesis of 6-CTAMRA-PEG control probe-2, it was subsequently determined that organic fraction contained large amount of product. For this probe, organic fraction was dried *in vacuo*, resuspended, filtered, and purified as described above, then combined with purified product from aqueous fraction). Temperature of column/solvent was elevated to prevent oligomerization of probes during separations. Because labeling of probes with 5(6)-CTAMRA produced two isomers (i.e. probes labeled with 5-CTAMRA or 6-CTAMRA), isomers were separated by HPLC during purification and 6-CTAMRA labeled probes were used for subsequent assays (see “Validation Data” below).

##### Preparation of Peptide-Loaded Resins

To streamline general SPPS of CMP-based fluorescent probes, rink amide resin was loaded with peptide sequence shared by multiple probes (i.e. Fmoc-[azGPO(tBu)]_3_-resin). Loaded resin was thoroughly washed and dried, after which small peptide aliquot was cleaved from resin in order to check purity and mass by analytical HPLC and MALDI-TOF MS, respectively. Resin loading density was determined using published UV-vis spectroscopy-based assay. Loaded resin was then stored under desiccation and used as starting material for multiple peptides used in this and other studies, including probe-1.

##### Probe Stock Preparation

Stock concentrations were determined by UV-vis spectroscopy. To prepare samples for UV-vis measurement, concentrated aqueous solutions of probes were diluted and added to 1-cm quartz cuvettes. 4-5 dilutions were prepared from each stock for replicate measurements. UV-vis scans were collected in continuous scan mode with a 200 nm/min scan speed, 1.0 nm data interval, and 1.0 nm bandwidth. Because it has been observed that fluorophores conjugated to N-termini of trimeric collagen peptides can self-quench when peptides assemble into triple helices (Zitnay, Li et al. 2017). The measurements were recorded at 65 °C to inhibit self-assembly. Absorbance of N-terminal CTAMRA fluorophore in each dilution was measured at 555 nm and averaged, after which probe concentrations were calculated using average A_555_ and molar extinction coefficient of 89,000 M^-1^ cm^-1^ (Hyun, Li et al. 2019). Concentrated probe stocks were then aliquoted and dried by vacuum centrifuge for storage prior to use, at which time they were resuspended as needed to achieve necessary concentration for subsequent assays.

## 3. Results

### 3.1 Physiological mechanical strains suppress collagen degradation by exogeneous collagenase(s) in cell-free tissue

Mechanical strain dependence of tissue collagen degradation has been observed at the microscale (i.e. at the size scale of cells) within tendon fascicles consituted by collagen fiber bundles. Collagen fibers, primarily composed of collagen fibrils, are mainly aligned along the fascicle-axis. The spatial distribution of cells and chromatin localization within live fascicle samples obtained from mice tails of WT and NSG mice was demonstrated by Hoechst staining (Fig. S1(*i*-*ii*)). Freeze-thaw cycles, PBS washes and long-term storage (ranging from several weeks to months) of the fascicle samples resulted in a ruptured plasma-membrane, as suggested by chromatin distribution along collagen fibers (shown in Fig. S1-*iii*), and produced a “cell-free” (i.e. protein synthesis free) intact ECM model. Mechanical strain-dependent tissue collagen degradation measurements within the bioreactor (see schematic in Fig. S2 and supp. info. S1) were facilitated for fascicle sizes up to ∼200 *μ*m. Precise quantification of local tissue mechanical strain distribution was made possibile using pattern-photobleaching method on fluorescently labelled samples (see methods, sub-section 2.2.3). Both fluorescent probes novel Aza-peptide or Col-F dye selected for the study showed affinity towards undeformed and deformed tissue configurations (Fig. S3). Three-point bending deformation of fascicle samples, shown in schematic in Fig. 1A, was observed to induce a highly heterogeneous distribution of strain (see Fig. S4C-*ii*). The observed strain distributions varied among three-point bending deformation experiments performed on different fascicle samples (see Fig. S4E). The deformed configuration of fascicle within the bioreactor was observed to be retained at least during the initial phase of tissue degradation upon addition of exogeneous collagenase. The retainment of tissue in deformed configuration at least during the initial hours of degradation is likely to result from the forces produced from adhesion to bottom polystyrene surface and the contact with a glass-corner surface in the case of fascicle samples. The observed difference in the strain distributions among different experiments (Fig. S4E) is an outcome of non-uniform sample deformation and not because of strain-quantification method. The robustness of the strain quantification method for each sample is duly supported by the same fluorescent labelling protocol, constant number of data points selected for Gaussian fitting, same size/spacing of photobleach stripes (Fig. S4B and Fig. S4C). Because strain magnitude calculations are based on undeformed (strain-free) fascicle configuration, the minimization of residual stresses within undeformed samples during our experiments is acheived by careful landing of the sample on the bottom plate of the bioreactor. Here, it is important to keep in mind that the application of a very small force (represented by toe-region of “stress-strain” curve for rat tail tendons (Fratzl, Misof et al. 1998)) is capable of straightening the microscopic collagen fiber crimps (Diamant, Keller et al. 1972). Our observation of the presence of microscopic crimps in SHG images of unstrained fascicles (lower-left panel of Fig. 1B at -1.50 and 0.20 hours) ensured the required minimum residual stresses.

**Fig. 1:**
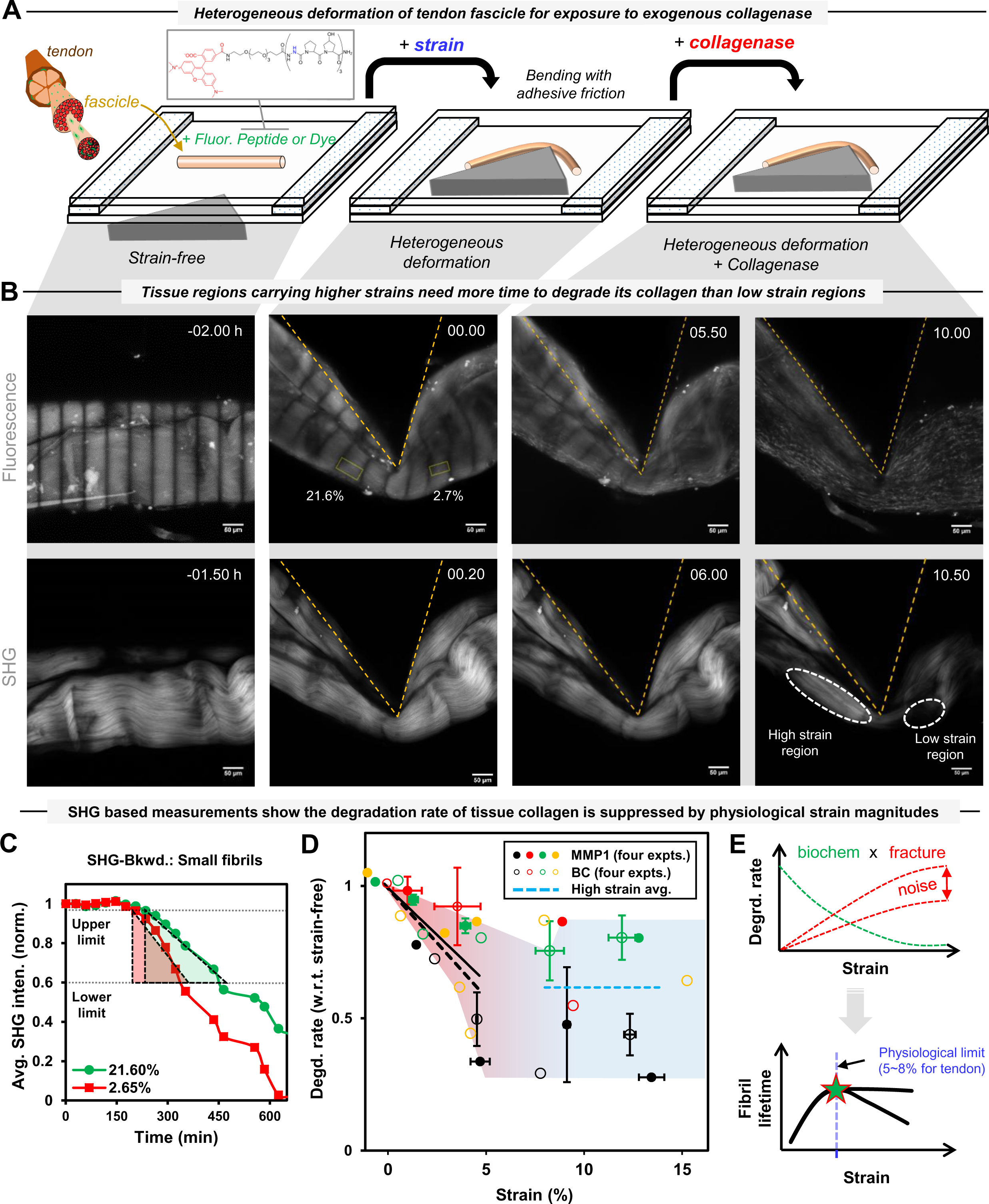
Heterogeneously deformed mice tail tendon fascicle resists degradation by exogeneous collagenase(s) in strain magnitude dependent manner independent of collagenase site specificity on collagen molecules. **A.** A schematic of heterogeneous deformation of a fluorescent fascicle (labelled by Aza-peptide or Col-F dye) in the bioreactor by three-point bending, with fascicle’s deformed configuration is retained by its adhesion to bottom polystyrene surface, for subsequent exposure to matrix metalloproteinase-1 (MMP-1) or bacterial collagenase (BC). **B.** Enzymatic degradation of a heterogeneously deformed fascicle at different time-points (L to R; in hours) of an experiment. Fascicle’s unstrained and strained configurations (at various time points) are indicated by pattern photobleached fluorescent (top-panel: L to R) and backward SHG (bottom-panel: L to R) composite images (i.e. projected sum). The SHG image of a fascicle shows collagen fiber crimps that straightening up upon mechanical deformation. The disappearance of SHG signal upon addition of collagenase shows collagen degradation. **C.** Collagen degradation rate of each strained region is measured based on the slope of the average SHG intensity change versus time during initial phase of tissue degradation (for details, see Fig. S5 and supp. info. S3). **D.** Mechanical strain magnitudes of up to ∼5% tightly suppress the degradation rate of fascicles by collagenase(s) as shown by the similar respective slope values (−0.07 and −0.08) for line fits (solid: MMP-1 and dashed: BC) to respective degradation rate versus strain data points in the separate set of four experiments for each collagenase despite the difference in site specificity of MMP-1 and BC on collagen molecules. For mechanical strains value > ∼8%, the degradation rate becomes almost independent of mechanical strain changes (dashed blue line). Fig. S10 shows the data for MMP-1 and BC separately. Error bars indicate ±SEM. **E.** Mechanical strain dependent degradation suppression of tendon tissue collagen content appears to be an outcome of biochemical regulation and fracture within the tissue (top). Mechanical strains up to ∼5-8% (i.e. physiological limits for tendon tissue) tend to maximize collagen fibril lifetime while higher noise due to tissue fracture starts dominating at larger strains magnitudes (i.e. pathological strains).

The abundance of fibrillar collagen within fascicle tissue samples (∼70-80% fibrillar collagen as their dry weight) and the sensitivity of SHG signal towards fibrillar collagen content of a tissue enhanced the accuracy of our fibrillar collagen content measurements. In general, SHG signal reflects a measure of electric susceptibility resulting from the permanent dipole-moment and alignment of the molecules constituting a material and therefore, reflects a material bulk property (Chen, Nadiarynkh et al. 2012). In tissues, besides collagen type-I and -II molecules (Campagnola 2011) assembled into fibrils, myosin within acto-myosin complexes (Campagnola, Millard et al. 2002) (Nucciotti, Stringari et al. 2010) also show non-zero values of susceptibility while collagen type IV and type III do not produce sufficient SHG signal for imaging (Pena, Boulesteix et al. 2005) (Cox, Kable et al. 2003). In tendon fascicles, the most abundant collagen-I molecules are assembled into collagen fibrils, which are further assembled into collagen fibers, and produce good SHG signal intensity. Theoretically, SHG signal intensity is a measure of the number of collagen molecules within a focal volume and the orientation of collagen fibrils (with respect to both the laser polarization angle and the laser propagation direction) (Mertz and Moreaux 2001, Stoller, Kim et al. 2002, Stoller, Reiser et al. 2002, Stoller, Celliers et al. 2003, Williams, Zipfel et al. 2005, Odin, Guilbert et al. 2008, Pavone and Campagnola 2014). The collagen molecules, within a collagen fibril and behaving as dipoles, are primarily aligned along the fibril longitudinal-axis. SHG signal is maximized for a collagen fibril (lying in a plane perpendicular to the laser propagation direction) that is lying parallel to the incident electric field and is minimized when the fibrils are oriented perpendicular (Odin, Guilbert et al. 2008). During collagen degradation, time dependent tissue collagen content measurements using a label-free SHG imaging method is considered to be more accurate (Fig. S17) than those based on fluorescent labelling (Fig. S18) due to several factors including the extent of labeling (or quenching), photobleaching and the specificity of the fluorophore, and thus, the use of SHG imaging tool within our study sounds a reasonable choice.

Following heterogeneous deformation within the bioreactor, a fascicle sample was exposed to exogeneous active collagenase (purified MMP-1 or BC) while being imaged for mechanical strains and collagen content. Local fascicle strains (based on photobleach stripe position, Fig. 1B, upper-panel) and tissue collagen content (indicated by SHG signal intensity in Fig. 1B lower-panel) at different stages of the experiment were used to calculate the strain dependence of collagen degradation. For each ROI, the local fascicle strain magnitude and the corresponding rate of SHG signal loss was measured to estimate corresponding collagen degradation rate (for two representative ROIs, see Fig. 1B (top-panel, second from left) and Fig. S5A). The degradation rate of each ROI using SHG signal in backward (Fig. 1C) and forward (Fig. S5A) directions is calculated within initial phases of the experiment followed by the addition of collagenase. Fluorescent labelling of the fascicles either with Col-F or aza-peptide in the control experiments was observed not to interfere with the degradation rates of undeformed fascicles by BC (Fig. S5C). Considering the uneven spatial distribution of collagen fibrils within the tissue, the SHG intensity was averaged over each ROI. The change in average SHG intensity with time for each ROI (representing a known mechanical strain magnitude) was used for comparison among several ROIs of a deformed fascicle sample (Fig. S5B(*i*-*ii*) based on backward and forward SHG signal) in terms of local tissue collagen degradation rates. Considering the relative higher cleavage efficiency of BC than MMP-1 (> ∼10 times, see Fig. S6) on monomeric collagen (Garnero, Borel et al. 1998), the samples were continuously imaged during degradation experiments involving BC. Despite the observed difference in absolute degradation rate of MMP-1 and BC on monomeric collagen (see Fig. S6), the time taken for complete degradation of tendon fascicles were similar (ranged upto several hours for MMP-1 and BC when maintained at nearly similar concentrations of ∼100 nM).

To assess the contribution of reaction buffer evaporation from the bioreactor, the changes in pH of reaction buffer and tissue collagen fibril packing were quantified to represent the alterations in reaction buffer composition. The evaporation of the reaction buffer carrying different solutes (i.e. BC or dextran) from the edges of the bioreactor was not observed to change pH values considerably (Fig. S7(*i*)). The compensation for evaporation losses from the micro-bioractor during the experiments was made every ∼2 hours by adding DI water based on corresponding evaporation rates of different reaction buffers (Fig. S7(*ii*)). Further, the effect of reaction buffer composition changes involving different solutes (i.e. dextran, calcium chloride (CaCl_2_) salt or sodium chloride (NaCl) salt) on tissue collagen fibril packing was measured by means of forward and backward SHG signal (Fig. S8). Varying dextran concentrations did not affect tissue collagen packing (Fig. S8A) but the increase in the concentrations of CaCl_2_ or NaCl (Fig. S8B and Fig. S8C respectively) promoted collagen fibril disassembly as indicated by a decrease in forward/backward (F/B) signal ratio. The superposition of mechanical strain on the tissue samples, in addition to presence of CaCl_2_ or NaCl, resulted in further changes in F/B ratios (orange markers in Fig. S8B). Salt concentration-dependent and mechanical strain-associated changes in the SHG signal are indicative of changes in tissue molecular architecture resulting from altered molecular packing within collagen fibrils and/or assembly of smaller collagen fibrils into larger fibrils.

Considering the influence of non-linearity of SHG signal (Pavone and Campagnola 2014) and other factors (mentioned below) on our measurements, the calculation of relative degradation rates among different ROIs (carrying different strain magnitudes) of a fascicle sample was made through several normalizations of the data. First, in order to measure the relative change in SHG signal with time and the corresponding slope-based degradation rates, the average SHG signal value from each ROI at various time points was normalized with respect to that of initial signal magnitude (at time *t* = 0). Secondly, the comparison of the degradation rates of ROIs within a sample with respect to that of a strain-free ROI was made using further normalization. In this step, the normalization of degradation rates for all ROIs of a fascicle with respect to that of strain-free region of fascicle or extrapolated degradation rate value at zero strain (by linear fit to data points up to ∼5% strain) was performed. The above mentioned normalization procedures helped comparing the mechanical strain-dependent degradation rates among different fascicle samples while minimizing the influence of several factors including sample-to-sample variation, changes in collagenase activity with time (before/after addition to reaction buffer) and the influence of evaporation dependent salt concentration changes. For each mechanically deformed fascicle exposed to a collagenase (MMP-1 or BC), the normalized degradation rates of its ROIs with different strain magnitudes are shown in Fig. S9. When superimposed (Fig. 1D), the degradation rates by MMP-1 or BC showed a linear suppression for strain magnitudes up to ∼5-8%. The superposition of degradation-strain data for MMP-1 and BC (Fig. 1D) showed same level of suppression of collagen degradation rate (indicated by similar slopes of line fits to the data-sets for MMP-1 and BC) at least for tendon tissue strain values up to ∼5%. Although noisy at strain values beyond ∼8%, the suppressed degradation rates for the fascicles were observed to be almost independent of further increases in the strain magnitudes.

### 3.2. Strain-suppression of the degradation rate is independent of mechanical deformation mode, and local tissue strain, degrdation rate and organization of collagen with time during degradation are correlated

The interrelationship between collagen degradation-dependent local tissue mechanical strain and structure changes was investigated in the case of cell-free fascicles deformed in a physiological mode i.e. uniaxial deformation. Interestingly, the strained sub-lengths of uniaxially deformed fascicles were observed to deform non-uniformly along the fascicle-axis. It must be noted here that the non-uniform strain distribution along fascicle-axis is not an outcome of any adhesion-friction (between sample and bottom glass-surface) because the samples floating in medium while uniaxially deformed showed a similar behavior. The strain dependence of the degradation rate for different ROIs (Fig. 2A(*ii*)) in the case of uniaxially deformed fascicles was observed similar to that of heterogeneously deformed (Fig. 1D). The strain-dependence of the fascicle degradation rate independent from micro-scale tissue deformation mode suggests tissue collagen degradation is intrinsic material property depends only on molecular composition and sub-micron scale packing.

**Fig. 2.**
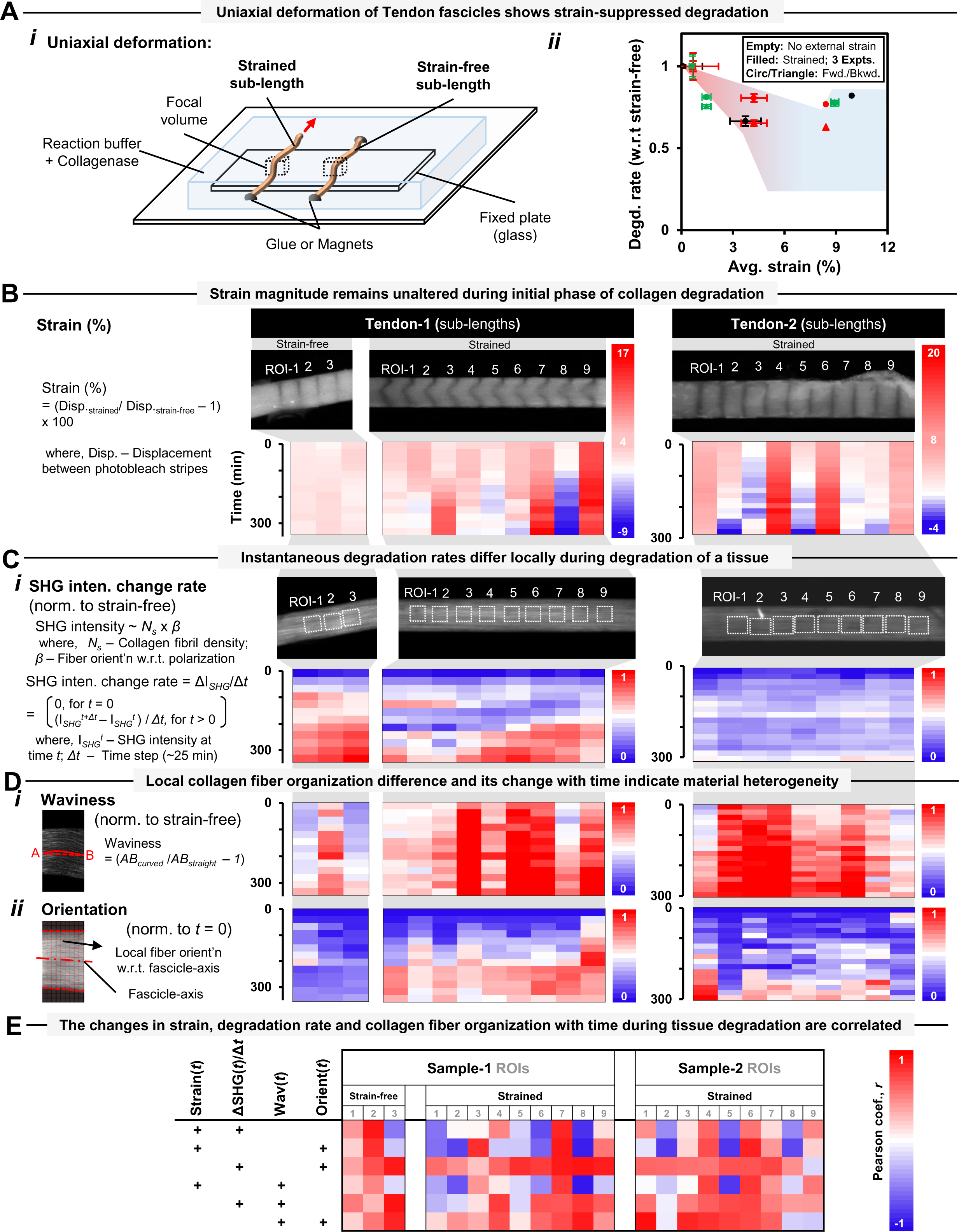
Strain suppression of tendon fascicle degradation rate is independent of mechanical deformation mode. The magnitudes of strain, degradation rate and collagen fiber organization differ spatially within a tissue but their changes with time during tissue degradation are correlated. **A. (*i*)** Schematic representation for uniaxial deformation test of a fascicle with its strain-free and strained sub-lengths simultaneously exposed to BC. Strains and collagen content are quantified based on pattern photo-bleaching and SHG imaging respectively. **(*ii*)** Initial strain dependence of average degradation rate of ROIs present within uniaxially strained and external strain-free sub-lengths (grouped based on < or > ∼8% strain; filled and empty markers respectively). **B.** Local strain magnitudes within ROIs of a uniaxially strained sub-lengths remain unaltered initially during degradation by BC whereas later phases show local strain relaxation. For ROIs within external strain-free sub-length of a fascicle, the strain values remain almost unaltered throughout the experiment. **C.** Local difference in instantaneous degradation rates within fascicle sample, measured based on change in forward SHG signal, are evident both within and between strain-free and strained sub-lengths. **D.** Local fascicle collagen fiber organization in terms of collagen fiber waviness (over ∼50 µm along fascicle-axis) and orientation (defined by relative number of collagen fibers inclined compared to aligned with fascicle-axis over each ∼2 x 2 µm window) and its change with time different locally within each sub-lengths. For calculations, see supp. info. S7. **E.** The values of strain, instantaneous degradation rate and collagen fiber organization (represented by waviness and orientation values) with time during tissue degradation are highly correlated. The change from positive to negative correlation with a fascicle is likely to result from residual internal stress relaxation as a result of collagen degradation.

The comparison of fascicle sub-lengths (i.e. strain-free and strained) structure with time, when degraded by BC, has been quantified in terms of strain magnitudes, instantaneous degradation rate of collagen and organization of collagen fibers within each ROI of a fascicle region as shown in Fig. 2B-2D. Strain magnitudes within both sublengths (i.e. strain-free and strained) remains almost unaffected during the initial phase of fascicle degradation (first ∼2 hours as shown in Fig. 2B). The observation of unaffected local tissue strains in the initial degradation phase are in agreement with our assumption of the degradation rate calculation based on initial tissue strain values while maintaining constant substrate concentration. Beyond the initial phase of fascicle degradation, local strain relaxation was observed for ROIs within strained sub-lengths (as indicated by strain fluctuations with time in Fig. 2B) whereas the strain values within strain-free sub-length did not change with time. Qualitative observation of overall instantaneous degradation rate with time (Fig. 2C) showed a higher degradation rate of strain-free sub-length of the fascicle than that of strained sub-lengths.

In order to ascertain the relationship among strain, collagen degradation rate and collagen organization within a tissue, we quantified local collagen fiber organization changes within a uniaxially deformed fascicle exposed to BC with time. Here, the local collagen fiber organization within a fascicle has been quantified by means of two parameters i.e. waviness and orientation (i.e. inclined to aligned fiber numbers) within each ROI, as shown in Fig. 2D(*i-ii*). Slicewise waviness, a measure of change in the parallel collagen fibers orientation when moving along fascicle length, is measured within each ROI on a single slice (or longitudinal plane) parallel to the fascicle-axis (for details, refer Fig. S11 and supp. info. S7). Slicewise waviness value of zero indicates all collagen fibers are parallel to each other. Here, it is important to mention that the slicewise waviness values were observed to vary within a fascicle when moving along its transverse direction (i.e. among longitudinal slices of a image stack as shown in Fig. S10A). The difference in slicewise waviness values within an ROI indicates a phase lag between parallel collagen fibers of fascicle, and therefore we considered median of slicewise waviness values to represent ROI local waviness for comparison among different ROIs of a fascicle. As shown in Fig. 2D-*i*, waviness values of ROIs (normalized with respect to respective strain-free configuration) are observed to be different initially (at time *t* = 0) in case of both strain-free and strained sub-lengths. The difference in waviness values among ROIs of a fascicle suggests local difference in collagen fiber organization within a fascicle. The changes in waviness values with time during the degradation of a strain-free and strained sub-length indicated sudden bursts or changes (Fig. 2D(*i*)) that might have resulted from local stress relaxation within collagen fibers. Although noisy, the qualitative increase in ROI waviness values with time was more rapid in case of strained sub-length than that of strain-free sub-length of a fascicle. On the other hand, the orientation values for a fascicle ROI compares the relative number of inclined collagen fibers (with respect to fascicle-axis) to that of aligned ones (for details, refer Fig. S10 and Fig. S12, and supp. info. S6). The orientation values (normalized with respect to value at time *t* = 0) for all ROIs increased considerably with time in the case of strained sub-length of fascicle when compared to that of strain-free sub-length. The difference between strain-free and strained sub-lengths of the fascicle in terms of the orientation value change with time (Fig. 2D-ii) qualitatively demonstrates tissue strain-dependence of local collagen fiber orientation distribution. Precise comparison of the orientation magnitudes in terms of range values (normalized with respect to respective values at time *t* = 0; for range value calculation, see supp. info. S6) with time in case of strain-free and strained sub-lengths of a fascicle (Fig. S13A) show a relative larger increase with time within the strained sub-lengths as compared to that of strain-free. The scatter-plots of these parameters (i.e. waviness or orientation) with strain values and corresponding fits to respective binned values (i.e. local averaging based on strain values) are shown in Fig. S13. The increase in waviness and orientation (indicated by inclined/aligned fiber ratio) magnitudes with time suggests the persistence of inclined collagen fibers of a strained fascicle relative to the aligned ones within a degradative environment, and has been represented by the schematic as shown in Fig. S13B.

The changes in strain, collagen degradation rate and collagen organization values among various ROIs of strain-free and strained sub-length with time during degradation (shown in Fig. 2B-2D) were used to observe correlation among these parameters. The corrleation is represented by means of Pearson’s corrleation coefiecient as shown in Fig. 2E where strain, collagen degradation rate and collagen organization values were observed to be highly correlated.

### 3.3. Fascicle collagen fibers might follow non-affine deformation but the deformation does not influence local permeation, tissue mass density or mobility significantly

Based on the observed strain-dependent changes in parameters (i.e. local strain, collagen content change and collagen fiber organization) with time in case of a fascicle exposed to collagen degradation (shown in Fig. 2), we further investigated strain-dependent local fascicle structure changes in the absence of tissue collagen degradation. It is important to note here that uniaxial deformation of the fascicle resulted in non-uniform strain distribution among various ROIs of a fascicle (Fig. 2B). Based on mechanical strain-dependent local tissue structure changes in terms of collagen fiber orientations within each ROI, we evaluated if collagen fibers deformation within each ROI (i.e. the region between two consecutive photobleach stripes) of a fascicle followed affine deformation or not (for details, refer to methods 2.2.5, Fig. S10 and supp. info. S6). For each uniaxially deformed fascicle, the changes in the average cross-sectional area and longitudinal elongation of each ROI (geometry defined by forward SHG signal) were determined to obtain corresponding stretch ratio values. The obtained stretch ratio of a ROI and collagen fiber orientation distribution in its undeformed configuration were then used to predict theoretical collagen fiber orientation distribution in deformed configuration of the ROI under affine assumptions. The difference between experimentally observed and affine predicted collagen fiber orientation distributions (Fig. 3A(i)) for all ROIs was made based on offset values using projections plots (for calculation of offset values, see supp. info. S6 and Fig. S10). Thus, the offset values indicate a measure of non-affine deformation behavior. Interestingly, the local deformation of collagen fibers within a ROI, whether affine or non-affine, was found to depend on the primary orientation of collagen fibers in the undeformed tissue configuration. Here, primary orientation of collagen fibers is based on the peak of Gaussian curve fitted to collagen fiber orientation distribution (for details, see supp. info. S6). Primary inclination angle between primary orientation of collagen fibers and fascicle-axis was observed to determine the affine or non-affine nature of collagen fiber reorganization within a deformed tissue. The regions of fascicle carrying collagen fibers inclined to the fascicle -axis were observed to follow non-affine deformation behavior while regions carrying collagen fibers oriented along fascicle-axis followed affine deformation (shown in figure 3A-ii as filled circles). The extent non-affine deformation within the tissue was found to increase linearly (with R-square = 0.9119) with the increase in the primary inclination angle with respect to fascicle-axis. The global deformation behavior of collagen fibers within a fascicle sample at larger length-scales, predicted by considering the region between two extreme photobleach stripes (∼500 *μ*m apart covering nine ROIs), has been observed as affine (represented as empty circles in Fig. 3A(ii)). As a side calculation, the local tissue non-affine behavior quantification in terms of the product of offset and range values (as represented in Fig. S14A) also showed a linear increase (with R-square = 0.88) as a function of primary inclination angle magnitude. The non-uniform strain distribution within different ROIs of a fascicle subjected to uniaxial deformation varied depending on the primary inclination angle magnitude (as represented in Fig. S14B) but the correlation between these parameters was not clear. The magnitude of primary inclination angle of collagen fibers within an undeformed fascicle seems to control local reorganization of collagen fibers within a fascicle tissue during its mechanical deformation, and might be influencing local tissue heterogeneity but cannot be concluded as the sole factor controlling local fascicle stiffness values.

**Fig. 3.**
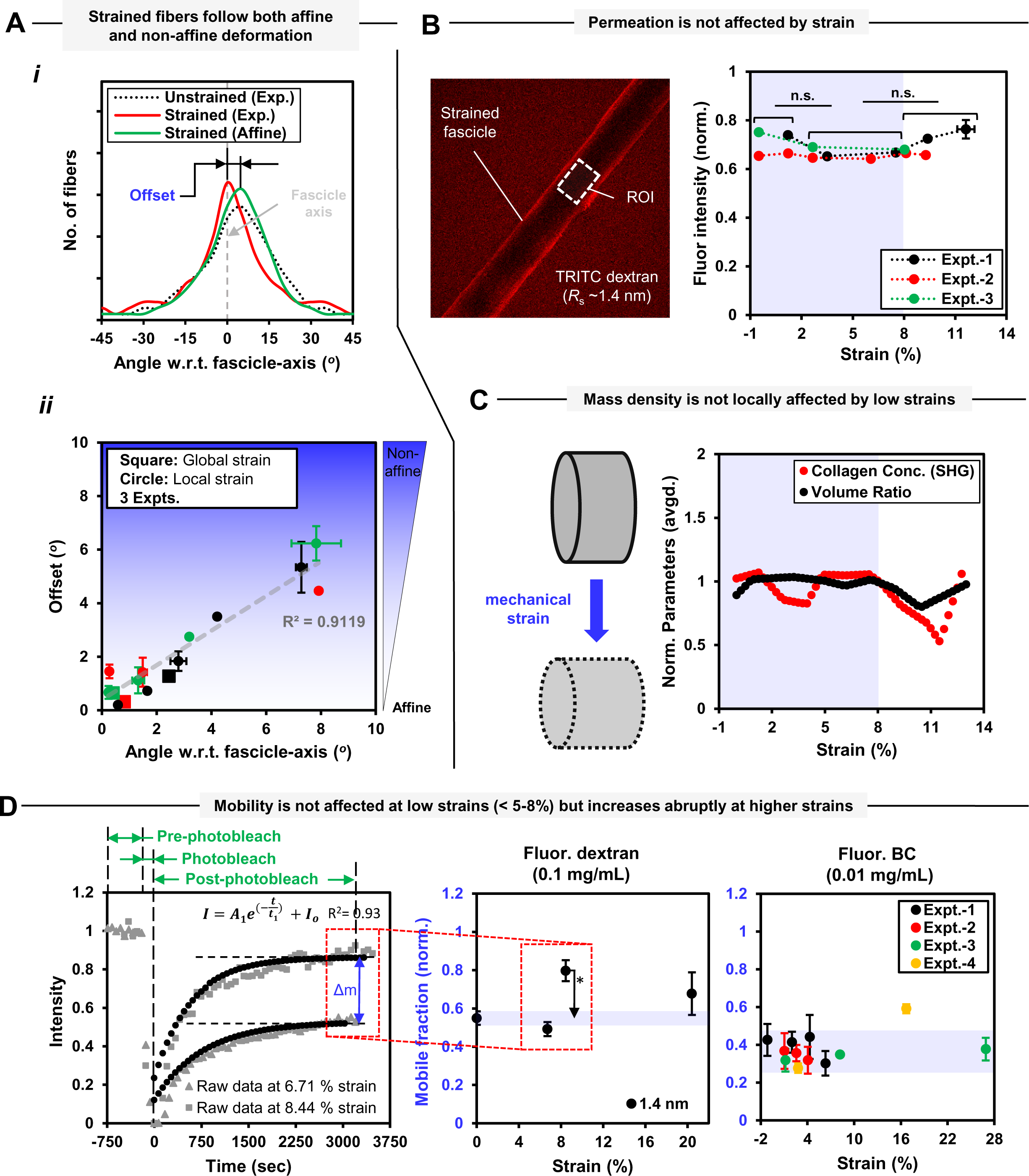
Strain-dependent tendon fascicle structure changes in the absence of tissue collagen degradation. Tendon fascicles tend to follow non-affine deformation locally at size scale of cells whereas the global deformation of the fascicle remains affine. Physiological mechanical strain magnitudes present within the tendon fascicle do not affect local tissue mass density, solute permeation and mobility. **A.** Local non-affine behavior (within each ROI) of a fascicle under uni-axial deformation defined in terms of offset magnitude increases linearly with the increase in primary fiber inclination angle with respect to to fascicle-axis. **(*i*)** Fiber orientation distributions (with respect to fascicle-axis based on quantile grouping) within a representative ROI in undeformed/deformed configurations and affine predicted fiber orientation distribution. Offset values quantify the deviation between experimentally observed and affine predicted fiber orientation distributions in deformed configuration of fascicle. **(*ii*)** Non-affine deformation is prominent in the regions carrying primarily inclined fibers with respect to fascicle-axis. For a fascicle, local non-affine behavior (within consecutive photobleach stripes) is indicated by filled markers and global behavior (within extreme photobleach stripes) is represented as empty markers. (for offset calculation using projection plots, see supp. info. S6). **B.** Solute permeation within fascicle is strain independent for strain magnitudes within physiological limits. (L) Real-time image (at z-position position equal to ∼half diameter) of a mechanically strained fascicle kept overnight in 1.4 nm Stoke’s radius (*R_s_*) TRITC-dextran solution at (0.1 mg/mL). (R) The solute concentration of TRITC-dextran molecules (1.4 nm Stoke’s radius and closer to collagenase molecules) estimated on the basis of average fluorescence intensities (normalized with respect to intensity of free dextran solution) in the strained regions of the fascicles. For more info., see Fig. S15 **C.** Average values of collagen concentration and volume ratio (piece-wise line fits to normalized parameter values) fluctuate with increasing fascicle strain magnitudes but the local tissue mass density (as indicated by product of parameters, see Fig. S17) remains within ±10% of at least for strain magnitudes up to ∼8%. **D.** Enzyme mobility within mechanically strained regions of a fascicle is uninfluenced by mechanical strain magnitudes up to ∼8%. Mobile fractions of 1.4 nm TRITC-dextran and fluorescent-BC molecules remains unaffected with mechanical strain up to ∼8% and undergo a sudden increase at larger strains indicating local fracture within the fascicle. Mobile fraction magnitudes of fluorescent-BC molecules is slightly lower than similar size TRITC-dextran molecules. For details on molecular kinetics of various size TRITC-dextran and fluo.-BC molecules, refer to Fig. S20. For fluo. labelling of BC at N-terminus, see supp. info S19. **\* = p < 0.05, ** = p < 0.01, *** = p < 0.001.**

Several possible molecular mechanisms that could result in strain-suppressed collagen degradation were explored by investigating the influence of local tissue mechanical strain magnitude on the characteristics of collagen-collagenase system. The characteristics of the collagen-collagenase system including solute permeation, local tissue mass density and solute mobility (Fig. 3B-D) were investigated in case of heterogeneously deformed fascicle samples. An image (at z-position equal to ∼half diameter of a fascicle) of a mechanically strained fascicle (Fig. 3B (L)), kept overnight in collagenase-size TRITC-dextran molecules (Stoke’s radius = 1.4 nm) at 37*°C*, shows the local fluorescent intensity distribution. The higher fluorescent intensity at the edges of the fascicle might have resulted from the sample curvature or higher dextran concentration near the sample surface. For different fascicle samples, the comparison of the average fluorescent intensity (normalized with respect to average fluorescent intensity values of free dextran solution surrounding the fascicle) was made with different regions of the fascicle, and experimental artifacts resulting from the measurements either close to fascicle surface or fluorescent signal attenuation (due to sample thickness) were excluded by the averaging of fluorescent intensity values over sub-volumes within a fascicle ROI located at same distance from fascicle surface (for more information, see Fig. S15). As shown in Fig. 3B (right), the solute permeation into mechanically strained sub-volumes of each fascicle in terms of average fluorescent intensity has been observed to vary by less than ±10%, thus suggesting tissue strain independence of solute permeation for strain magnitudes up to ∼8%.

The effect of tissue strain on local tissue mass density has been quantified via means of the changes in local collagen content calculated based on absolute SHG signal because SHG intensity is proportional to square of collagen concentration and polarization direction (Raub, Tromberg et al.). Considering the influence of tissue depth on SHG signal attenuation and polarization, the volumetric average value of SHG signal within a sub-volume (at certain depth) of each ROI of a tissue was obtained and subsequently scaled based on the inclination angle collagen fibers (with respect to polarization direction of incident light) to obtain absolute SHG signal intensity for each ROI (for details, refer to Fig. S16 and supp. info. S3). The absolute SHG signal value (obtained separately for forward and backward directions) at different strain values were normalized with respect to SHG signal values from corresponding ROIs in strain-free configuration (or the extrapolation of values for up to ∼5% strains) and averaged at each strain magnitude as shown in Fig. S17A. The estimation of changes in tissue mass density values at various strain magnitudes, collagen concentration (based on SHG signal) within deformed fascicle ROIs was further normalized with respect to the change in respective ROI volume at corresponding strain (see Fig. S17A). Here, volume ratio (VR) measurements (VR = V_deformed_ /V_undeformed_) were made for each ROI for its deformed and un-deformed configurations (see Fig. S17B) within a fascicle where the fascicle geometry was based SHG signal (average of forward and backward signal). The piecewise fits to both collagen concentration (based on normalized SHG signal) and normalized VR values were obtained as a function of strain magnitude within heterogeneously deformed fascicles (Fig. 3C) to quantify corresponding tissue mass density variations. Although the fluctuations in the collagen concentration and VR values as a function of strain were observed, the tissue mass density values stayed within +/-10% limit at least for strain magnitudes up to ∼8% as indicated by the product of two parameters (collagen concentration and VR) as shown in Fig. S17C. Similar to the strain dependence of volumetric changes within heterogeneously deformed fascicle samples (red-dots in Fig. 3C), the uniaxially deformed fascicles were also observed to undergo a volume-loss that remained within +/-10% limit for strain values up to ∼8% as shown in Fig. S17D. As side note, the calculations of collagen concentration and VR based on fluoresent signal (due to Col-F labelling) of fascicles were also made (Fig. S18) but were not considered for tissue mass density analysis due to advantages associated with SHG signal over the fluorescent signal. Based on SHG based collagen content measurments, strain-induced tissue collagen mass density changes are present within fascicle samples but the changes are not significant for tissue strain magnitudes up to ∼8%.

In order to elucidate local tissue strain-dependent solute mobility, the kinetics of various size dextran molecules and fluorescently labelled BC molecules in terms of mobile fraction and time constant values was measured using FRAP method (see Fig. S20). As recovery kinetics of molecules during FRAP measurements usually depend upon experimental setttings such as bleach area, bleach geometry, time of bleach, acquisition bleaching and concentration of solute, these variables were either kept constant among various experiments or data were normalized (for calculations and various settings, supp. info S9). We selected three different sizes of fluorescent dextran molecules with Stoke’s radii of 1.4, 8.5 and 16 nm (for size specifications of various dextran solutions, refer to Table 1) and BC molecules that were fluorescent labelled only at N-terminus (for fluorescent labelling of BC at N-terminus, see supp. info. S9 and Fig. S19). Mobility of dextran molecules within unstrained fascicles has been observed to depend on the size of the molecules (see Fig. S20A), but the larger molecules appear more mobile counter-intuitively. For strain magnitudes of up to ∼8%, strain suppressed mobility was only observed for larger size (16 nm Stoke’s radius) dextran molecules (see Fig. S20B) while the relatively smaller–size molecules (1.4 nm and 8.5 nm TRITC-dextran or fluorescent-BC) didn’t show any tissue strain dependence. Interestingly, the fascicle regions carrying larger strain values (> ∼8%) showed larger mobile fraction values indicated by a sudden increase for both fluorescent dextran (1.4 nm) and BC molecules when compared to the values at low strain values (shown in Fig. 3D and Fig. S20). At larger tissue strain magnitudes, mobile fraction values for fluorescent dextran (1.4 nm Stoke’s radius) and BC showed a similar strain magnitude dependence qualitatively but differed in the magnitudes. The mobile fraction values of fluorescent-BC molecules were observed to be lower than 1.4 nm dextran molecules (see Fig. 3D (center and right)) while respective time constant values have been found relatively higher (see Fig. S20 B-C), indicating slower rate of flow for fluorescent-BC molecules as compared to dextran molecules within the strained fascicles. The change in mobility of collagenase size molecules at larger strain magnitudes within fascicles (in Fig. 3D) suggest critical changes in tissue structure. The changes in fascicle molecular structure at strain values larger than ∼8% were also evident in strain-dependent molecular unfolding of collagen molecules present within the fascicles (see Fig. S21) at strain values larger than ∼8% where collagen hybridizing peptides (CHP) showed increased binding to unfolded collagen trimers within the tissue (Zitnay, Li et al. 2017).

### 3.4 Contractility of developing chick hearts maintain tissue collagen-I levels in the presence of endogeneous collagenases

In order to understand the mechanical strain-dependent collagen degradation within a live tissue by endogeneous MMPs, isolated beating chick hearts were investigated with tissue contractility or beating considered as a measure of the mechanical strain. In general, the presence of vascularisation, various cell types, etc., within adult heart tissues result in complexity of cell-ECM interactions (Gonzalez-Rodriguez, Guevorkian et al. 2012) (Chien, Domian et al. 2008). Importantly, the embryonic development of chick heart tissue to E4 stage parallels the expression of collagen-I in addition to other excitation-contraction proteins. Additionally, the stiffness of E4 hearts, primarily determined by collagen type-I levels, is optimal for the beating of embryonic heart cells (Majkut, Idema et al. 2013) (Chiou, Rocks et al. 2016). The use of mechanical signaling by heart cells for beating of E4 heart tissues, because electrical signaling dominates in case of highly complex adult heart tissues, was our basis for the selection of E4 chick heart tissues for cell based mechanical deformation of ECM. Mass spectrometry (MS)-calibrated measurements of heart collagen content (*µ*g per heart) suggested power-law scaling of tissue stiffness values with collagen-I levels as shown in Fig. 4A. The strain-free configuration of the heart tissue obtained by inhibiting tissue contractility or beating (via blocking myosin activity with blebistatin) resulted in a loss of tissue collagen-I levels (as well as stiffness values as shown in Fig. 4B). MS-measurements showed the presence of collagen-I and endogeneous collagenases (MMP-2, -15) within the heart tissues. The loss of collagen within non-beating heart tissues appears to result from the degradation by endogenous collagenases. Here, it is important to mention that the addition of exogeneous BC to beating heart samples resulted in a loss of heart tissue collagen-I level in strain-dependent manner, duly verified by degraded collagen-I fragments with coomassie brilliant blue gel assay (Fig. S22). Interstingly, the blocking of endogeneous collagenase activity by pan-MMP inhibitor (GM6001) was observed to rescue tissue collagen degradation within the strain-free heart tissue samples (Fig. 4C).

**Fig. 4.**
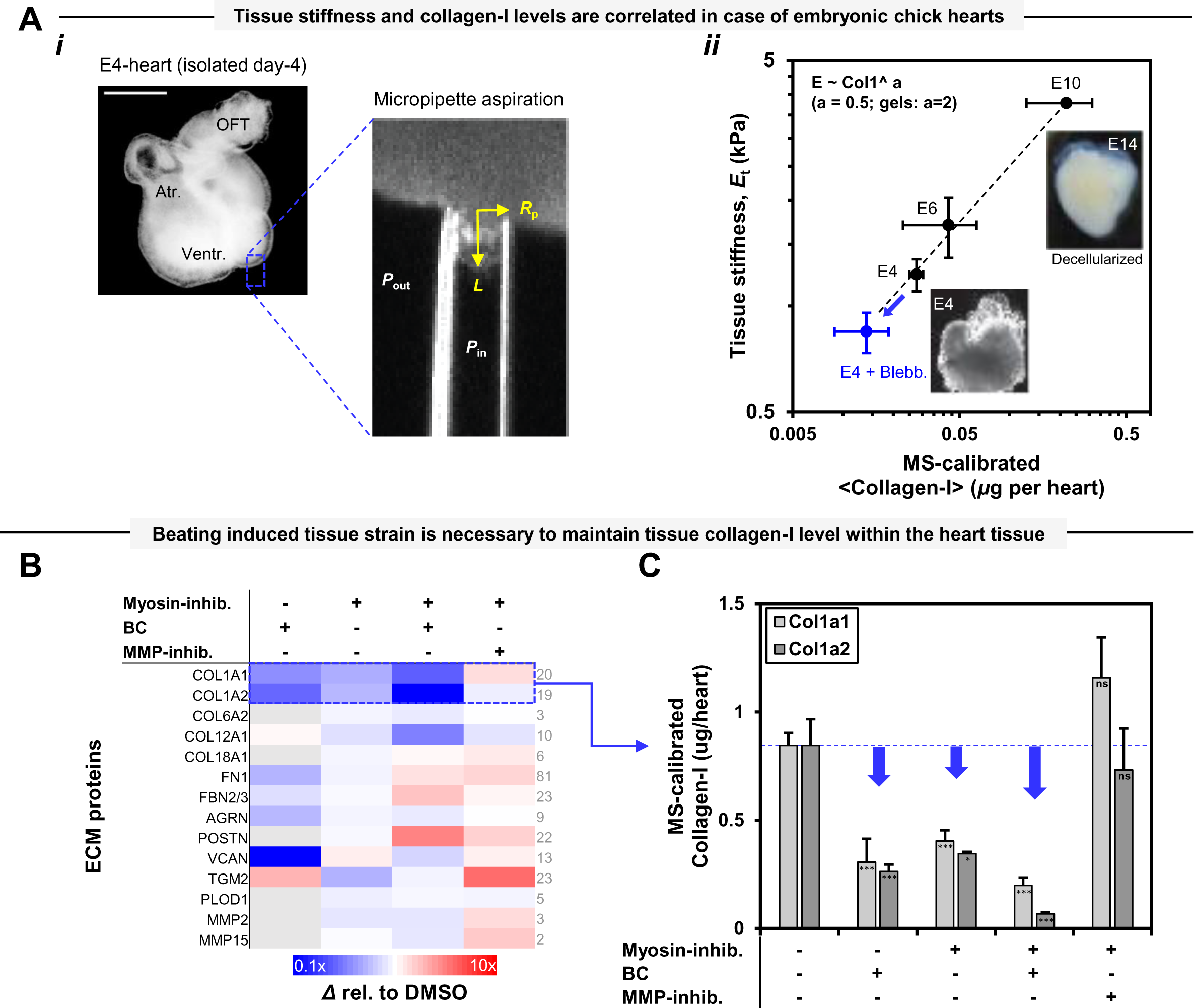
The inhibition of contractility of ex-vivo beating embryonic chick hearts increases collagen degradation rate by MMPs while the inhibition of MMP activity rescues the degradation of collagen. **A. (*i*)** Image of an embryonic day-4 (E4) chick heart. ‘OFT’: Outflow tract, ‘Atr’: Atrium, ‘Ventr’: Ventricle (scale bar = 500 µm). Right inset: Tissue stiffness, *E_t_*, measured by micropipette aspiration. **(*ii*)** Stiffness *E_t_* of developing hearts (E4, E6, E10; n=8,4,2 hearts per lysate, respectively) shows power-law scaling versus mass spectrometry (MS)-calibrated measurements of collagen concentration (*µ*g per heart) and suggest collagen as a major determinant of tissue mechanics (for details, refer to Fig. S21B). Error bars indicate ±SEM. **B & C.** MS-generated heat-map of all detected ECM proteins in E4 hearts after treatment with Blebbistatin, collagenase (BC), or both (n=8 hearts per condition). Heat-map illustrates fold change relative to DMSO control. Myosin-II inhibition (by blebbistatin) and ECM-softening (by BC) both result in reduction of signal for many ECM proteins, particularly collagen-I subunits COL1A1 & 2. The addition of pan-MMP inhibitor ‘GM6001’ rescues the effects of blebbistatin on many ECM proteins.

### 3.5 Mechanical strain-dependent degradation of tissue collagen by collagenase is correlated to unfolded triple-helical collagen content

In order to ascertain mechanical strain-dependent changes in collagen molecular structure and collagen degradation rate, the degradation rates of TGM cross-linked fascicle sub-lengths that were uniaxial deformed and exposed to BC, were investiagted. Here, strained TGM cross-linked fascicle sub-lengths were compared to strain-free counterparts (without TGM cross-linking) where collagen degradation rate is masured based on forward SHG signal while collagen triple-helix unfolding with the tissue was quantified using CHP labelling (Fig. 5). TGM is likely to form amide cross-links between Glutamines and Lysine residues (present at several sites of alpha-I and alpha-II chains, Fig. S23A) of neighbouring collagen molecules (Orban, Wilson et al. 2004). TGM induced cross-links are likely to induce increased collagen triple-helix unfolding within mechanically deformed TGM cross-linked samples as compared to strain-free control samples (i.e. without TGM treatment) (Fig. 5 (*i*). We observed a sigmodical increase in both collagen degradation rate and level of unfolded triple-helical unfolding level and in mechanically deformed TGM-cross-linked samples (Fig. 5 (*ii-iii*)). The increase in the level of unfolded triple-helix collagen content and collagen degradation rate of fascicle samples was observed up to ∼3 times to that of strain-free control samples. Importantly, collagen degradation rate values and unfolded triple-helical collagen content levels were positvely correlated (Fig. 5 (*iv*)). Further, the local changes in strain and instantaneous degradation rate with time in case of TGM cross-linked samples (as well as external strain-free controls) that are subjected to mechanical deformation and exposed to BC, have been shown in Fig. S24(*i*-*ii*). While local differencees in strain magnitudes of different regions within each sample sub-length were observed despite uniaxial deformation (Fig. S24 (*i*), the strain magnitudes did not alter during degradation by BC at least during the initial phase of tissue degradation.

**Fig. 5.**
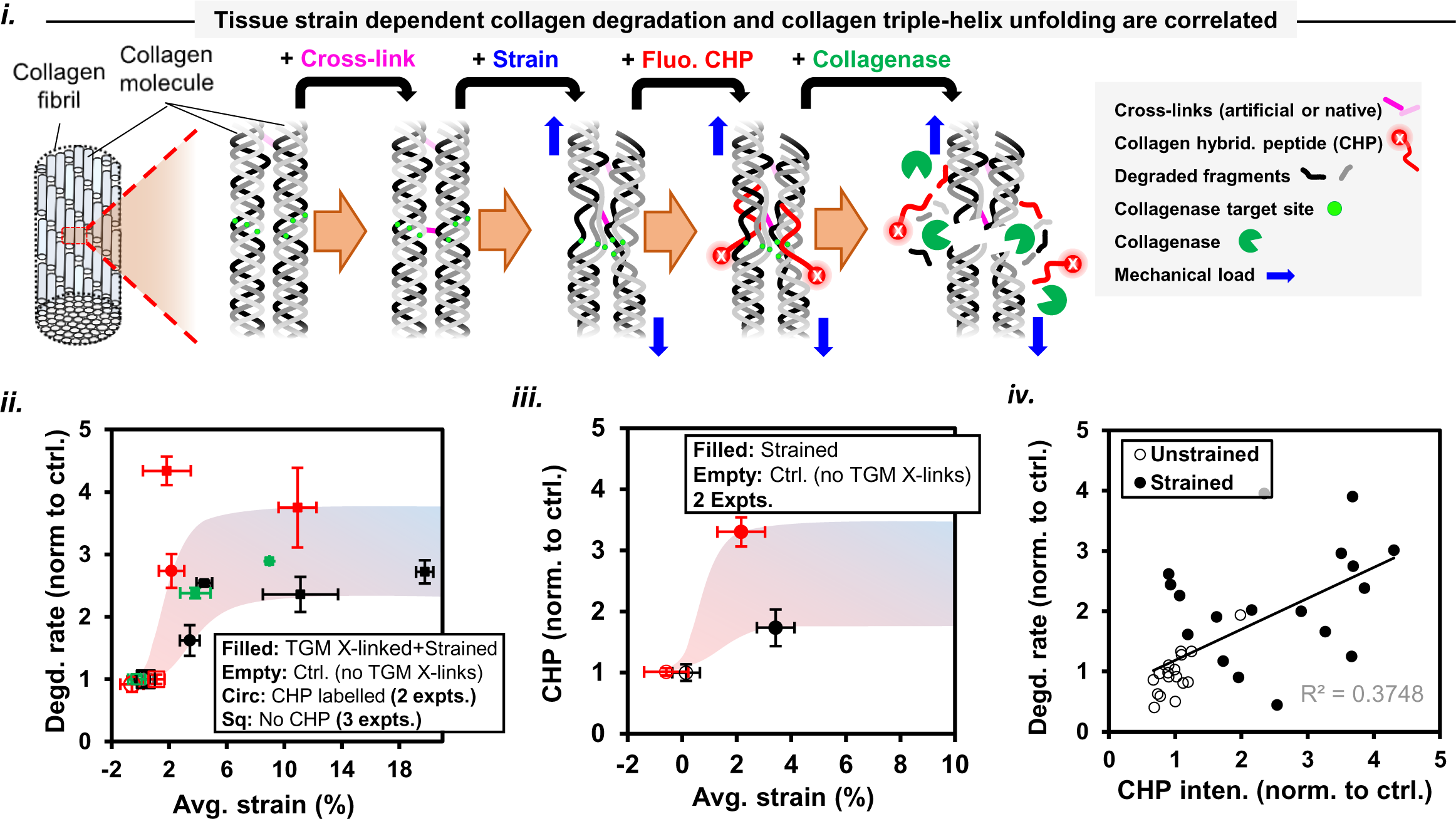
Mechanical strain dependent degradation of exogeneous enzyme cross-linked tissue collagen by BC is determined by unfolded triple-helix collagen content within the tissue. **(*i*)** Schematic showing collagen degradation rate at increasing tissue strain values follow a sigmodal increase (as compared strain-free control samples). **(*ii*)** Sigmoidal increase in collagen triple-helix unfolding levels at larger strain magnitudes (normalized with respect to strain-free control samples) as reflected by the binding of fluorescent collagen hybridizing peptide (CHP). **(*iii*)** Tissue collagen degradation rate and collagen triple-helix unfolding levels are positively correlated.

## 4. Discussion

The cell-free version of tendon fascicles obtained using freeze-thaw cycles (Gilbert, Sellaro et al. 2006) (Burk, Erbe et al. 2014) (Roth, Glauche et al. 2017) can be assumed to carry intact ECM collagen that is free of endogeneous collagenases, protein synthesis. Novel low-cost bioreactor design is best suited for local tissue deformation studies at size-scale of cells while providing information of tissue collagen level changes in a high-throughput manner. Thus, the bioreactor design overcomes the limitations posed by the conventional mechanical testing techniques including accuracy of tissue strain quantification, sample holding, installation on micro-scope and extent of information extracted from samples. Given high frequency of molecular vibrations and interactions resulting from thermal motion, the static deformation of a “cell-free” fascicle sample can be assumed to accurately capture tissue strain-dependent collagen-collagenase molecular interaction behavior despite the tendon tissue sustaining cyclic loads physiologically. SHG signal loss during degradation represents a loss of collagen triple-helix conformation within collagen fibrils as evident in case of heat denatured collagen-I from bovine achilles tendon (Theodossiou, Rapti et al. 2002). Fluorescent tag based tissue collagen quantification during imaging is likely to be limited by several factors including photobleaching during image acquisition, quenching of fluorophores, non-specific binding of fluorophore to non-collageneous proteins within tissue, strain-dependent affinity of fluorophores. Hence, SHG signal based tissue collagen degradation rate measurements can be considered more accurate than that of conventional fluorescent tag based measurments. In general, SHG signal attenuation within thicker tissues limits the measurment of forward SHG signal. The novel design of the bioreactor together with the size of fascicle (up to ∼200 *μ*m diameter) facilitates simultaneous imaging in both forward and backward directions. In contrast to BC based degradation experiments, the samples exposed to MMP-1 were kept in the incubator for ∼3 hours during tissue degradation experiments to exclude the influence of evaporation, temperature sensitivity of MMP-1 activity. The incubation of samples during degradation experiments involving MMP-1 produced in lesser time-points for curve fitting than that of involving BC towards slope-based collagen degradation rate measurements. Our calculations based on assumption of linearity of collagen content degradation with time is duly supported by previous report involving collagen gel degradation by MMP-1 (Welgus, Jeffrey et al. 1980) (Welgus, Jeffrey et al. 1981). Undoubtedly, the provision for mechanical manipulation within the bioreactor and dipping nature of lens of upright microscope together limits the sealing and results in reaction buffer evaporation from the edges of the bioreactor.

Changes in reaction buffer composition due to evaporation might affect collagen-collagenase interactions by influencing local solute concentration and/or collagen fibril molecular packing. Making up of evaporation losses in case of longer duration of experiments, use of internal controls towards data normalization, excluded such influences on our observations. The local strain measurements within each fascicle sample were made for ∼50 um size regions (along fascicle-axis), which is slighly larger than the size of fibroblast (∼15 um), closely represents the size-scale of cell micro-environment. Our local tissue strain based measurments are uninfluenced by limited repeatability of strain distribution among different fascicles samples due to the use of internal controls, data normalizations. The use of pattern-photobleach and Gaussian peak based photobleach stripe position prediction within the sample, while maintaining same photobleach stripe width, stripe-stripe spacing as well as number of data points for Gauss curve-fitting, further increases the accuracy and repeatability of the measurements. Pattern photobleaching of fascicle at a single plane (at ∼ half-diameter of a fascicle) instead of photobleaching at multiple planes through the tissue depth ensured investigation of same sample location with respect to microscope lens upon tissue deformation and enhanced accuracy. The use of fluorophores on samples for strain quantification purposes is likely to interfere with collagenase-collagen interactions but the selected fluorophores did not significanly interfered with the sample degradation (Fig. S5C).

As the initial phases of tissue collagen degradation are almost free of local tissue strain-relaxation (Fig. 2B), the SHG based tissue degradation rate comparison based on initial tissue strain sounds rationale. Our observation of MMP-1 and BC showing a large difference towards soluble monomeric collagen-I cleavage rate is in agreement with a previous report (Garnero, Borel et al. 1998). The difference in MMP-1 and BC activity on monomeric collagen is presumably a result of the number of collagen molecule cleavage site(s), difference in the cleavage mechanism and/or target sites on collagen molecule. The similarity of strain magnitude dependent suppression of tendon fascicle collagen degradation rate by MMP-1 and BC within physiological strain magnitudes (∼5-8% strain for tendon) appears to result from strain-dependent tissue collagen fibrillar architecture changes, and might also apply to other collagenases. In case of collagen fibrils, the most abundant collagen form within tissues, the packing of collagen molecules within a fibril exposes only “fibril-surface” collagen molecules to cleavage by collagenase (Welgus, Jeffrey et al. 1980) (Welgus, Jeffrey et al. 1981). Mechanical deformation of tendon tissue has been reported to affect constituting collagen fibril architecture in terms of its D-banding, triple-helix unfolding, etc. (Peter and Richard 2007) (Marc André, Po-Yu et al. 2008) (Fratzl, Misof et al. 1998) (Zitnay, Li et al. 2017), and suggests physiological strain values (up to ∼5-8% strain) within tendon tissue cause reversible structural changes while larger strains are likely to cause permanent tissue damage at molecular level. The similar strain-dependent suppression of collagen degradation rate by MMP-1 and BC within the physiological strain limits, despite different cleavage mechanism of collagenases, demonstares tight inter-relationship between strain and tissue collagen degradation. At larger strain values (beyond ∼8% strain), the noisy nature of strain-dependent collagen degradation appears as an outcome of tissue micro-fracture, molecular unfolding, etc. (Fratzl, Misof et al. 1998) (Zitnay, Li et al. 2017). Taking together the data for different strain magnitude regimes (Fig. 1E), the strain-dependent degradation appears to be the result of strain-dependent biochemical regulation and micro-fracture within the tissue, and suggests physiological mechanical strains are optimal in minimizing tissue collagen degradation without causing considerable tissue damage.

The observed strain-suppressed degradation rate for fasciles exposed to BC while deformed in different modes i.e. heterogeneous and uniaxial (Fig. 1A and Fig. 2A), suggests the strain-suppression of degradation rate as an intrinsic material property of tissue collagen. As collagen fibrils are oriented along collagen fibers within a fascicle, the strain-dependent collagen-fiber organization changes (in terms of waviness, orientation values) during degradation by BC suggest a strain dependence of tissue collagen fibril life-time (Fig. 2). The strain magnitudes sustained by tissue collagen fibrils might differ from micro-scale tissue strains as the exact relationship between tissue at two length-scales remains unknown to date. As shown in Fig. 2B-2D, the spatial difference in strain values, instantaneous collagen degradation rate and collagen fiber organization within fascicle ROIs indicate local tissue material heterogeneity experienced by the cells. The observed high Pearson’s correlation coeficient (*r*) values for parameter (i.e. strain values, instantaneous collagen degradation rate and collagen fiber organization) values with time (shown in Fig. 2E) suggests interdependence of these parameters. The observed positive or negative *r* - values for ROIs within sample fascicle is likely to result from residual internal stress relaxation as a result of tissue collagen degradation. The quantification of the increase in waviness or orientation changes with time (during degradation by BC) within strained sub-length of fascicles suggest ∼5% tissue strain as a critical value towards determining the nature of strain relaxation (compressive or tensile). Based on the difference in strain magnitudes sustained by inclined and aligned collagen fibers within a tissue, the outcome as relatively larger number of inclined fibers following degradation by collagenase (with respect to to aligned ones shown as schematic in Fig. S13B) within a strained sub-length might have been resulting from either strain-stablization of inclined collagen fibers (as compared to that of aligned ones) or stress-relaxation within aligned collagen fibers. Hence, the local changes in collagen content and collagen fiber organization appear to be determined by the nature of tissue strain i.e. physiological or non-physiological.

The observed heterogeneity in the local behavior of strained fascicles (Fig. 2B-D) with collagen degradation despite collagen fiber parallelism and orientation along fascicle-axis, indicates a locally different in material response. Despite putative global affine observation of collagen fiber deformation behavior (averged over ∼500 *μ*m fascicle length) within our experiments, the primary collagen fiber inclination plays key role in dtermining local fascicle deformation behavior. The dependence of the local fascicle deformation behavior (∼50 *μ*m fascicle length) on the primary collagen fiber inclination (Fig. 3A-ii) at the size-scale of the cell emphasizes the importance of local collagen orientation in controlling strain-dependent tissue collagen fiber reorganization (Fig. 3A). Undeformed fascicles carry crimps with a pitch of ∼140 *μ*m natively (Legerlotz, Dorn et al. 2014). We observed “crimp-waves” to be out of phase in the undeformed fascicles when seen in transverse direction to loading. Upon deformation of the tissue, crimps underwent reorientation to result aligned collagen fiber along fascicle-axis. The loss of “crimp-waves” phase to result collagen fibers parallel to fascicle-axis upon application of mechanical deformation suggest a local shear among collagen fibers as load transfer mechanism within fascicles as reported elsewhere (Szczesny and Elliott 2014). In addition to local collagen fiber orientation distribution within undeformed fascicles, the local strain distribution within a fascicle might have been influenced by other factors such as non-collagenous protein distribution, applied load magnitudes (physiological or non-physiological), spatial distribution of cells, pitch of crimps, etc., (Legerlotz, Dorn et al. 2014) (Zuskov, Freedman et al. 2020) which further depend on age and condition (i.e. physiological or pathological) of a tissue.

The insight into the possible mechanisms of strain suppressed degradation of the tendon fascicle made in terms strain-dependent changes in solute permeation, tissue mass density and mobilty of collagenase-size molecules further emphasize the significance of physiological strain within tissues. The size of freely diffusing solute particle in a solution in terms of Stoke’s radius (as compared to M.W.) was used as a reference for our measuments keeping in mind the fact that the Stoke’s radii for dextran molecules is larger than similar M.W. protein molecules (Tyn and Gusek 1990). The solute permeation was not significantly influenced within physiological strain limits for the tissue. The fluctuations in collagen concentration and VR with strain indicate collagen fiber reroganization (in agreement with non-affine deformation behavior of collagen fibers within fascicle) was evident but the tissue mass density changed insignificantly (remains within +/-10%) for physiological strain limits of the tissue. Although, fascicle collagen fibers have been assumed to lie in the image plane i.e. perpendicular to the direction of light propagation in our analysis, the observed strain-dependent fluctuations in tissue collagen concentration might have been influenced by the tissue strain-dependent reorientation of collagen fibers as a result of the collagen fiber movement (into/out of the image plane) at least within physiological strain limits. The strain-dependent tissue volume fluctuations in a heterogenously deformed fascicle are in agreement with that of observed in the case of uniaxially deformed fascicles. The influence of strain-dependent tissue structure changes on local molecular kinetics of various size dextran molecules (Stoke’s radii of 1.4 nm, 8.5 nm and 16 nm) and fluorescent-BC show dependence on solute-size. Commercially available BC is a cocktail of several M.W. molecules (60-130 kDa) while M.W. of most MMPs and Albumin is around ∼60 kDa. Because Albumin has a Stoke’s radius of ∼3 nm (Tyn and Gusek 1990), the use of fluorescent-BC in our FRAP experiments can be assumed to precisely capture general mobile behavior of MMPs in the tissues. The increase in mobile fraction values with increase in the size of Dextran molecules appears counter-intuitive (Fig. S20A), and might be resulting due to several factors including different shape of molecules (smaller Dextran molecules are rod-like while symmetry increases with size due to branching), affinity of hydrophobic rhodamine in TRITC-tag (James W 1996) in dextran molecules towards ECM as well as different extent of fluorescent labelling, etc. Regardless of above mentioned factors, our tissue strain-dependent measurments provides a clear insight into the relative changes based on an internal control. Both fluorescent-BC (∼3 nm) and collagenase size dextran (1.4 nm) molecules show a sudden increase in the mobility values at strains larger than physiological strain limit of ∼8% of tendon fascicles that are likely to result from the local tissue fracture and collagen molecular unfolding (Fig. S21) (Zitnay, Li et al. 2017). The lower mobility of fluorescent-BC than that of similar size hydrophillic Dextran molecules might be resulting from the affinity of BC molecules towards ECM collagen. Importantly, no significant change in mobility of fluorescent-BC molecules has been observed for fascicle samples strained within the physiological limits. The collective observation of solute permeation, tissue mass density and mobility within physiological tissue strain limits show strain independence but suggest an alternate mechanism for strain suppressed degradation of tissue collagen that is likely to depend on “conformational-access”.

In agreement with strain-suppressed collagen degradation in a “cell-free” fascicle tissue exposed to exogeneous collagenase(s), our evaluation of live tissue samples, carrying a cocktail of many MMPs, TIMP-2 (Brauer and Cai 2002) that act together to maintain tissue collagen levels in strain-dependent manner, further supports our hypothesis. Collagen-I level dependent stiffness of E4 beating chick hearts, which is optimal for beating by heart cells (Majkut, Idema et al. 2013) (Chiou, Rocks et al. 2016) together with the presence of endogeneous MMP-2 as collagenase (Aimes and Quigley 1995), makes the heart tissue a suitable candidate for our investigation. We modulated the heart tissue strain by inhibiting the tissue beating through the use of molecular inhibitor of acto-myosin contractility. Compared to almost constant collagen-I level in beating E4 chick hearts, the decrease in collagen-I within an hour of non-beating (strain-free) indicates strain dependence of tissue collagen degradation by endogeneous MMPs. Further, the inhibition of endogeneous MMPs activity by a pan-MMP inhibitor was observed to rescue collagen degradation in case of strain-free ECM. Analogous to strain-suppressed degradation of tissue collagen as observed in the case of fascicles exposed to exogeneous collagenases, the addition of exogeneous BC to E4-beating chick hearts degrades tissue collagen-I within ∼1 hour but the extent of collagen-I degradation is more pronounced in the case of strain-free chick heart samples. It is important to note that the observed strain-suppressed degradation of collagen-I within the E4 beating chick heart tissues was unaffected when protein synthesis was blocked by cycloheximide, and suggest strain suppressed degradative sculpting of ECM collagen within the tissue.

TGM molecules, forming amide cross-links between Glutamine (*Q*) and Lysine (*K*) (*Q*s or *K*s form less than 1% of total number of residues within triple helical region of a collagen molecule, Fig. S23-A) residues of neighboring collagen molecules, result molecular level structural alterations within a fibril (Fig. S23). While native Lysyl oxidase induced cross-links are distributed throughout the fibril, ∼38 kDa TGM induced collagen cross-links are likely to be present on the surface of collagen fibrils present in a tissue. Given the sensitivity of SHG signal to collagen fibril packing, the change in SHG signal in case of strain-free (TGM cross-linked and without TGM cross-linking, Fig. S23 B-*i*) and strained TGM cross-linked fascicle samples (Fig. S23 B-*ii*) is most likely to result from an altered inter-molecular spacing collagen molecules within a collagen fibril (Lutz, Sattler et al. 2012). At lower tissue strain magnitudes including strain-free configuration, TGM cross-links are likely to suppress tissue collagen degradation by restricting collagenase dependent collagen triple helix unfolding in agreement with previous reports involving Ribose based collagen cross-linking (Bourne, Lippell et al. 2014). Mechanical deformation of tendon fascicle has been reported to induce inter-molecule silding among collagen molecules within a fibril (Peter and Richard 2007) (Marc André, Po-Yu et al. 2008) (Fratzl, Misof et al. 1998), the constraints imposed by the presence of TGM cross-links are likely to result increased triple-helix unfolding at larger tissues strains (Zitnay, Li et al. 2017) (Bourne and Torzilli 2011). As the degradation of unfolded collagen triple-helices is energetically favourable than that of folded collagen present in abundance within different tissues, the increase in collagen degradation rate at larger strain magnitudes is most likely mediated through exposure of unfolded collagen strands to collagenases. Depending on the tissue strain magnitude, the rate of collagen degradation within TGM cross-linked mechanically deformed samples is either faster or slower than the strain-free tissue samples.

Based on our observations of insignificant changes in solute permeation, mobility or tissue mass density with tissue strain while considering the collagen triple-helix unfolding in the case of TGM cross-linked fascicle samples and the mechanosensitive nature of the collagen at molecular or fibril (Camp, Liles et al. 2011) (Flynn, Bhole et al. 2010) (Saini, Cho et al. 2019), we propose that mechanical strain suppresses the degradation by restricting the conformational access of tissue collagen cleavage sites to collagenase molecules. Tensile loading of tendon tissue has been reported to influence the structure of collagen fibrils (Misof, Rapp et al. 1997) (Folkhard, Mosler et al. 1987) as indicated by the changes in characteristic D-banding upon straightening of macroscopic collagen fiber crimps (Fratzl, Misof et al. 1998) (Diamant, Keller et al. 1972). Although, the direct observation of tissue strain-dependent collagen molecular conformational changes is still lacking, our observation of strain-dependent collagen triple-helix unfolding in the case of TGM cross-linked fascicles deformed within the physiological strain limits and collagen triple-helix unfolding in fascicles (without artificial cross-links) at strains larger than the physiological limits clearly support strain-dependent molecular conformation changes in colllagen molecules constuting a collagen fibril. Circadian rhythms are also attributed to the change in tissue collagen level in terms of small diameter non-covalently cross-linked and non-load bearing collagen fibrils within mice tendon every 24 hours (Chang, Garva et al. 2020) which motivates the need to further understand how different size collagen fibrils contribute towards load transfer within a tissue.

## Author Contributions

K. Saini, S. Cho, D. Discher designed experiments; J. Irianto, J. Andrechak, L. Dooling., C. Alvey, A. Kasznel, D. Chenoweth, K. Yamamoto contributed key materials; K. Saini, S. Cho, M. Vashisth, A. Jalil performed and analyzed experiments; K. Saini wrote, K. Saini, L. Dooling and D. Discher revised and finalized the manuscript.

## Acknowledgments

The authors in this study were supported by the National Institutes of Health National Cancer Institute under Physical Sciences Oncology Center Award U54 CA193417, Human Frontiers Sciences Program grant RGP0024, National Heart, Lung, and Blood Institute award R01 HL124106, SERB Indo-US Postdoctoral Fellowship 2016-157, the US–Israel Binational Science Foundation, National Science Foundation, Materials Science and Engineering Center grant to the University of Pennsylvania and also grant agreement CMMI 15-48571. This content is solely the responsibility of the authors and does not necessarily represent the official views of the National Institutes of Health or other granting agencies. Leica SP8-MP upright microscope at Penn Vet imaging core please also reference NIH grant S10 OD021633-01

## Supplementary Information

### S1. Mechanical manipulation bioreactor design, and installation on MP microscope

The bioreactor has glass coverslips of thickness 130–160 *μ*m (Fisherbrand™ Cover Glasses) as upper plate and middle plate that are glued to each other and bottom polystyrene surface of petri-dish using nail-paint (shown in Fig. S2). Together with thickness of nail-paint film (∼40 *μ*m) and side glass plates, a height of ∼220 *μ*m is obtained between bottom polystyrene surface and upper-plate that provides a sufficient room to guide the motion of movable glass plate for nearly in-plane mechanical deformation of fascicles for sizes up to ∼ 220 *μ*m. The sharp movable glass plates of slightly larger thickness (160–190 *μ*m) were obtained by manually breaking the glass coverslips (VWR 48366-205) to the desired corner-shape for subsequent manual sliding into the space between upper glass plate and bottom polystyrene surface for heterogeneous deformation without rolling of the sample. The bioreactor with petri-dish as bottom plate was used in all experiments to maintain the desired reaction buffer around samples during the experiments and petri-dish was kept on heating plate of the micro-scope as shown in Fig. S2 for imaging purposes. In order to place the sample in the bioreactor, wet fluorescently labelled fascicle (kept at 4 deg. C) were taken out of the Eppendorf tube using sharp tweezers. The fascicle samples were gently put on a larger size PBS drop lying on bottom polystyrene surface that was carefully soaked to reduce drop’s volume facilitating minimum residual strains as well as air-drying of the sample. The sample adherence to the hydrophobic polystyrene surface was evident from the absence of sample floating when PBS was immediately added to maintain sample hydration whereas the presence of strain-free configuration of sample on polystyrene surface was duly confirmed by SHG image of the sample. Although, sufficient care was taken to minimize the dehydration of the samples at each step of the experiment, the samples undergoing complete dehydration followed by rehydration in PBS (at least for 30 minutes) were found to be degraded by collagenase (BC or MMP-1). The presence of the crimps within a sample via SHG image confirmed the strain-free configuration of the sample adhering to polystyrene surface after the transfer from Eppendorf tube to bioreactor (otherwise, this step wise repeated). It must be noted that during the sliding of movable glass-plate on bottom polystyrene surface in order to produce local strain gradients within the sample, the sample must be surrounded by minimum volume of the medium so that only interaction between wet sample and dry polystyrene surface may take place with subsequent addition of sufficient PBS to maintain sample hydration. In addition to local adhesion forces between sample-polystyrene surfaces in the fascicle regions locally pushed by movable corner, the larger longitudinal dimensions of the sample (with respect to fascicle diameter) facilitate sample local deformation along fascicle-axis (similar to the fascicles whose ends are glued). Although, the adhesion between fascicle and polystyrene surface together with the forces along fascicle longitudinal direction (exerted due to movable corner or sample-end conditions) was observed to be sufficient to retain the deformed configuration of a fascicle sample at least during initial phase of the degradation, the replacement of the bottom polystyrene surface with glass or other surface(s) may be used to tune the magnitude of adhesive forces exerted on the sample by bottom surface.

### S2. Strain quantification of fluorescently labelled fascicle samples

Laser settings for pattern photobleach on fascicle samples: Leica SP8 Confocal/Multiphoton Upright Microscope having 20X water immersion lens (of 1.0 NA) and Leica Application Suite (LAS) software with FRAP wizard was used to create a photobleach pattern on the fluorescently labelled fascicles using a non-polarized 20 mW laser beam emitted from the resonator passed through focusing optics. Keeping in mind the sample curvature, local tissue heterogeneity, fluorescent intensity variation due to sample depth, etc., an optimal z-position based on higher fluorescent intensity as well as maximum area on the sample (at a depth of ∼half diameter) was selected for creating a photobleach pattern with the following settings: line freq.= 200 Hz; exposure time = 5.24 sec. per image; intensity of laser for photobleached/non-photobleached regions = 90/0.1%; no. of iterations = 180. The pattern created on a sample had ten evenly spaced rectangular stripes of ∼5 *μ*m width (covering sample diameter) and 55 *μ*m spacing between them (center to center) as shown in Fig. S4B-i. Based on the size of image i.e. 553 *μ*m x 553 *μ*m, the photobleached region results a total area of ∼7500 *μ*m^2^ (= 5×150×10 for a 150 *μ*m diameter fascicle) that covers approximately 2.45% of total area of the image where the laser stays at high intensity for 0.128 seconds (2.45% of 5.24 sec.) during each iteration. The total fluence for photobleached region is computed as laser power × exposure time/square area, which comes out as 0.307 x 10^-7^ J/mm^2^ for each iteration and a total fluence of 55.29 J/mm^2^ for 180 iterations performed in an experiment, seems a reasonable choice towards good photobleached pattern contrast and minimum sample damage due to laser exposure (Jayyosi, Fargier et al. 2014).

Strain quantification by Gaussian fitting: In order to make the local displacement measurements between each pair of consecutive photobleach stripes on the sample in undeformed and deformed configurations while keeping in mind local sample heterogeneity, we fitted Gaussian curves to a ROI path (i.e. yellow line covering two-consecutive stripes in Fig. S4B-*i*) selected in a direction transverse to the photobleach stripe length (Lee, Wong et al. 1999). Here, a single composite image was obtained by summing up fluorescent intensities of all images within a stack covering fascicle dimensions diametrically along laser propagation direction. Using ImageJ package, a ROI path on the composite image was created to capture fluorescent intensity magnitude as function of length along fascicle-axis direction where the fluorescent intensity was transversely averaged (parallel to the stripe-length) over certain distances the two neighboring stripes were observed parallel to the selected stripe within a composite image. For fitting a Gaussian curve to each valley of a ROI path carrying several photobleach stripes, a custom MATLAB script was used that automatically selected a point of minimum intensity near each photobleach stripe and a certain number of points on either side of the photobleach stripe. The changes in local length of ROI path (on both sides of each photobleach stripe) affecting the number of points selected for Gaussian fit was found to influence the obtained stripe (based on Gaussian) width as shown in Fig. S4D due to local sample heterogeneity where the Gaussian peak position was observed uninfluenced. Therefore, we selected fixed local ROI length (∼ 25 *μ*m) as analysis length along with good R-square values (> 0.90) for all Gaussian peak position-based measurements. Based on the mentioned criteria, Gaussian peaks were used to calculate the displacement between each pair of consecutive photobleach stripes (Fig. S4B-ii) in undeformed and deformed configurations (Fig. S4C-*i*). On the basis of the displacement values between each pair of consecutive photobleach stripes in undeformed and deformed configurations, the average value of strain within a region lying between parallel stripes were estimated for different regions of a deformed fascicle (Fig. S5A-*i*).

### S3. SHG signal for collagen content measurement with tendon fascicle and comparison with fluorescent signal

Collagen content of a fascicle (within the photobleached region) was obtained by creating a projected sum composite image from the stack(s) (in forward and backward directions) of SHG images covering whole fascicle sample diametrically while keeping constant slice spacing of 1 or 2 *μ*m within an experiment). Once collagenase was added to the reaction buffer carrying deformed fascicle, SHG image-stacks from the photobleached region were captured continuously (using automated control of the micro-scope and keeping same settings for laser and detectors) to track sample degradation dynamics at different time-points of the experiment. An upright multi-photon microscope TCS SP8 MP (Leica) was utilized to image fibrillar collagen content within the tendon fascicles. The tunable Coherent Chameleon Vision II Ti:Sapphire laser (680-1080 nm) produced linearly polarized pulses spectrally centered at 910 nm of 1.99 W power. Incident light at 910 nm was focused onto the sample with a 20X water immersion HCX APO L (1.00 NA) objective. Due to collagen fibril size dependent emissivity (Lacomb, Nadiarnykh et al. 2008), the signal emitted in forward direction being collected by a 0.9 NA condenser lens whereas in backward direction by a highly sensitive HyD detector. The SHG laser power was adjusted to 3%/Gain:90.0/Offset:51.9 so that sufficient SHG signal was obtained without any obvious damage to the sample as reported elsewhere (Williams, Shear et al. 1999) (Williams, Zipfel et al. 2005). Each z-stack acquired the whole volume of the fascicle sample with 200 Hz line frequency (having 2048 × 2048 pixels) using LAS X (Leica Application Suite X). The efficiency of SHG signal generation by collagen fibrils increases at the shorter wavelength but also result reduced cell viability and decreased effective imaging depth as result of more scattering (Zipfel, Williams et al. 2003) (Williams, Shear et al. 1999), hence, we selected ∼900–910 nm wavelength for all of our experiments to probe deeper sample depths without loss of SHG intensity in forward and backward directions.

### S4. Comparison of MMP1 and BC activity on soluble collagen

The comparison of MMP1 and BC activity on soluble collagen was made using ELISA experiments where normalized intensity change (with respect to negative control) was used for degradation rate quantification. The normalized intensity was obtained using the following

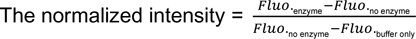

Degradation rate (*μ*g/min) = A * B * C

where, A = slope of normalized intensity change (with respect to to negative control) versus time (in minutes); B = Final collagen conc. (*μ*g/mL) in a reaction well; C = Volume of solution in the well (in mL). As shown in Fig. S6A, BC cleaves soluble collagen molecules almost ∼50 times faster than MMP-1 but the observation of catalytic activity of MMP-1 and BC (at different concentrations shown in Fig. SB-*ii*) on soluble collagen also indicates that the catalytic activity of MMP-1 is sensitive to the presence of BSA (0.01% solution) while BC activity remains almost unaffected by BSA molecules.

### S5. Influence of reaction buffer composition changes on molecular packing within collagen fibrils

The composition of a reaction buffer in terms of the concentrations of various salts or solutes might change with time as a result of evaporation losses from the bioreactor that in turn may influence reaction buffer pH values or packing of collagen molecules within collagen fibrils quantified by means of backward/forward SHG signal. Fig. S7-*i* indicates no significant change in pH value of PBS (as a reaction buffer) due to solution volume change due to evaporation. The make-up for evaporation losses from the bioreactor has been made by adding DI water based on the evaporation rate of various solutions .i.e. PBS, PBS + dextran, PBS+BC as shown in Fig. S7-*ii* where the presence of different solutes were observed to influence the evaporation rate of the medium.

Although, the make-up for evaporation losses from the bioreactor was made every ∼2 hours by adding sufficient volume of DI water to the medium based on the evaporation rate of the medium for respective composition, the changes in forward and backward SHG signal (representing larger and smaller diameter collagen fibrils) at various solute concentrations were used to quantify the solute concentration dependent changes in the molecular packing of collagen fibrils within a fascicle tissue. As shown in Fig. S8A, the changes in various dextran concentrations (i.e. 0.11, 0.20 and 0.62 mg/mL) were not observed to influence molecular packing within collagen fibrils. The presence of CaCl_2_ (0, 5, 25 mM) or NaCl (0, 300 mM) around the fascicle samples (shown in Fig. S8B and Fig. S8C) was observed to promote the disassembly of collagen fibrils within the tissue as suggested by the decrease in F/B ratio values (obtained from the magnitudes of absolute forward and backward SHG signal) from different ellipsoidal regions of a fascicle sample with longitudinal limits defined by photobleach stripes. The superposition of the mechanical deformation on the fascicle in addition to the presence of the salts was observed to further decrease the F/B ratio value, suggesting further disassembly of collagen fibrils present within the tissue.

### S6. Quantitation of local collagen fiber orientation distribution and deformation behavior

Local collagen fiber orientation within a fascicle sample indicated by SHG signal can quantified by means of structure tensor (Rezakhaniha, Agianniotis et al. 2012) (Avila and Bueno 2015). Although tendon fascicles carry bundles of parallel collagen fibers, there are native crimps where these fibers show an inclination to fascicle-axis. Interestingly, collagen fiber orientation also changes when moving in the transverse direction of fascicle (i.e. diametrically as shown in Fig. S10A-*i*), and upon deformation of the fascicle, collagen fibers tend to align along fascicle-axis (Fig. S10A-*ii*). The local collagen fiber orientation (averaged over voxel of size 1-2 *μ*m^3^) was used to predict local collagen fiber orientation distribution within each ROI (i.e. fascicle tissue region lying between a pair of consecutive stripes) with respect to fascicle-axis as shown in Fig. S10B where the fascicle-axis orientation for each ROI represents the average inclination of the ROI edges obtained from the composite image (sum projected image of respective forward SHG stack). Using the known collagen fiber orientation distribution (with respect to fascicle-axis) within a ROI of undeformed fascicle, the affine predicted collagen fiber orientation distribution for respective deformed configuration can be estimated based on local tissue strain magnitudes (see methods: sub-section 2.2.5). As listed below, the comparison between affine predicted and experimentally observed collagen fiber orientation distribution in deformed configuration of a ROI can be made using quantile-quantile (Q-Q) plots (Fig. S10C) where the quantitation can be made in terms of offset and range values (Fig. S10C) obtained from the projection plots (for methods: refer (Lake, Cortes et al. 2012)).

I. Collagen fiber orientation distribution histograms were obtained for each ROI of fascicle using forward SHG images stack, with orientation obtained at each image taken 2 *μ*m apart and projected summed). Angles were taken b/w −90 to +90 deg. (0 deg. representing horizontal line)
II. Orientation of each ROI (local fascicle-axis) with respect to horizontal was determined by sum-projection of each ROI-stack. Straight line on ROI edges (upper and lower) were fitted to determine the average inclination of fascicle-axis.
III. Histograms representing distribution of collagen fibers with respect to local fascicle-axis within each ROI were obtained for angles b/w −90 to +90 degrees.
IV. The change in displacement between consecutive stripes were used to calculate deformation gradient along fascicle-axis (i.e. *F*_11_) within each ROI of a fascicle. Average cross-sectional area (CSA) along local tissue axis within each ROI was obtained in undeformed and deformed configurations of the sample by fitting ellipse to fascicle cross-section (various slices along longitudinal direction of fascicle and averaging). The change in radius was used to estimate deformation gradient in transverse direction (i.e. *F*_22_). *F*_21_ and *F*_12_ were assumed as negligible (set to zero) because sample loading was along fascicle-axis and only parallel photobleach stripes were considered for local strain estimations within a tissue). The obtained deformation gradient matrix was then used towards further calculations.
V. Orientation distributions of each ROI in undeformed and deformed configuration were grouped into 100 Quantiles (with 1% increment) using a custom MATLAB script. Angular values were approximated with linear interpolation wherever required.
VI. Quantile grouped data for undeformed configuration was used to obtain affine predicted orientation distributions for each ROI using respective deformation gradient matrix.
VII. Experimental and affine predicted orientation distributions were compared using Q-Q plots. Experimental values (100 groups) and affine predicted values (100 groups) were used to obtain projection plots where quantile difference (Exp. - Affine) were plotted along ordinate axis and the average of two ((Exp.+Affine)/2) was plotted as abscissa.
VIII. Offset and range were estimated to quantify deviation between experimental and affine predicted distributions in each ROI. Offset value represents the deviation (in degrees) of the median (of quantile differences) from 0 whereas range values represents +/-34.1% spread about median (i.e. 15.9 and 84.1 percentiles).
IX. Offset and Range values across all ROIs were compared

In order to obtain primary collagen fiber orientation with respect to fascicle-axis within different regions (i.e. ROIs) of an undeformed fascicle, obtained collagen fiber orientation distributions were grouped into 100-quantiles for angles with respect to fascicle-axis. Based on the extreme values of obtained quantile angles, collagen fiber distribution histograms were obtained for each class size of 2.5 deg. for angles between −45 to +45 degrees. Gauss curve was fitted to the obtained distribution (best fitting curve to each 60-degree interval) where Gauss-peak represented the primary collagen fiber orientation whereas Gauss-width indicated spread of orientations of fibers with respect to fascicle-axis).

### S7. Waviness estimation

Tissue waviness is calculated by the spatial averaging of local collagen fiber orientation changes when moving along longitudinal direction of a fascicle (Fig. S11) based on the following steps

I. Local collagen fiber orientation (w.r.t horizontal) vector-field per slice (pixel-size = ∼1×1 *μ*m) was obtained to represent local orientation within each slice of a tissue ROI
II. Pixels lying within the tissue at each slice were selected for further consideration while considering rest as background
III. Based on the orientation of each ROI (fascicle-axis orientation based on projected sum of forward SHG image) with respect to horizontal, slice-wise collagen fiber orientation with respect to fascicle-axis and corresponding pixel coordinates (x, y) were determined for each slice
IV. x-values and corresponding angles values were locally grouped and averaged into 30 bins 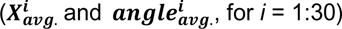 i.e. every ∼2 *μ*m along fascicle-axis
V. Based on 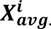 and 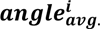 values and setting 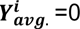 for *i*=1 was used (assuming starting 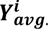 on fascicle center line). 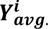 (for 1 < *i* <= 30) values were obtained using following relation to predict local waviness of slice based on local fiber orientation 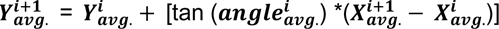
VI. Line segments between consecutive points 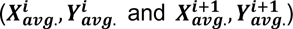 were then used to represent local length of curve and estimate total length of the curve (*L*_curve_)
VII. Displacement between starting and end points of the curve were then used to calculate end-to-end displacement (*L*_e2e_)
VIII. Waviness within each slice of a ROI was estimated based using the relation i.e. Waviness = ((*L*_curve_ /*L*_e2e_)-1)*100
IX. Above steps were then used to estimate the waviness values for all slices lying within an ROI of a fascicle covering entire tissue-depth.
X. Median value of waviness for each ROI was selected to represent the final waviness of a ROI.
XI. Normalized waviness value (with respect to waviness value before fascicle deformation) for a ROI with time (after addition of BC to tissue) were estimated to represent waviness variation with time

### S8. Fluorescent labelling of BC molecules at N-terminus

BC molecules were fluorescently labelled as per supplier protocol by mixing the two reactants in two different proportions per following steps

I. Fluorescent labelling of Bacterial Collagenase (BC) molecules was done with NHS-Ester 555 targeting N-terminus (for specifications of reactants, see materials)
II. Two different relative proportions of BC with respect to fluorophore were reacted as per supplier recommended protocol for 2 hours at 4 deg. i.e. 50 *μ*L (10 mg/mL) of fluorophore is added to 0.5 mL of 20 mg/mL (producing 4X-BC) or 1 ml of 10 mg/mL of BC (producing 1X-BC)
III. Overnight dialysis of the samples was done using 10 kDa cut-off (as mass of BC lies in 65-130 kDa range) in PBS at 4 deg.
IV. UV scanning of labelled BC samples and control samples (without labelling) were compared to verify BC labeling by fluorophore
V. Higher conc. (4X) of BC in reaction resulted in higher labelling efficiency (Fig. S19-A)
VI. Obtained conc. for 1X solution was 6.31 mg/mL whereas for 4X, it was 13.157 mg/mL (based on extinction coefficient of unlabeled-BC)

High Performance Liquid Chromatography (HPLC) was used to determine if all different masses of BC molecules present in the solution were labelled by the fluorophore (Fig. S19-B) as BC solution contains at least 7 different proteases ranging in molecular weight from 68-130 kDa.

I. Two different concentrations of fluo. BC solutions obtained through labelling were investigated where higher conc. BC to fluorophore showed more labelling
II. Samples of unlabeled-BC (1 mg/ml), Fluo-BC (13.1 mg/ml and 6.3 mg/ml) in PBS were kept at −20 deg. for long term storage and thawed at 4 deg. for 24 hours before testing.
III. Samples were diluted in PBS to similar final concentrations i.e. unlabeled-BC (1 mg/ml), Fluo BC (1.31mg/ml) and Fluo-BC (0.63 mg/ml) just before HPLC run
IV. Samples were run in polarity-based HPLC by transition of mobile phase from polar to non-polar ratio of 100:0 to 60:40 (DI water:Acetonitrile) in 30 mins
V. Signal intensity (peak height) at 280/555 was observed proportional to the sample concentration values
VI. Fluo-BC samples show small polar masses absorbing at 280 nm (highlighted by red box and ellipses) which could be because of self-degradation of the enzyme because of the presence of Alexa fluorophore at N-terminus of BC while they are not present in unlabeled-BC
VII. Both Fluo-BC samples show absorbance at 555 (highlighted by green box and ellipses) for range of masses while 555 absorbance is absent in unlabeled-BC indicating range of masses is fluorescently labelled

### S9. FRAP measurements

For each experiment, TRITC-dextran or fluo.-BC solutions in PBS were prepared and added to the medium carrying strained a fascicle sample making desired final concentrations and were kept overnight at 37 deg. before making FRAP measurements. All experiments were performed on Multi-photon microscope TCS SP8 MP (Leica) in FRAP mode using a 20 mW laser with a 20X /1.0 numerical aperture objective, with a 5.24 A.U. pinhole, which yielded a 7 *μ*m thick optical section. At center of image (of size 69.33 x 69.33 *μ*m at zoom = 7), a square ROI of 5 *μ*m x 5 *μ*m was selected as the region for bleaching. Each measurement was made in the regions situated deep into the tissue (at least than 25 *μ*m away from the fascicle surface or edges) and each image with 512 x 512 pixels took 1.295 seconds to acquire representing a time-point. For imaging, a lower value of laser intensity of 0.01% was selected to avoid photobleaching during image acquisition while the intensity was set to 90% for bleaching sessions. All of the FRAP experiments consisted of pre-bleach, bleach and post-bleach sessions for 303.3, 151.15 and 3367 seconds respectively (using bi-directional scan for extracting maximum data time points). Pre-bleach images were used for quantifying photobleaching during image acquisition at 0.01% laser intensity and making necessary corrections/normalization of the data.

For extracting characteristics of molecular kinetics from recover FRAP experiments, single exponential curves were fitted to normalized recovery data. In order to avoid fluctuations in the data for curve fitting purposes, the estimated ROI intensity was binned over constant intervals (of approximately 65 seconds). In order to normalize the data, double normalization was performed (Phair, Gorski et al. 2004) for the intensity values obtained from photobleached regions with respect to average background intensity as well as photobleach during image acquisition. After double normalization, photobleach experimental data was scaled between 0 to 1 with respect to maximum value as average intensity during pre-bleach session and minimum values as intensity value at end of beach session. For normalization protocol, we specified three different ROIs for bleached spot, background intensity and all image respectively, to obtain average fluorescent intensity in these ROIs at various time points of pre-bleach, bleach and post-bleach sessions. We used following normalization formulation to calculate normalized intensity in the bleached region for all data sets

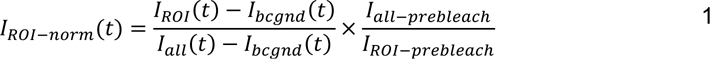

Where, *I*_*ROI*−*norm*_(*t*) is normalized intensity of bleached region, *I*_*ROI*_(*t*) average intensity from bleached region at all time points, *I*_*bcgnd*_(*t*) average background intensity at all time points, *I*_*all*_(t) average intensity of the full image at different time points, *I*_*all*−*prebleac*ℎ_ average intensity of all images averaged over all time points in pre-bleach session and *I*_*ROI*−*prebleach*_ as average intensity of bleached region averaged over all time points in pre-bleach session. Recovery characteristics of fluorescent molecules after photobleaching are obtained by fitting an exponential curve to recovery data where amplitude(s) and time (or decay) constant(s) of the curve represent mobile fraction(s) and time taken for recovery respectively as represented by following equation

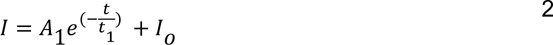

For single exponential curve fitting to recovery data, the mobile fraction value is given by

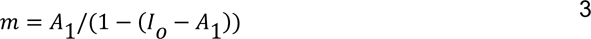

### S10. Synthesis and Purification of Fmoc-azGPO(tBu)-OH, and validation data

To streamline aza-peptide probe synthesis, protected tripeptide Fmoc-azGPO(tBu)-OH was synthesized and used as principal building block for SPPS. General protocol outlined below was developed using standard principles of Fmoc SPPS on 2-chlorotrityl chloride resin; one notable reference is included here (Bonkowski, Wieczorek et al. 2013).

#### Initial Resin Loading with Hydroxyproline

To a 500-mL SPPS vessel, Fmoc-Hyp(tBu)-OH (30.71 g, 75 mmol, 1.5 eq.) and DCM (200 mL) were added and stirred briefly. To this mixture, 2-chlorotrityl chloride resin (34.25 g, 50 mmol, 1.46 mmol/g) and DIEA (26.1 mL, 150 mmol, 3 eq.) were added. Reaction mixture was stirred for 5 h. After 5 h, MeOH (70 mL, ∼2 mL/g resin) was added to cap remaining reactive trityl groups, and resin was stirred for 1 h. Resin was washed with DMF (5x).

#### Proline coupling

N-terminal Fmoc group was removed by stirring resin in ∼200 mL 20% piperidine/DMF (2 x 30 min). Resin was washed with DMF (5x). Fmoc-Pro-OH (50.60 g, 150 mmol, 3 eq.), HBTU (56.89 g, 150 mmol, 3 eq.), DIEA (52.3 mL, 300 mmol, 6 eq.), and DMF (300 mL) were combined in a 1000-mL round bottom flask and stirred to activate for several minutes. Activated coupling solution was added to resin and stirred overnight (15.5 h). Resin was washed with DMF (5x).

#### Aza-glycine coupling

Deprotection and wash steps were carried out as described above. CDT (24.62 g, 150 mmol, 3 eq.), Fmoc-carbazate (38.14 g, 150 mmol, 3 eq.), and DMF (350 mL) were combined in a 1000-mL round bottom flask and stirred to activate for several minutes. Activated coupling solution was added to resin and stirred for 2 days (45.5 h). Resin was washed with DCM (3x) followed by MeOH (3x) and dried by vacuum.

#### Cleavage & Drying

Dry resin was transferred to a 2000-mL round bottom flask. Protected tripeptide was cleaved from resin by stirring in a 1-L cleavage cocktail of 30.0% TFE, 0.3% TFA in DCM for 4 h. Reaction mixture was filtered through filter paper into separate 2000-mL round bottom flask. Resin was rinsed several times with DCM and rinses were filtered into crude peptide solution. Solvent was evaporated *in vacuo*.

#### Purification

Crude product was purified by flash chromatography (ISCO) in gradient of 70-100% EtOAc/Hex (EtOAc containing 0.1% TFA). Fractions containing product were pooled and dried *in vacuo*. Purified product was azeotroped 3 times with ∼300 mL toluene. Final product was dried on hi-vac and aliquoted as needed.

#### Validation Data

HPLC gradients and conditions are as noted. CHCA was used as matrix for all MALDI-TOF MS. For each probe, HPLC and MALDI-TOF MS data for corresponding 5-CTAMRA-labeled isomers (removed during purification) are shown alongside data for purified 6-CTAMRA products used in assays. For validation data, see Fig. S25 and Fig. S26.

---

**Fig. S1.**
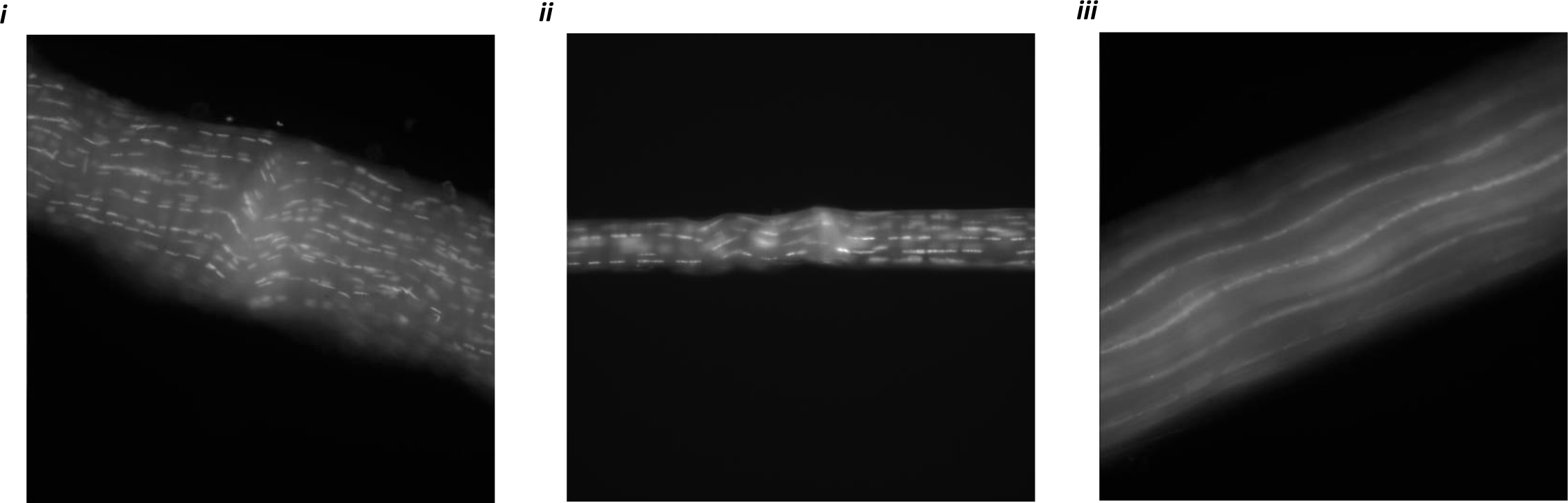
Freeze-thaw cycles produces intact ECM of fascicles as a result of cell-death. Chromatin distribution (using Hoechst staining) in **(*i*-*ii*)** Live fascicle samples from WT mice tail (immediately after euthanizing) and from NSG mice tail (after FA fixing) together indicate localized chromatin. **(*iii*)** Starved fascicle from WT mice after freeze thaw cycles and long-term storage (few months) at −20 °C produces distributed chromatin along fascicle length.

**Fig. S2.**
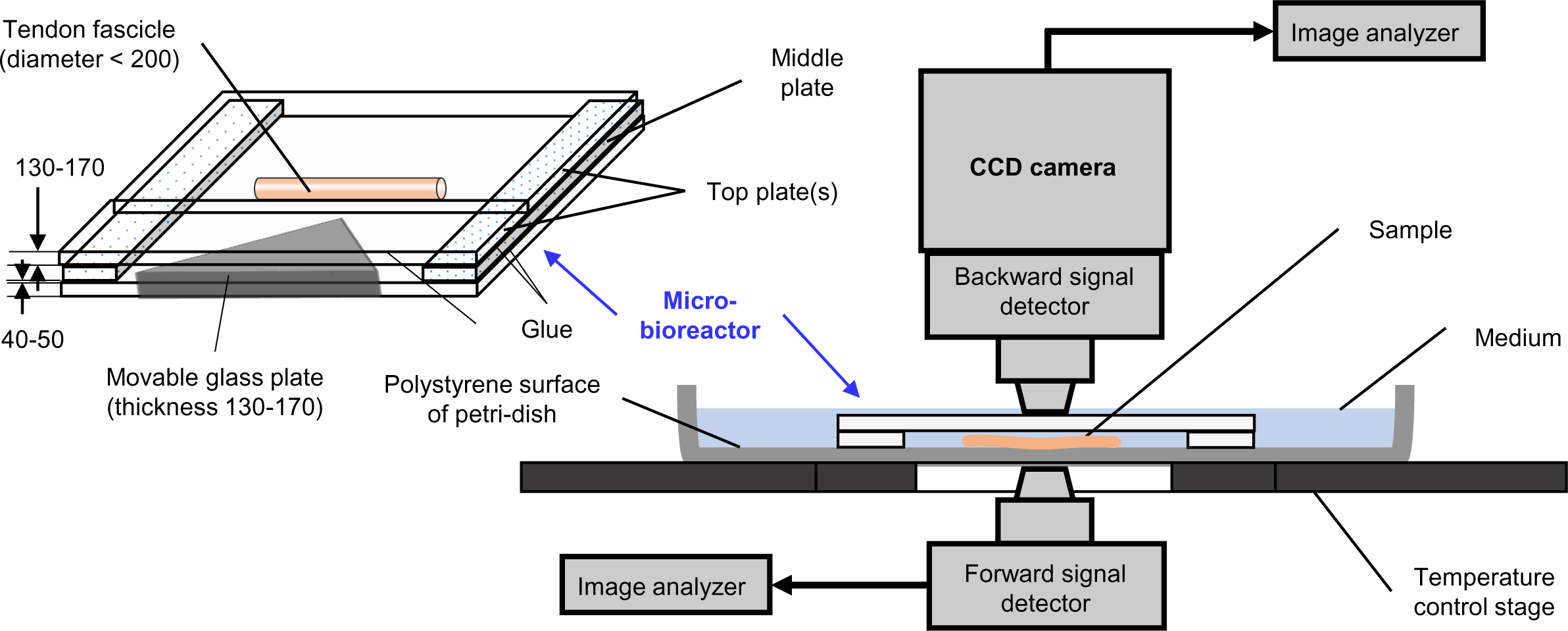
Schematic of a micro-bioreactor for heterogeneous deformation of a fascicle and its installation on a MP-microscope for SHG imaging of mechanically strained fascicles (all dimensions in microns).

**Fig. S3:**
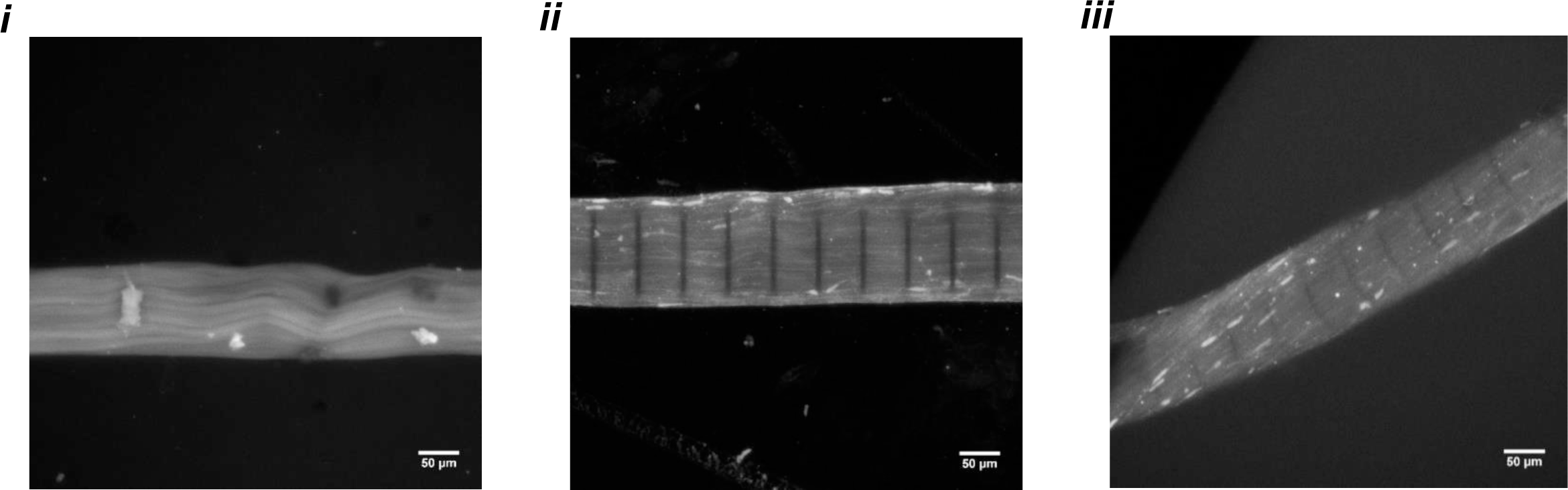
Aza-peptide show affinity towards mechanically unstrained and strained tendon fascicles. **(*i*)** Mechanically unstrained fascicle labelled with fluorescent aza-peptide. **(*ii*)** Mechanically unstrained fascicle with fluorescent aza-peptide after pattern photobleaching. **(*iii*)** Mechanically strained fascicle labelled with fluorescent aza-peptide. Scale bar = 50 *μ*m.

**Fig. S4.**
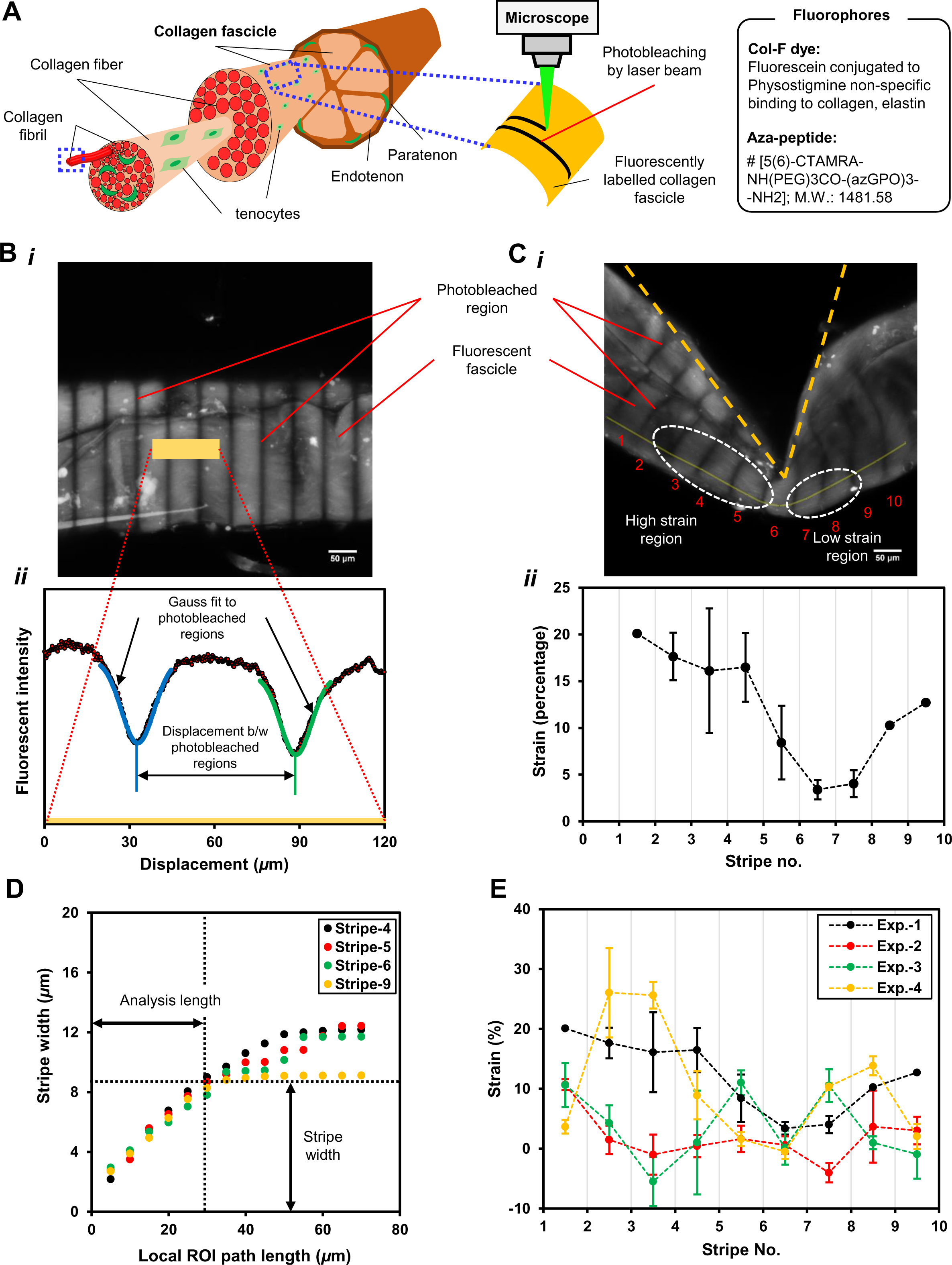
Structural hierarchy level of a fascicle (i.e. bundle of collagen fibers) in mice tail tendon tissue and local strain quantification using pattern photobleaching. **A.** Fluorescent labelling of a fascicle (by Col-F dye or aza-peptide) for creating a pattern through photobleaching. **B. (*i*)** Pattern photobleaching of fluorescently labelled unstrained fascicle with a representative path (yellow bar) across stripes (or photobleached regions) selected for intensity measurement on the composite image (projected sum). **(*ii*)** Intensity profile along the representative path, with Gaussian curves fitted for local displacement quantification. **C. (*i*)** Mechanically deformed fluorescent fascicle representing regions of high and low strain magnitudes. **(*ii*)** Mechanical strain magnitudes between each pair of consecutive stripes i.e. ROIs along the longitudinal direction (shown as light yellow curve) of fascicle. **D.** Gaussian width increases as a function of ROI path length indicating local tissue heterogeneity. ROI path length of 25 *μ*m has been selected as the length for determining Gauss peak position on the fascicle throughout the study. **E.** Strain distribution for different ROIs of various samples indicate non-repeatable strain distributions under heterogeneous deformation and indicating heterogeneity of strain distribution.

**Fig. S5.**
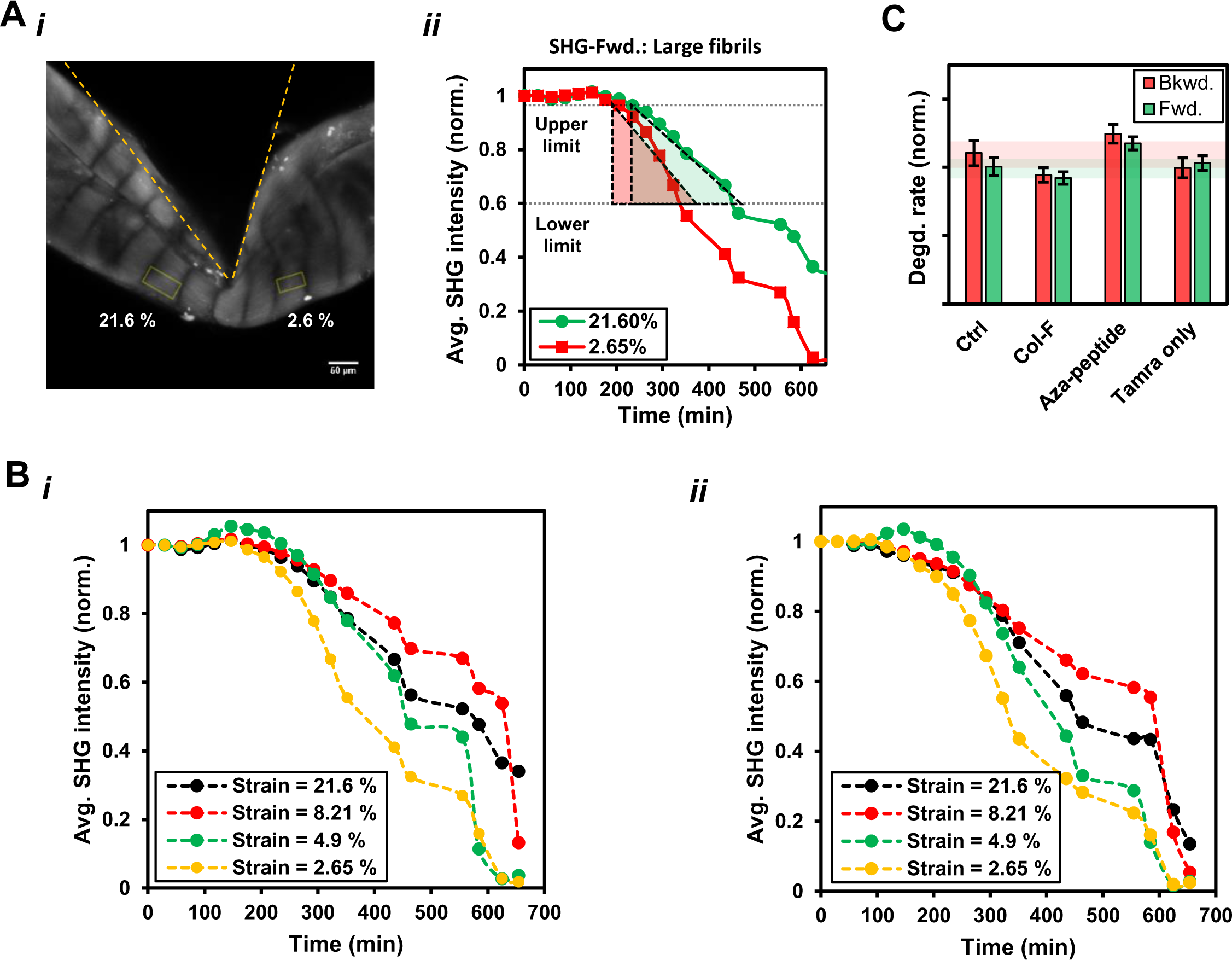
Tissue collagen degradation rate measurement within different strain carrying regions (or ROI) of a fascicle. Fluorescent labelling of the fascicle was observed not to interfere with the tissue collagen degradation. **A. (*i*-*ii*)** Representative ROIs located in the high and low strain region of a fascicle (left panel) over which SHG signal (based on the composite image obtained by projected sum of z-stack) was locally averaged. Average forward SHG signal from these regions decreases at different rates when the rate of each strained region is represented as the slope of the average SHG intensity versus time data. For slope measurements, the initial phase of the experiment was considered (normalized SHG intensity change from 0.95 to 0.60 (∼30-40% approx.) to minimize the influence of strain relaxation or change in collagenase/substrate concentrations. **B.** Average SHG signal intensity with time from the different strain regions of a fascicle that is mechanically deformed and exposed to BC using **(*i*).** Average backward SHG signal **(*ii*)**. Average forward SHG signal. **C.** The degradation rates of the fascicles labelled by different fluorophores *i.e.* Col-F dye, Aza-peptide, TAMRA-only (fluorophore with peg-linker) or un-labelled (or control) by BC were not significantly different. Col-F labelled fascicles degrade at similar rates as unlabeled samples, whereas degradation rate is the same or slightly higher for samples labelled with Aza-peptide. TAMRA only didn’t show similar binding affinity to samples as shown by aza-peptide.

**Fig. S6.**
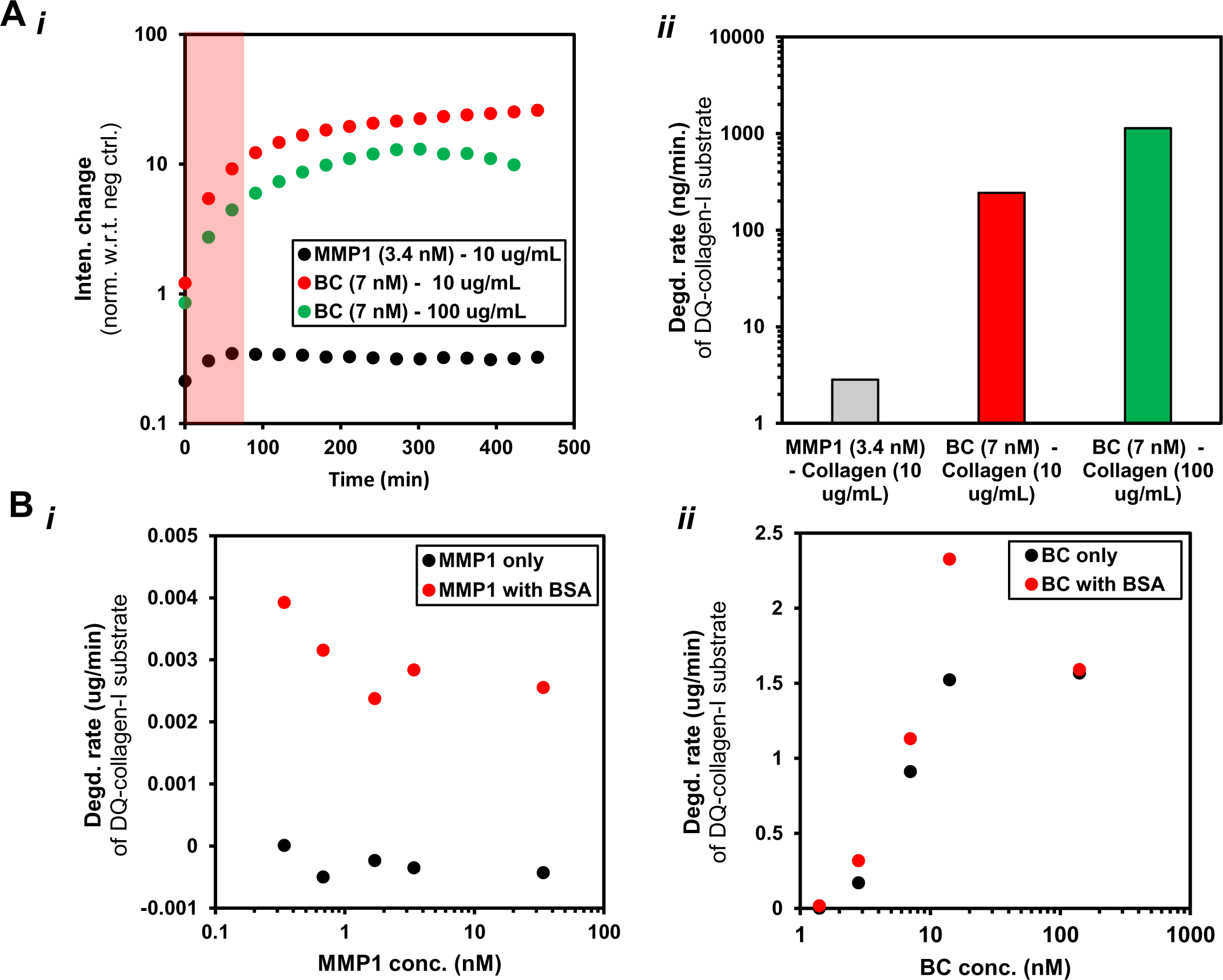
The comparison of activity of MMP-1 and BC on monomeric collagen type-I fluorescein conjugate (DQ™ Collagen type-I from bovine skin). **A. (*i*-*ii*)** BC cleaves monomeric collagen type-I (from bovine skin), fluorescein conjugate almost ∼50 fold faster than MMP-1 as indicated by enzyme-linked immunosorbent assay (ELISA) **B. (*i*-*ii*).** MMP-1 activity is influenced by the presence of BSA (bovine serum albumin 0.01 % solution) in the medium while BC activity remains unaffected by the presence of BSA.

**Fig. S7.**
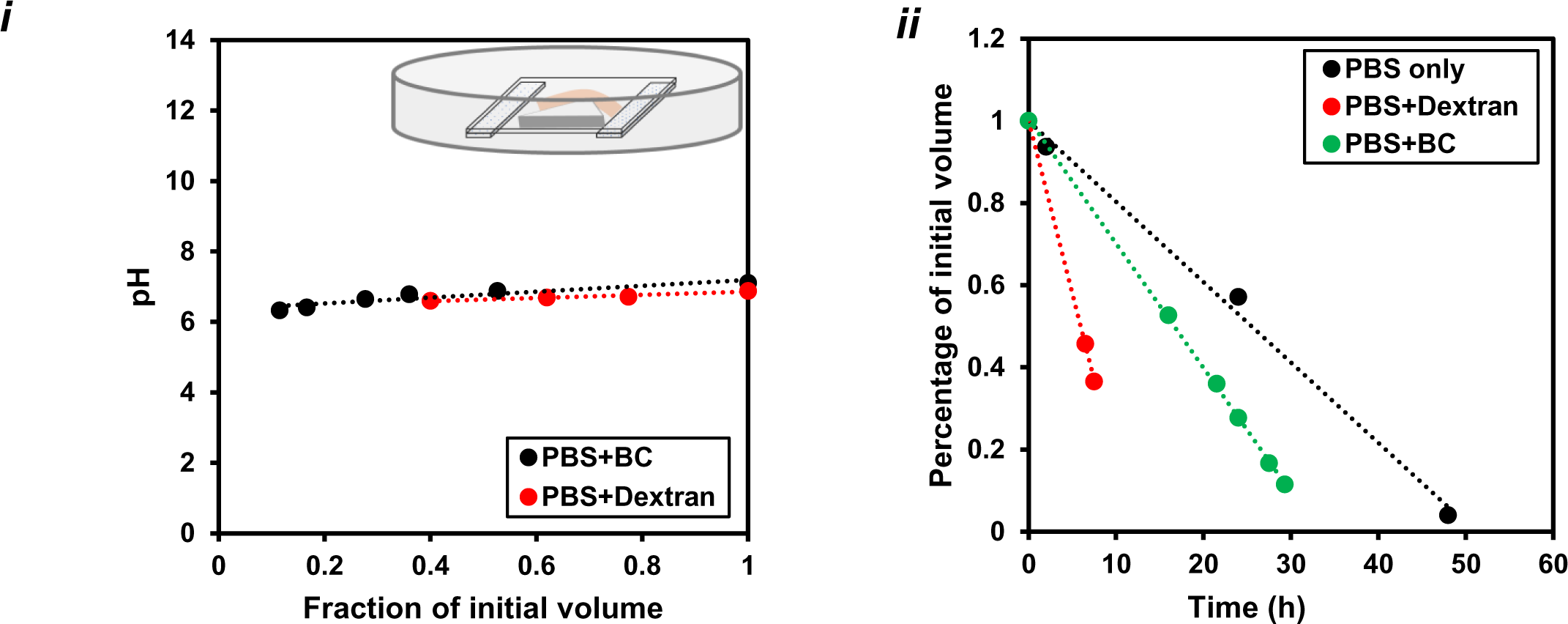
The evaporation of reaction buffer from micro-bioreactor does not affect its pH considerably. The evaporation rate of reaction buffer is influenced by the presence of different solute(s). **(*i*-*ii*).** Reaction buffer evaporates from edges of micro-bioreactor (polystyrene petridish of 5 cm diameter and maintained at 37 deg. C; initial volume = 5 mL). Evaporation rate (based on normalized volume of reaction buffer with time) of the reaction buffer depends on the solute (BC or TRITC-dextran) i.e. evaporation rate is higher for BC and highest for TRITC-dextran when PBS act as solvent.

**Fig. S8.**
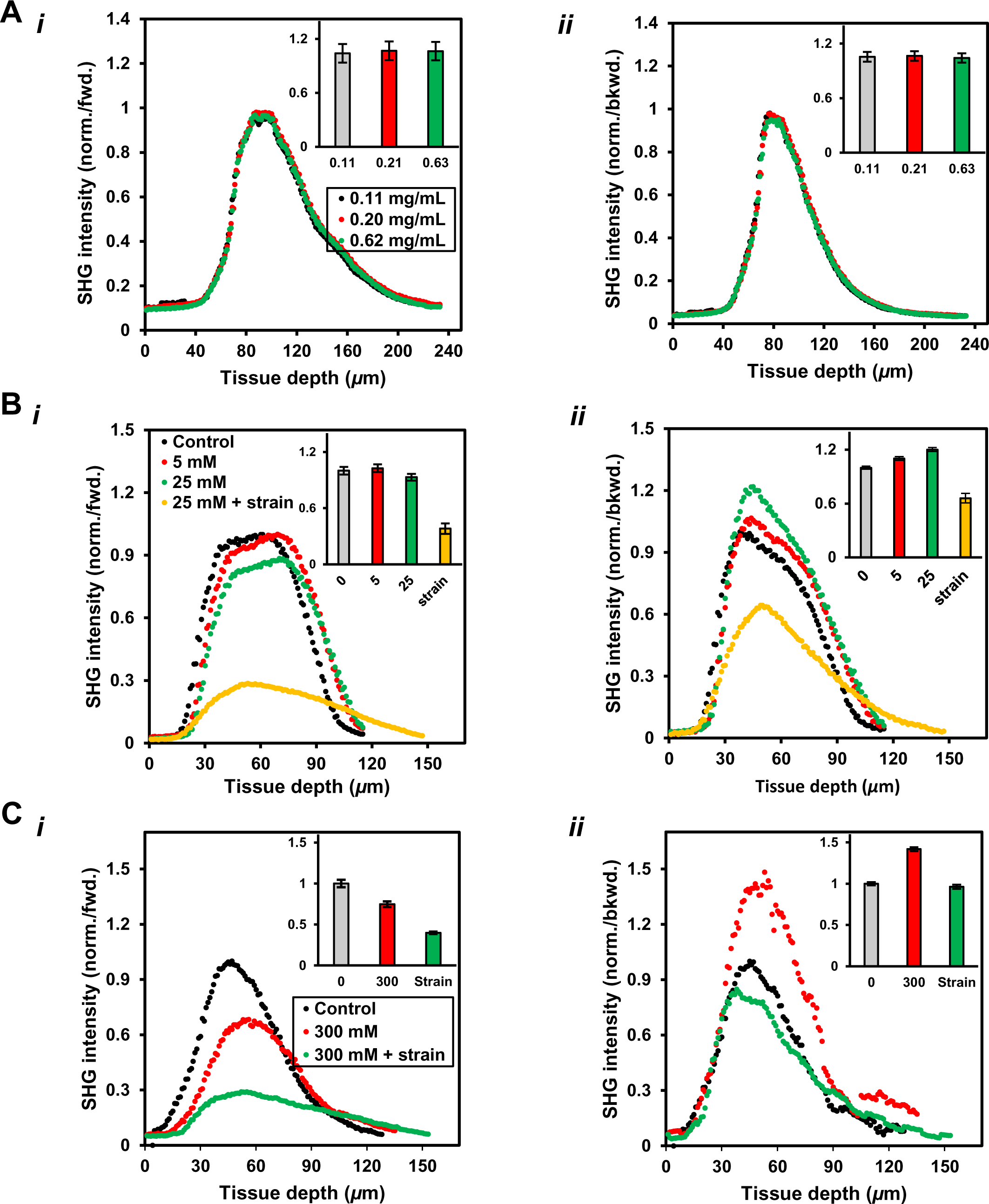
The change in the composition of reaction buffer might affect molecular packing of collagen fibrils present in the fascicles as indicated by SHG signal in forward and backward directions. **A. (*i*-*ii*)** Dextran concentration change in the medium as a result of evaporation do not affect the packing of larger and smaller size collagen fibrils as reflected by intensity of SHG signal with depth in forward and backward direction. **B. (*i*-*ii*)** The increase in CaCl_2_ salt concentration promotes disassembly of larger diameter collagen fibrils in fascicles into smaller size fibrils while superimposing mechanical strain on a fascicle (in presence of CaCl_2_) promotes further disassembly of all sizes collagen fibrils. CaCl_2_ concentration range:-physiological: 1.1–1.4 mM; fluctuating (in ion channels): 0.5–80 mM). **C. (*i*-*ii*)** The increase in NaCl salt concentration promotes disassembly of larger diameter fibrils resulting more and more smaller diameter fibrils appearing in backward SHG signal whereas strain in presence of NaCl leading disassembly of

**Fig. S9.**
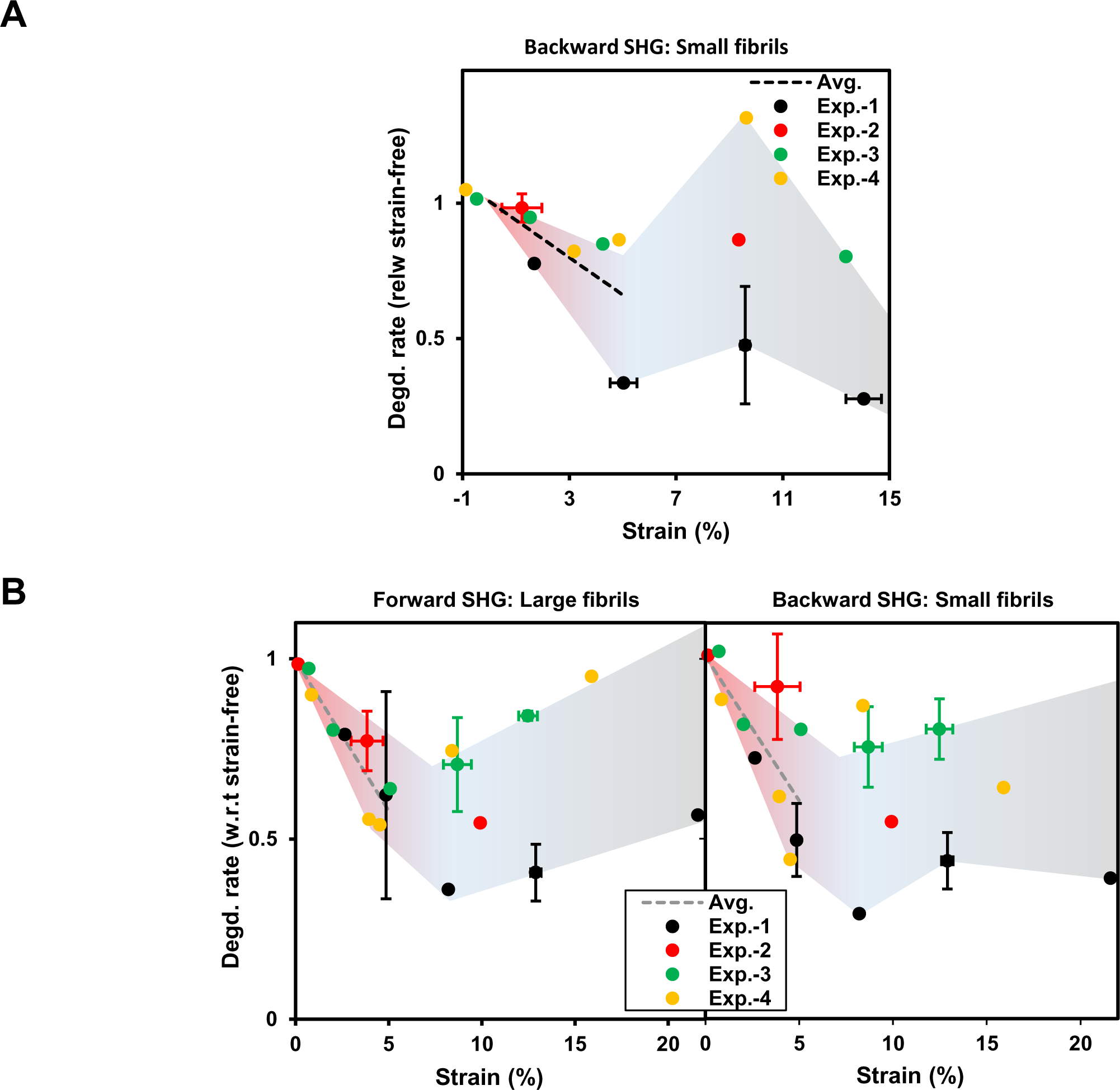
Heterogeneous mechanical strain decreases the degradation rate of fibrillar collagen in the fascicles within physiological strain limits when exposed to different collagenases. Strain magnitude dependence of degradation rate when heterogeneously strained fascicles are exposed to **A.** MMP-1 based on backward SHG signal **B.** Bacterial collagenase (BC) based on forward and backward SHG signal.

**Fig. S10.**
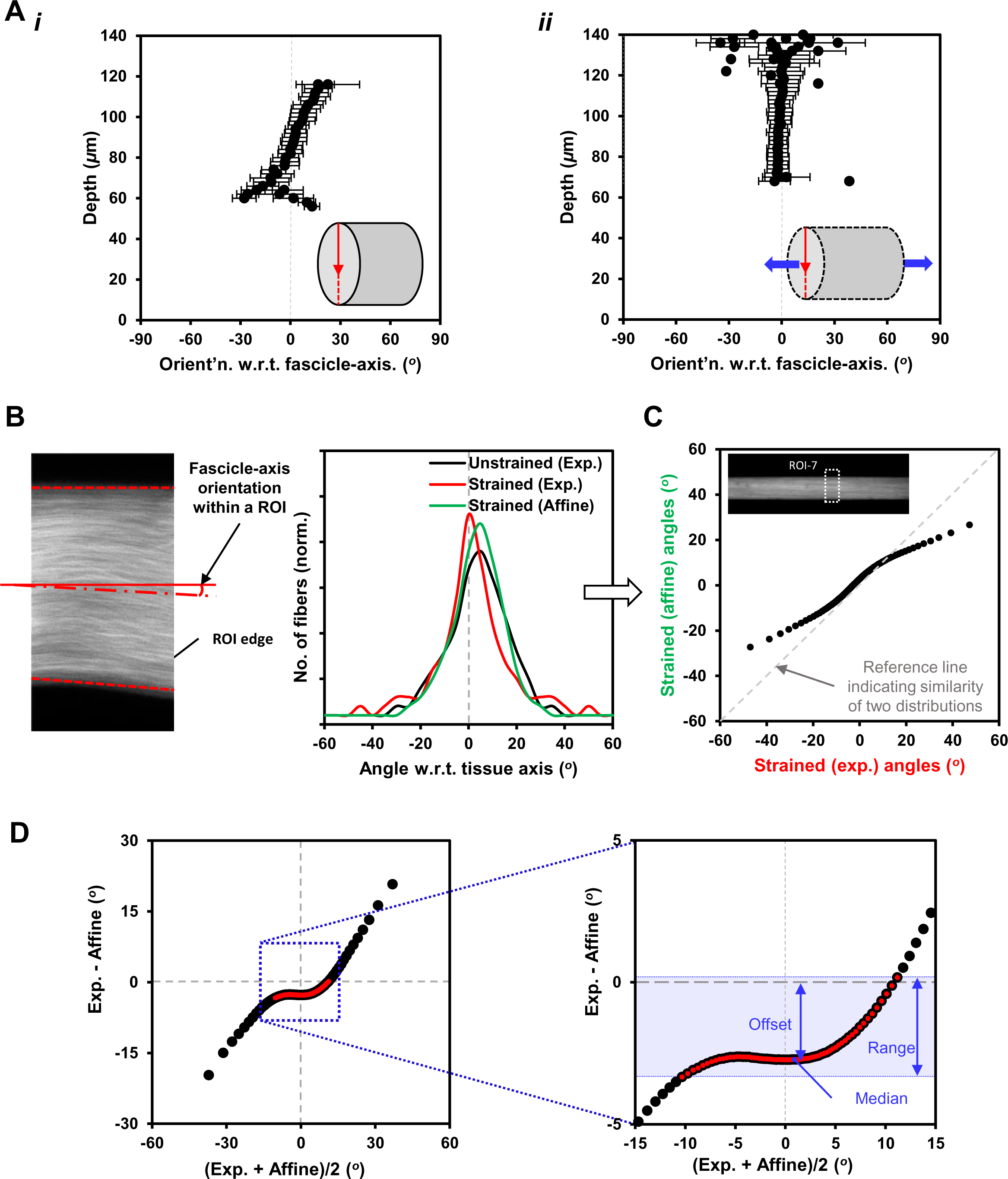
Each ROI of a fascicle carries both aligned and inclined fibers (with respect to fascicle-axis) in the undeformed configuration and the fibers tend to orient along fascicle-axis upon the deformation of fascicle. The deformation behavior is quantitatively analyzed by means of projection plots. **A. (*i*-*ii*)**. Slice-wise primary fiber orientation with respect to tissue depth (along sample diameter as indicated by red-arrow) in undeformed and deformed configurations of a ROI. **B.** Local fascicle-axis orientation within a ROI is estimated based on fascicle geometry predicted by forward SHG signal. **C.** The distribution of collagen fiber-orientations with respect to fascicle-axis within a representative ROI of a fascicle in undeformed and deformed configurations along with affine predicted orientation distribution. Comparison of experimentally and affine deformed fiber orientation distributions of same ROI by means of quantile-quantile (Q-Q) plot. fascicle (based on quantile grouping). **D.** Projection plot for quantitative comparison of experimental and affine predicted distributions for a ROI. Deviation between two populations and selection of ±*σ* (i.e. ±34.1% indicated by red-dots) spread of quantile differences about median for quantitative prediction of offset and range values (methods: Lake et al, Biomech Model Mechanobiol. 2012).

**Fig. S11.**
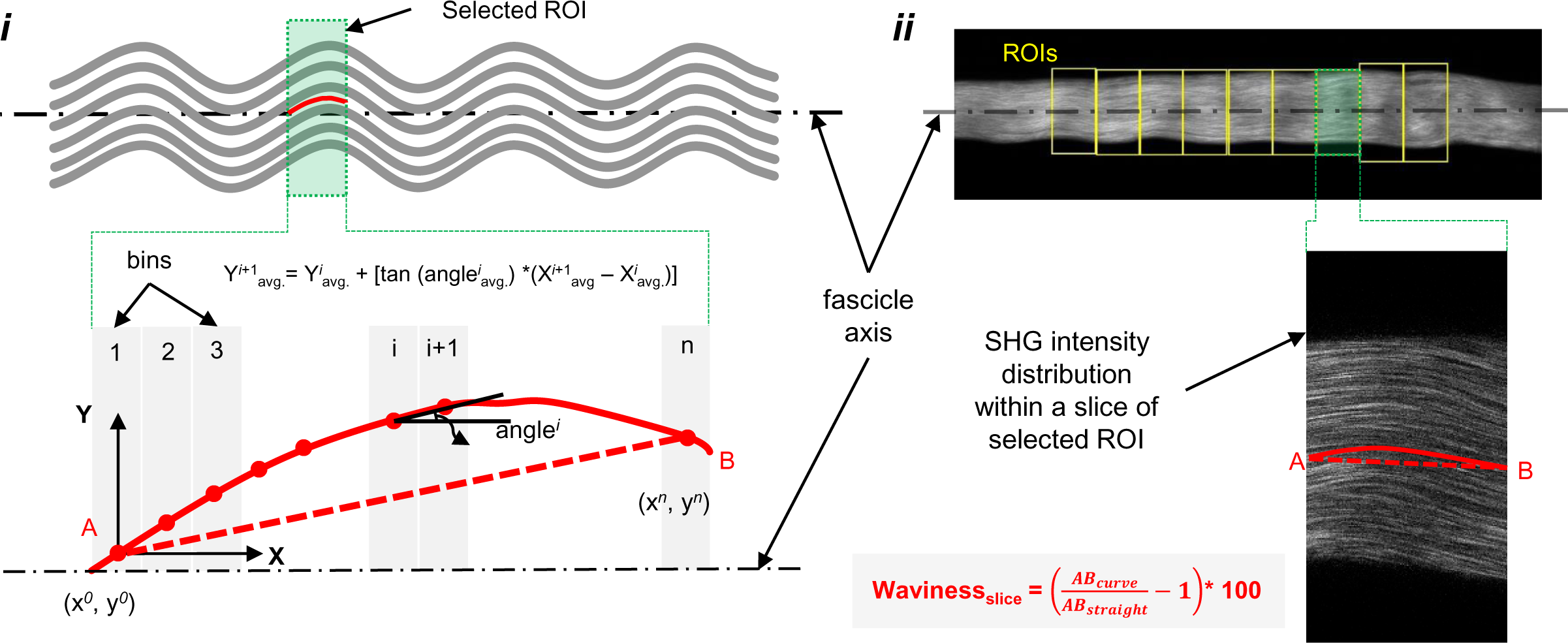
The waviness measurement for each slice is based on spatial averaging of fiber orientations where orientation distributions are measured based on forward SHG signal while the local waviness magnitude for a ROI represents the median value of waviness values measured for all slices within a ROI. **(*i*)** Waviness quantification from spatial distribution of fiber orientations. **(*ii*)** Forward SHG image (projected sum) of a tendon tissue sample indicating various ROIs (top), and single slice within a ROI selected for waviness analysis (bottom).

**Fig. S12.**
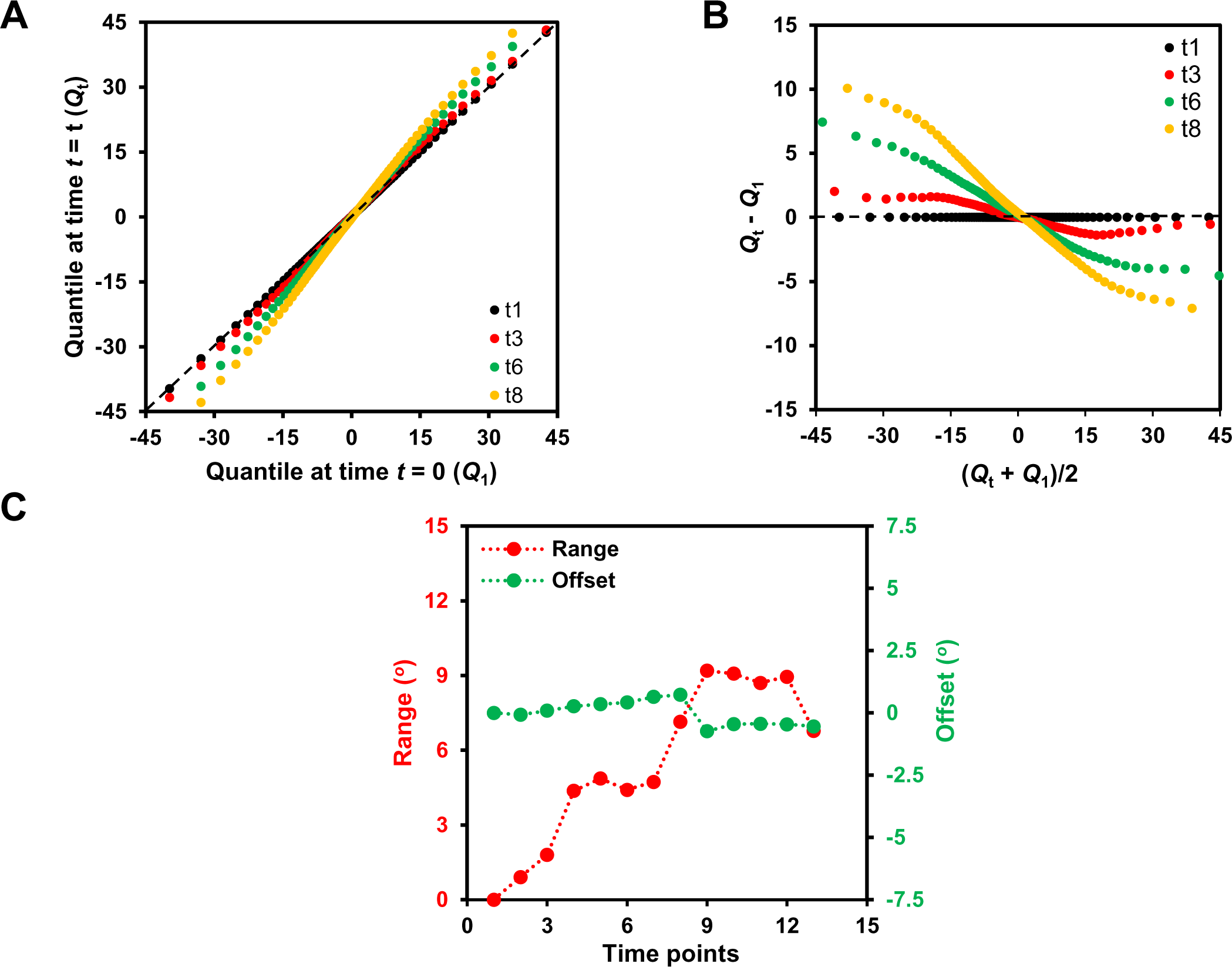
Projection plots for comparison of orientation distributions within a ROI at several time-points of fascicle degradation experiment and estimation of range and offset parameters. **A.** Quantile-quantile (*Q-Q*) plot for a ROI at four different time points indicating “fat-tails”. **B.** Corresponding projection plots at time points on the basis of mean and difference of quantiles. **C.** Range and offset magnitudes at various time points in a ROI where offset values remains almost unaltered while range values change considerably. Deviation between two populations and selection of ±*σ* (i.e. ±34.1% indicated by red-dots) spread of quantile differences about median for quantitative prediction of offset and range values where median value represents offset while the spread of quantile differences values about median indicates range magnitude also shown in Fig. S10D (for methods: Lake et al, Biomech Model Mechanobiol. 2012).

**Fig. S13.**
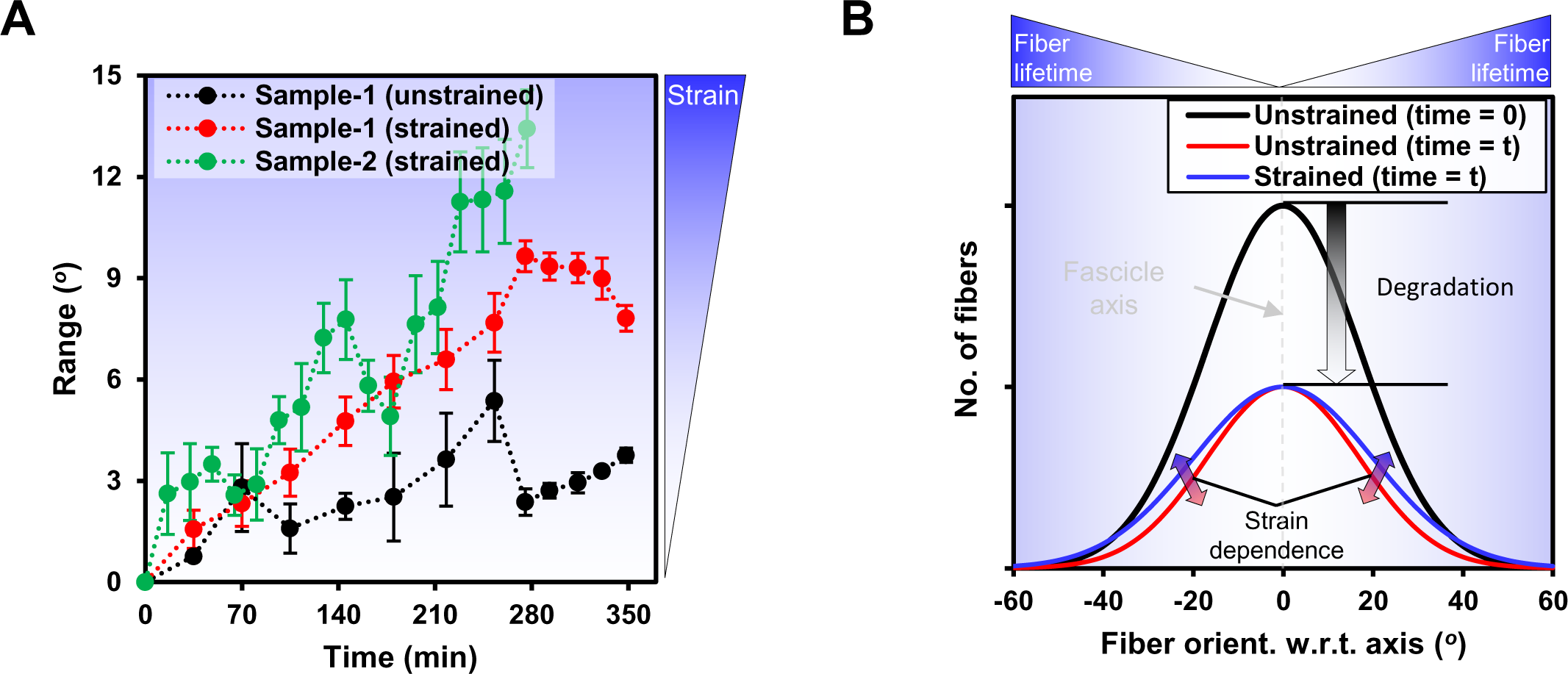
Degradation of uni-axially deformed tendon fascicle by BC results more inclined fibers (with respect to fascicle-axis) with time than that of strain-free samples. For different ROIs of samples exposed to BC (also shown in heat-maps Fig. 2D), **A.** Quantitative comparison of fiber orientation distributions of various ROIs within strained and unstrained fascicles by means of range values obtained from projection plots. A larger increase in the range magnitudes with time in mechanically strained samples as compared to that of unstrained samples. Dots in Fig. S12(A-B) represent the parameters (i.e. waviness and orientation) values and corresponding strain values for mechanically (un)deformed samples (indicated by dot color) at various time points during exposure to BC whereas solid lines indicate respective bin values of strains (and corresponding parameter values) for each sample. **B.** Fibers in an unstrained fascicle (black-line) follow nearly uniform degradation (red-line) as indicated by smaller change in range values with time while mechanically strained samples ends up with more inclined fibers (blue-line) relative to aligned ones as indicated by comparatively larger range values with time.

**Fig. S14.**
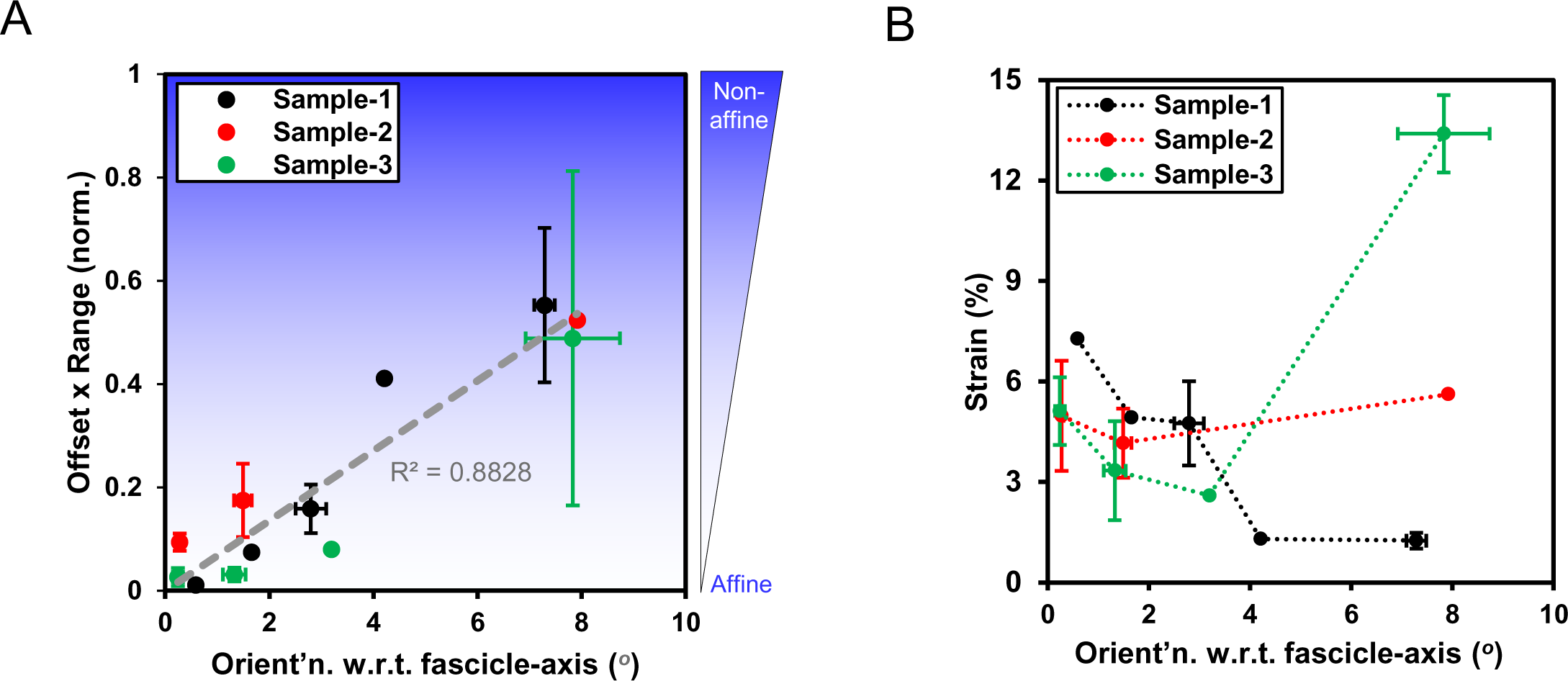
Local tissue non-affine behavior and stiffness variation within tendon fascicles subjected to uni-axial deformation depends on local fiber orientation distribution with respect to fascicle-axis. **A.** Regions of tendon fascicle carrying aligned fibers (with respect to fascicle-axis) follow affine deformation whereas the regions dominated by inclined fibers tend to follow non-affine deformation. Non-affine behavior indicated by the product of offset and range values (each parameter normalized with respect to maximum value within each experiment) estimated based on projection plots (see supp. info. Fig S10D). **B.** Local strain magnitudes under uniaxial deformation of fascicle (indicating local tissue stiffness) depends on the local fiber orientation (with respect to fascicle-axis) in the undeformed sample.

**Fig. S15.**
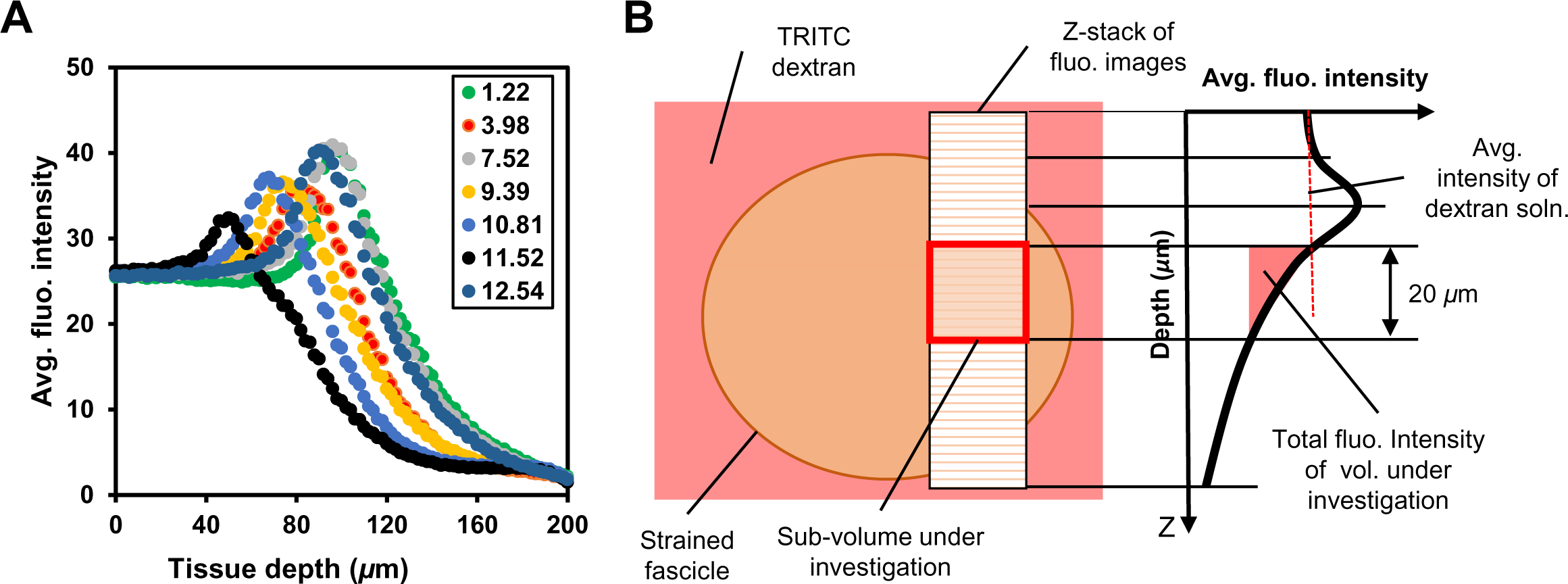
Quantification of TRITC-dextran (1.4 nm Stoke’s radius) permeation into various mechanically strained sub-volumes located at constant distance from surface of a fascicle using volume averaged fluorescent intensity within the sub-volume. **A.** Average fluorescent intensity varies with tissue depth (z-direction) for various ROIs (carrying different strain values (%) shown as legends) of a fascicle. **B.** The schematic of a fascicle’s cross-section and representative curve showing area average fluorescent intensity variation with sample depth (z-direction where peak might have been outcome of curved surface of the fascicle). Sub-volume under investigation for volume average fluorescent intensity measurements lies at approx. constant distance from the fascicle surface for each ROI. Z-depth of the sub-volume is 20 *µ*m (as higher depths are prone to fluorescent signal attenuation as a result of sample thickness) with its top edge lying at z-position where average fluorescent intensity becomes almost equal to background intensity (i.e. point of intersection of average intensity curve and line representing average fluorescent intensity of dextran solution).

**Fig. S16.**
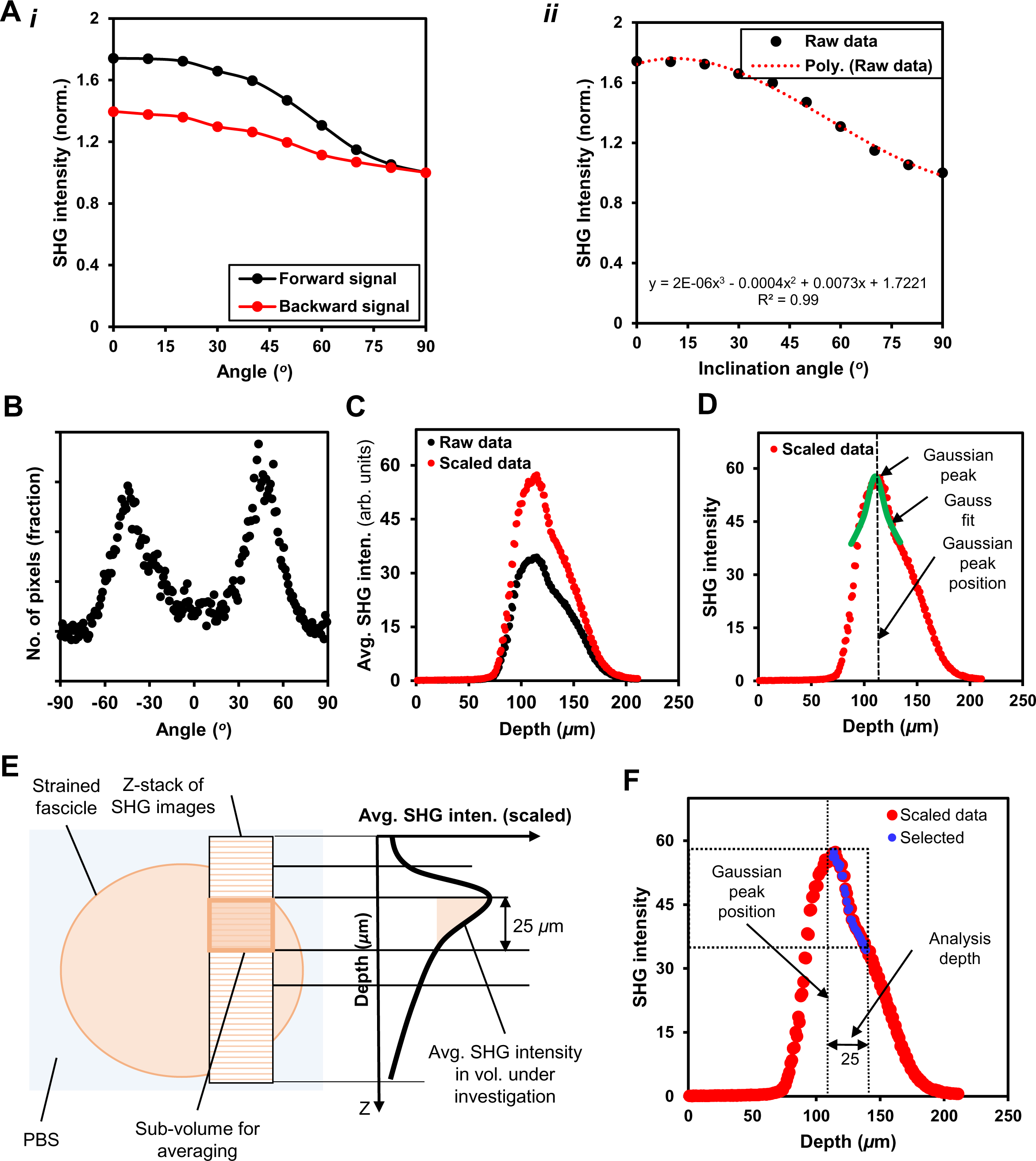
Collagen concentration measurement (absolute SHG intensity per unit volume) in each ROI of a mechanically strained fascicle sample. **A. (*i*)** SHG intensity variation as function of inclination angle between fascicle longitudinal axis and the direction of incident light polarization. **(*ii*)** The variation of average SHG intensity from a fascicle sample as a function of the inclination angle and function used for interpolation of values. **B.** Pixel-wise collagen fiber orientation distribution determination within a slice of the z-stack based on structure tensor [for methods, see Rezakhaniha, R., et al., 2012, Biomech. & Model. in Mechnblgy., 2012]. **C.** The absolute (or scaled) SHG intensity of each image (of a ROI shown as red dot) is determined by the scaling of measured average intensity (black dots as raw data) based on constituting fiber orientation distribution and angular inclination (orientation of each pixel with respect to polarization direction of incident light) using interpolation function and used fot estimating tissue depth dependent absolute SHG intensity. **D.** The location of peak (Gaussian fit) on tissue depth dependent abs. SHG data has been used predict the surface of the fascicle within each z-stack of a ROI. **E & F.** Sub-volume edges are determined by using the Gaussian peak as reference while keeping sub-volume depth as 25 *µ*m (for all measurements) and selecting points for volumetric averaging (blue-points).

**Fig. S17.**
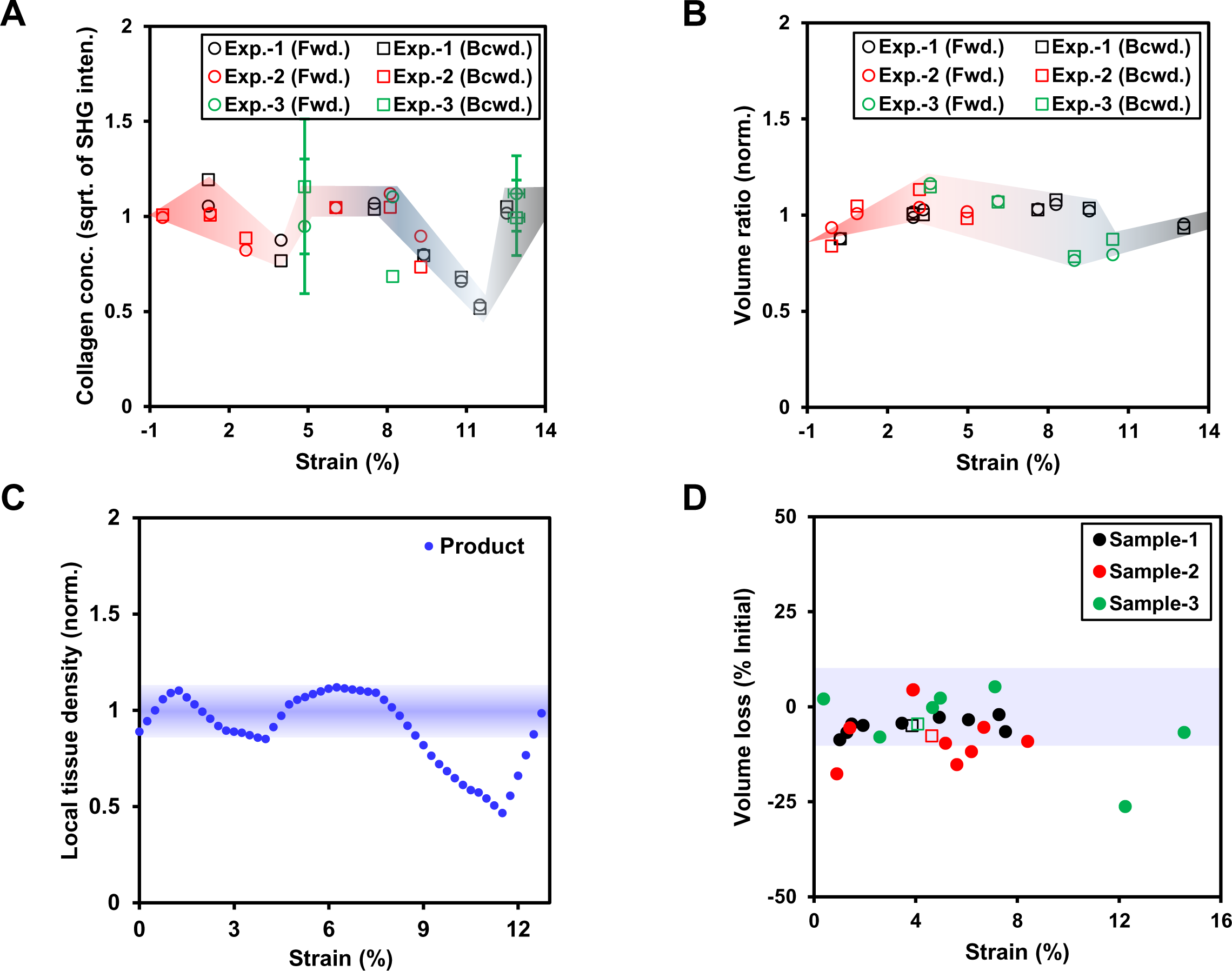
Tissue mass density based on SHG signal fluctuates for a fascicle that is deformed within physiological strain limits (up to ∼8%) but remains within ±10% of its magnitude for heterogeneous and uni-axial deformation modes. **A.** Average collagen concentration for various deformed ROIs (carrying different strains) is measured as square-root of average SHG signal from the selected sub-volumes (per Fig. S16). The estimated collagen concentrations at various strain values are normalized with respect to average SHG signal value for strains values up to ∼8% for each heterogeneously deformed fascicle and averaged for forward and backward directions at each strain value. **B.** The local tissue volume (i.e. ellipsoid between consecutive parallel stripes) at different strain magnitudes within the heterogeneously deformed fascicle is represented as the normalized volume ratio of each region calculated by the ratio of absolute volume of deformed region to its initial volume. The geometric limits of each local volume are obtained by fitting an ellipse to the cross-section of fascicle based on SHG signal. **C.** Local tissue mass density for each heterogeneously deformed fascicle is obtained as the product of collagen concentration (based on deformed volume) and normalized volume ratio values at each strain magnitude (on the basis of piece-wise line fits), showing insignificant variation (within ±10%) in tissue collagen density within strain magnitudes up to ∼8%. **D.** Percentage volume loss based on Poisson’s ratio (PR) at increasing strain magnitudes within uni-axially deformed fascicle stays within ±10% of its average magnitude. Filled circles indicate local PR of each region for sample while empty squares indicate global PR values based on overall size of sample (between two extreme stripes).

**Fig. S18.**
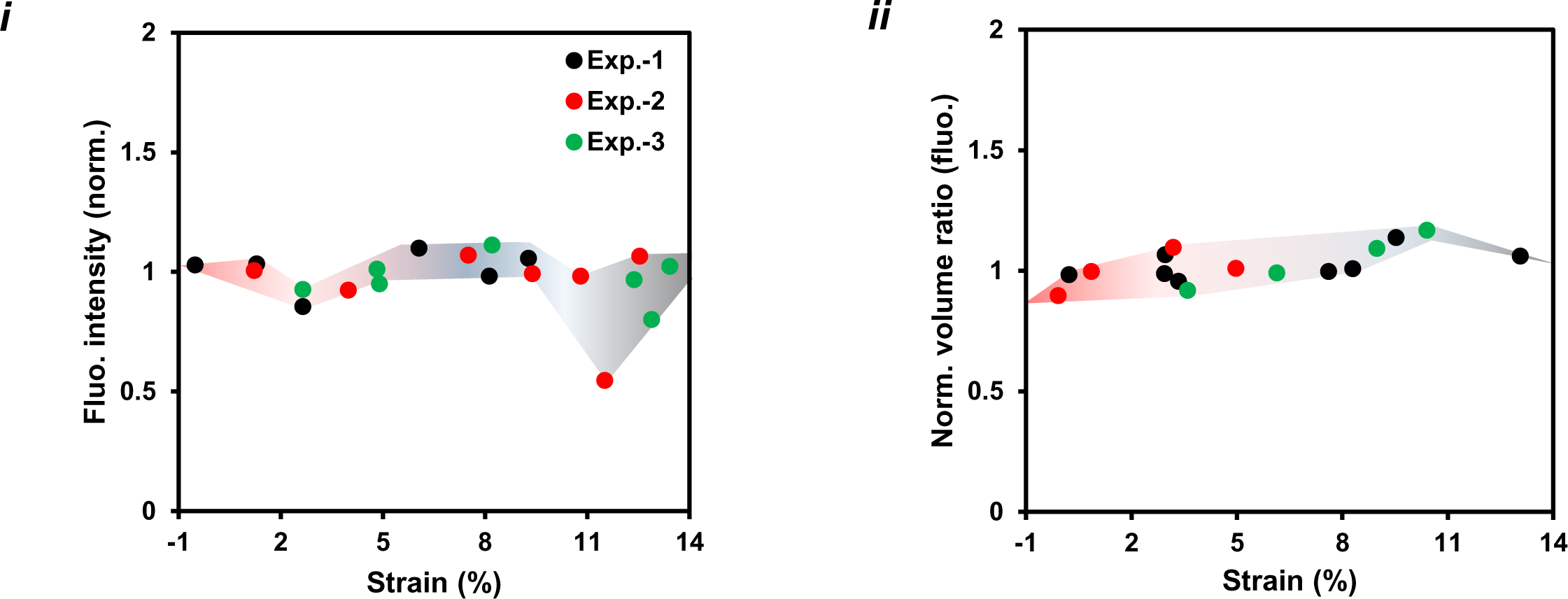
Volumetric and collagen content measurements at various strain magnitudes within each fascicle based on Col-F dye are different from SHG based collagen content measurements. **(*i*).** Collagen content versus strain measurements. **(*ii*).** Volume ratio versus strain.

**Fig. S19.**
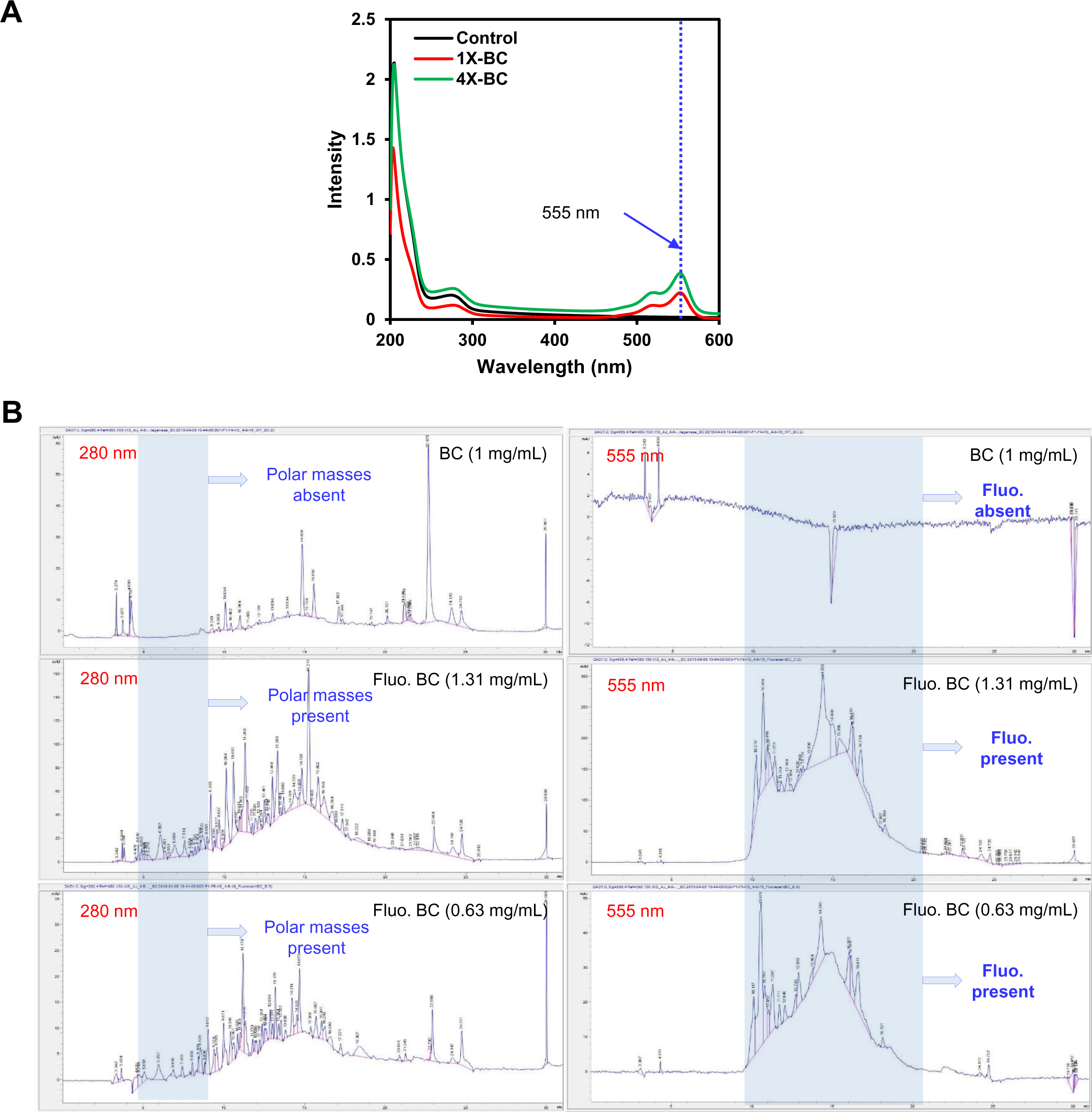
Fluorescent labelling of BC molecules at their N-terminus by Alexa Fluor™ 555 NHS Ester (Succinimidyl Ester). **A.** Higher proportion of BC with respect to fluorophore (Alexa Fluor™ 555 NHS Ester) results in higher labelling efficiency as indicated by UV scanning of fluorescently labelled samples at 555 nm. 4X: 50 *μ*L of fluorophore (10mg/mL) is reacted to 0.5 mL of BC at 20 mg/mL; 1X: 50 *μ*L of fluorophore (10 mg/mL) reacted with 1 mL of 10 mg/mL of BC. **B.** Unlabelled and fluo. labelled BC samples (at ∼ 1 mg/mL) on HPLC show the presence of fluorescently labelled masses (at 555 nm). Peaks at 280 nm (highlighted in sky-blue) indicate presence of polar masses in fluo. BC that might be resulting from self degradation of fluo. BC.

**Fig. S20.**
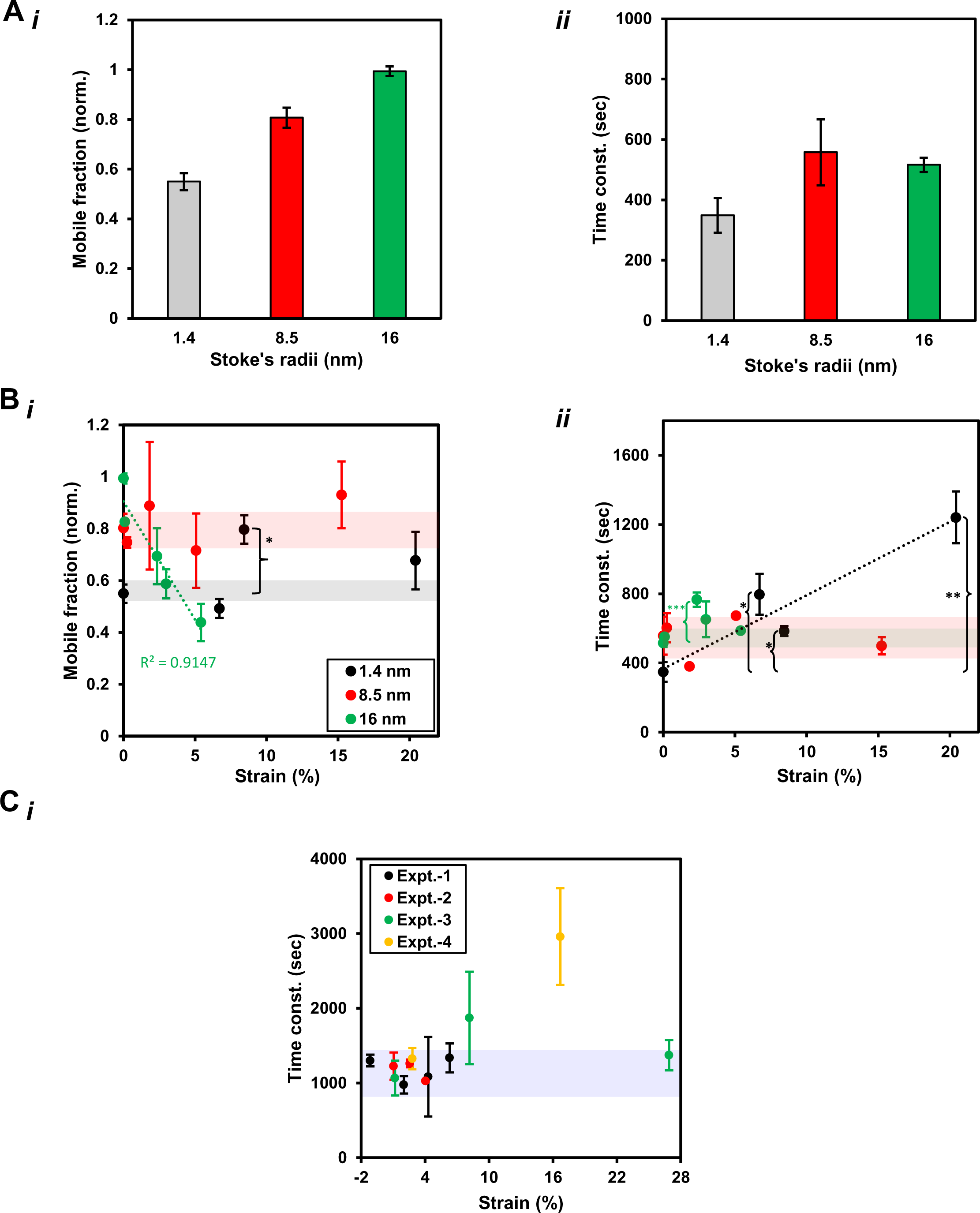
Molecular kinetics of different size fluo. dextran molecules and fluo. BC within mechanically deformed fascicles show mechanical strain dependence. **A.** Mobile fractions and time constants for 1.4 nm, 8.5 nm and 16 nm Stoke’s radii TRITC-dextran molecules in unstrained tendon fascicles. **(*i*)** Mobile fraction values shown an increase with larger size of TRITC-dextran molecules. **(*ii*)** Smaller-size TRITC-dextran molecules (1.4 nm Stoke’s radius) move faster than larger size TRITC-dextran molecules (8.5 nm and 16 nm Stoke’s radii). **B.** Kinetics of TRITC-dextran molecules within a mechanically strained tendon fascicles is determined by the size of dextran molecule and the magnitude of mechanical strain. Fluorescence recovery kinetics of different size TRITC-dextran molecules (indicated by Stoke’s radii) quantified in terms of **(*i*)** Mobile fraction and **(*ii*)** time constants, suggests that smaller size dextran molecules (1.4 nm) are less mobile than larger size ones (8.5 nm and 16 nm) in undeformed fascicles. Mechanical strains (up to ∼8%) do not influence mobile fraction of small size dextran molecules (1.4 nm) while the mobile fraction values of larger size dextrans (16 nm) decreases linearly with increasing mechanical strain values. 1.4 nm dextran molecules tend to move slower at larger strain values within the sample while time constants of larger-size dextran molecules (8.5 nm and 16 nm) do not change significantly with increasing mechanical strain magnitudes. **\* = p < 0.05, ** = p < 0.01, *** = p < 0.001.** **C.** Time constants of fluo. BC molecules are unaffected within physiological strain magnitudes within fascicles (up to ∼8%) while larger strains cause sudden increase in time constant values.

**Fig. S21.**
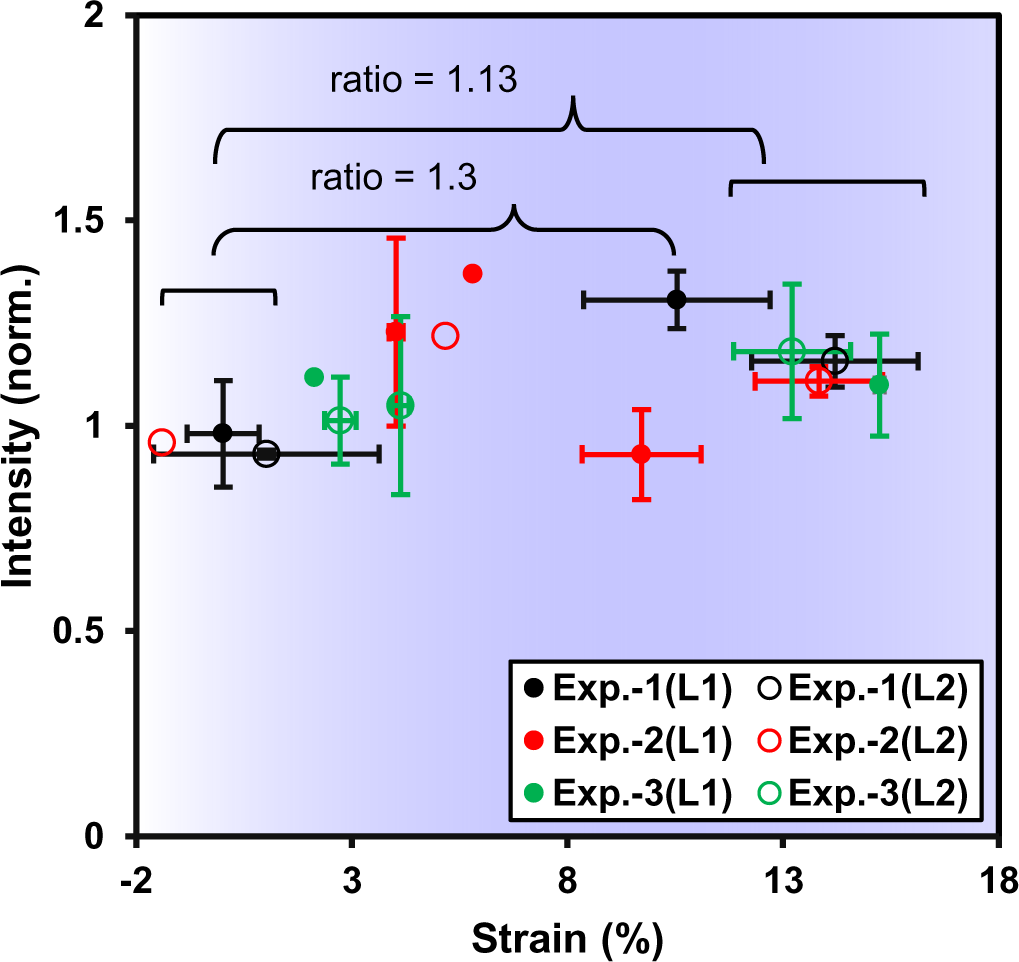
Collagen hybridizing peptide (CHP) shows increased binding towards the high strain regions of a heterogeneously deformed fascicle in the micro-bioreactor indicating increased collagen molecular unfolding at strain > ∼8%. L1, L2 indicate tow different locations of same sample but situated more than at least 1000 um away from each other along the fascicle length. (CHP = -[-glycine(G)–proline(P)–hydroxyproline(O)-]_n_; CF = carboxyfluorescein, n = 6-10)

**Fig. S22.**
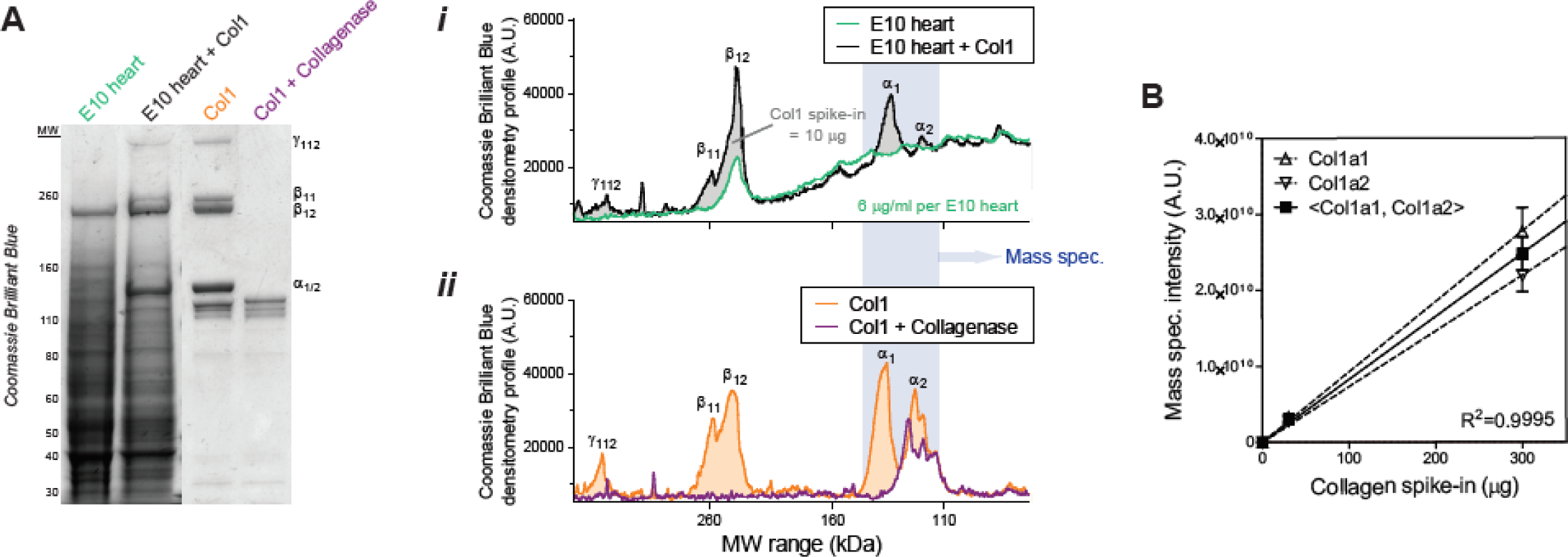
Collagenase (bacterial collagenase) generates degraded fragments of collagen-I present in embryonic chick hearts. **A.** Coomassie Brilliant Blue gel image of E10 heart lysates, E10 hearts with spike-ins of purified rat-tail collagen-I, purified collagen-I only, and collagen-I with bacterial collagenase (n=2 hearts per condition). (***i***-***ii***) Densitometry profiles show collagenase degrades collagen-I and removes all trimer (γ), dimer (β), monomer (α) bands. **B.** MS analysis of gel bands (100–140 kDa) spiked in with known amounts of purified collagen-I provide calibration curves that can be used to approximate total collagen content (in *µ*g’s) in embryonic hearts (per Fig.4A**-ii**).

**Fig. S23.**
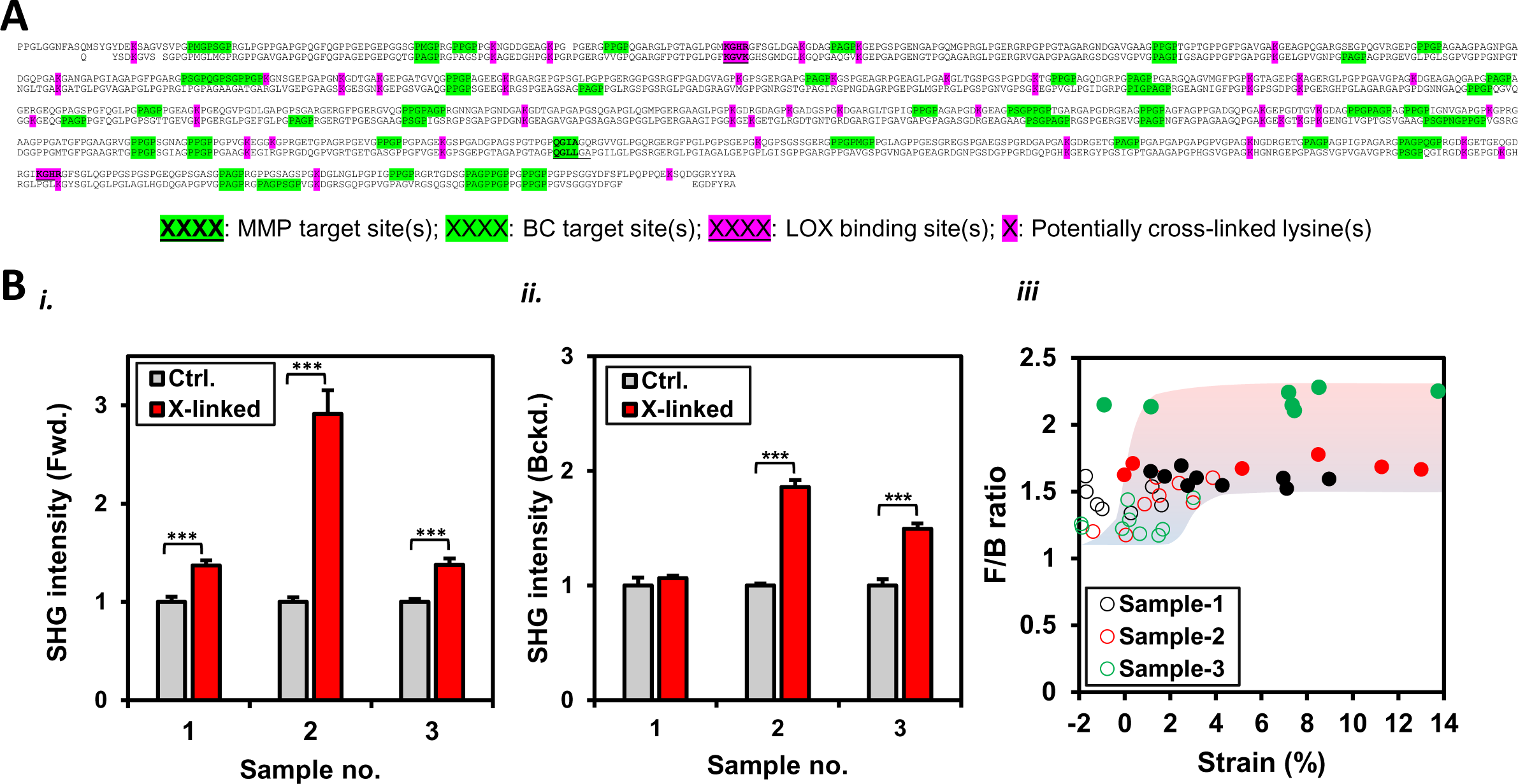
Structural changes in mice tendon fascicle collagen-I following TGM cross-linking. **A.** FASTA sequence(s) of showing cleavage (by MMPs or BCs) and cross-link (by TGM or Lysyl oxidase(s)) susceptible sites within aligned alpha-1 and alpha-2 chains of collagen-I (mice; downloaded from UniProt). **B.** Collagen fibril structural level changes within TGM cross-linked tendon fascicle samples and influence of mechanical strain are evident in SHG signal. **(*i-ii*)** Altered inter-molecular spacing among collagen molecules within collagen fibrils of a strain-free TGM cross-linked fascicle sample compared to control samples (i.e. without TGM cross-links) is reflected by a significant SHG signal in forward (Fwd.) and backward (Bckd.) directions. **(*iii*)** F/B ratio magnitudes within mechanically strained TGM cross-linked samples (represented by filled markers) are significantly different (p < 0.001) from F/B ratio values within strain-free control samples (empty markers). **\* = p < 0.05, ** = p < 0.01, *** = p < 0.001.**

**Fig. S24.**
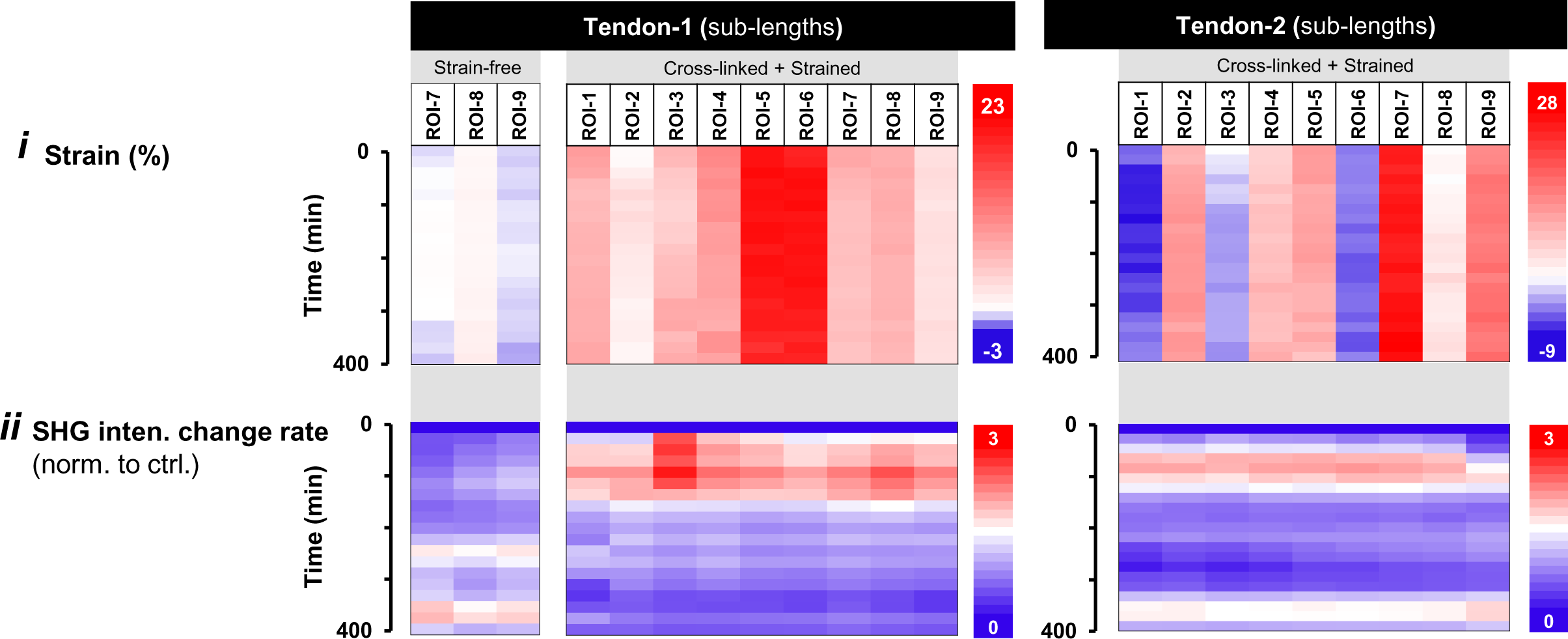
TGM cross-linked fascicle samples under uniaxial deformation and subjected to collagen degradation by BC. **(*i*)** Strain magnitude and **(*ii*)** Collagen degradation rate (for calculations, see Fig. 2C) with time for ROIs within strain-free and strained sub-lengths of tendon fascicles.

**Fig. S25.**
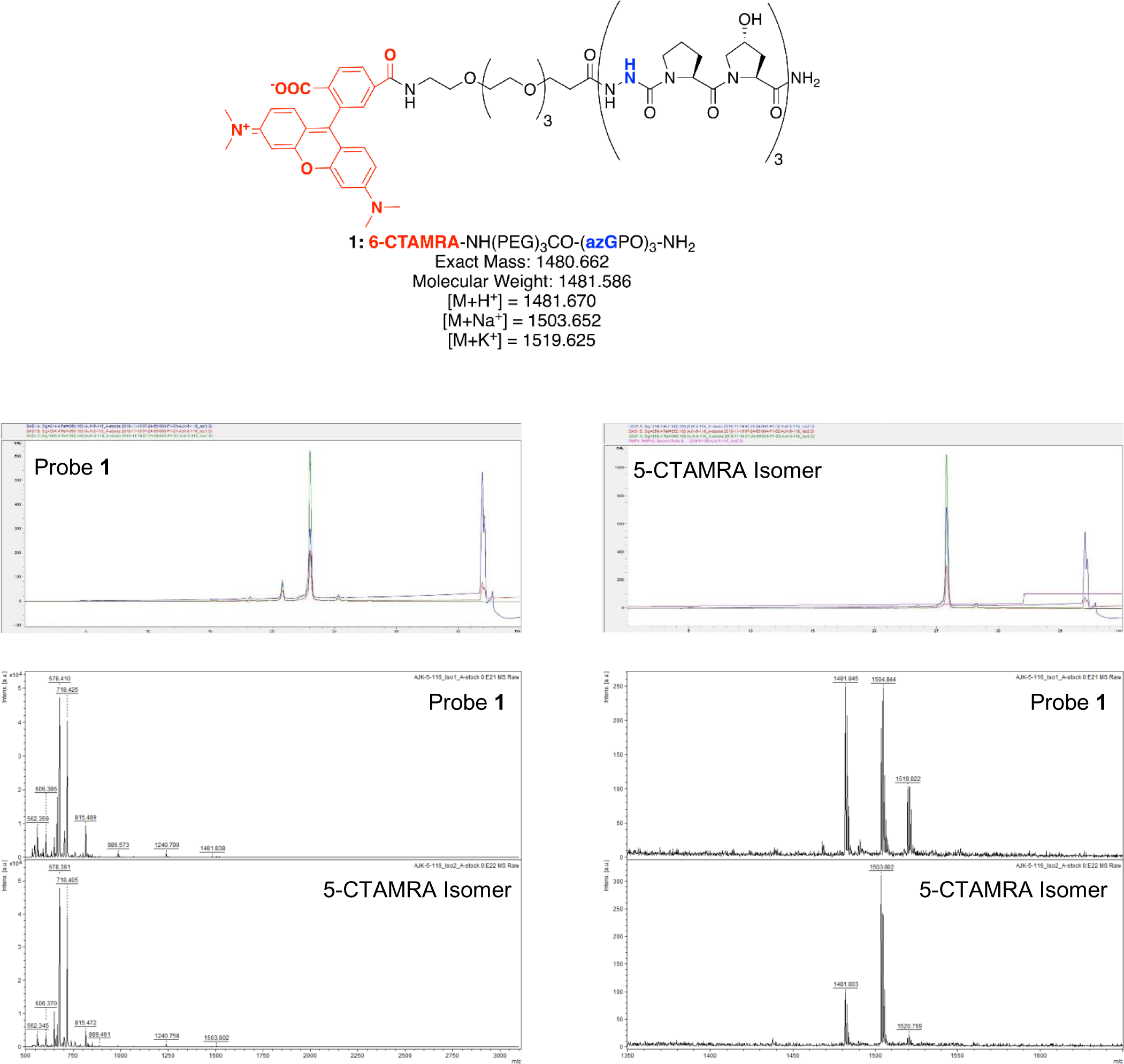
Validation data for aza-peptide probe **1**. **(Top)** Backbone structure and mass data, including predicted ion adducts for MALDI-TOF MS in positive ion mode. **(Middle)** HPLC trace for purified product (10-30% ACN/H_2_O (0.1% TFA), 80 °C, 30 min). **(Bottom)** MALDI-TOF MS spectrum represented in two views: wide mass range (left) and zoomed mass range (right), included to clarify the presence of ion adducts in this data.

**Fig. S26.**
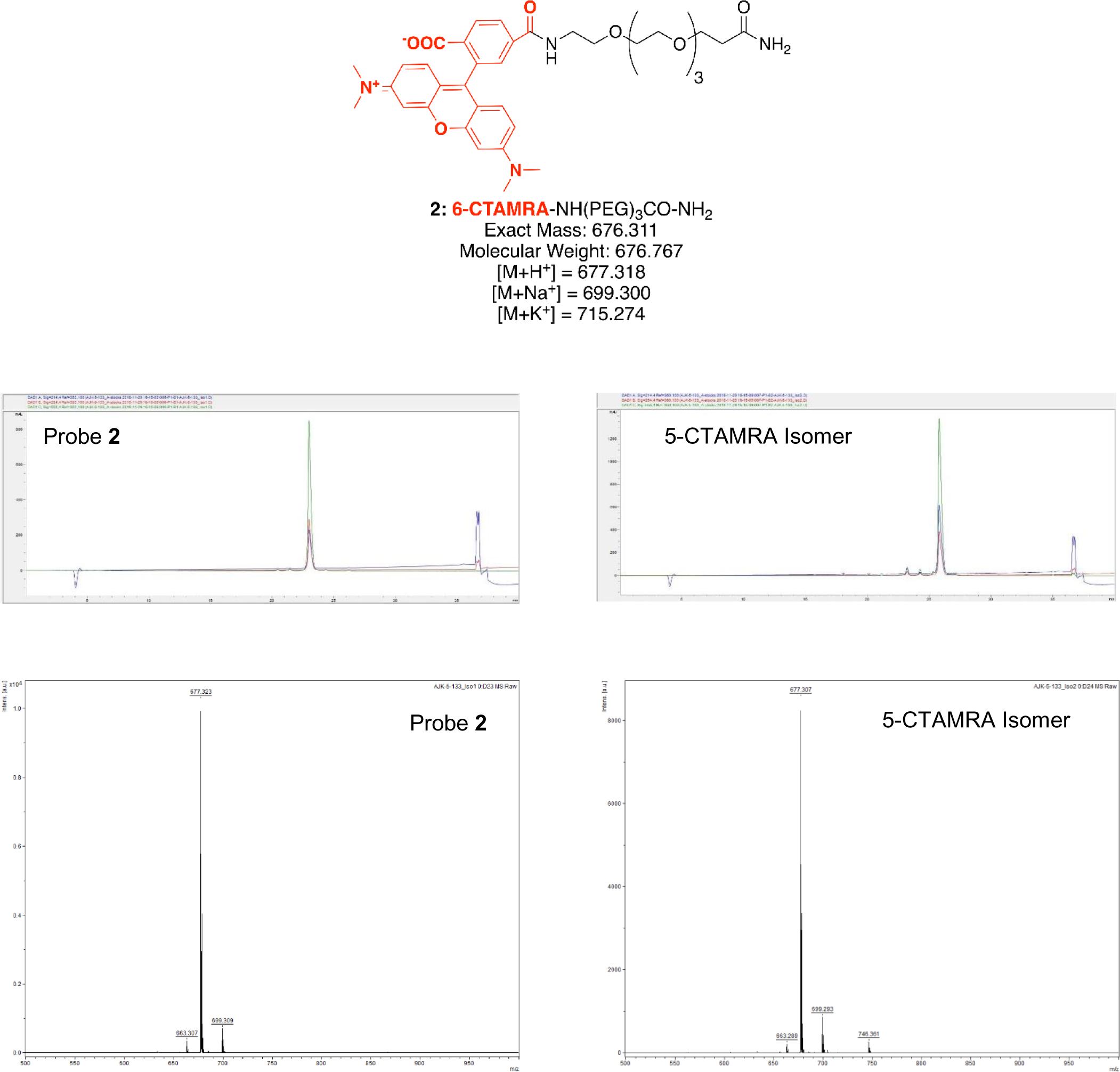
Validation data for 6-CTAMRA-PEG control probe **2**. **(Top)** Backbone structure and mass data, including predicted ion adducts for MALDI-TOF MS in positive ion mode. **(Middle)** HPLC trace for purified product (10-30% ACN/H_2_O (0.1% TFA), 80 °C, 30 min). **(Bottom)** MALDI-TOF MS spectrum.

